# Functional study of *Phaeodactylum tricornutum* Seipin homolog highlights unique features of lipid droplets biogenesis in diatoms

**DOI:** 10.1101/2024.10.11.617903

**Authors:** Damien Le Moigne, Fabien Richard, Catherine Albrieux, Mégane Mahieu, Matteo Arrighi, Grégory Si-Larbi, Pierre-Henri Jouneau, Valérie Gros, Mathilde Louwagie, Sylvaine Roy, Chems Amari, Marta Carletti, Rachel Bonnarde, Yangmin Gong, Yufang Pan, Hanhua Hu, Olivier Bastien, Juliette Jouhet, Alberto Amato, Eric Maréchal, Juliette Salvaing

## Abstract

Diatoms are a major phylum of microalgae, playing crucial ecological roles. They derive from secondary endosymbiosis of a red alga by an unknown heterotrophic eukaryote, leading to a complex intracellular organization. In response to unfavorable conditions (stress), diatoms store oil in lipid droplets (LD), raising interest for applications, in particular biofuels. In spite of numerous investigations aiming to increase their oil content, LD biogenesis mechanisms in these organisms remain poorly understood. In this study, we functionally characterized the homolog of Seipin, a major actor of LD biogenesis, in the diatom *Phaeodactylum tricornutum.* PtSeipin shares conserved structural features with other Seipins, yet presents unique characteristics, that appear common to diatoms and more broadly Stramenopiles. We provide evidence that Stramenopiles Seipins were inherited from the host during secondary endosymbiosis. The localization of PtSeipin highlights that LD biogenesis can arise simultaneously from the endoplasmic reticulum (ER) and the plastid’s most external membrane. Finally, the knock-out of PtSeipin leads to a strong increase of TAG accumulation, a feature that is not observed in other organisms and is greatly enhanced following high light exposure. Our results suggest a redirection of lipid fluxes towards TAG synthesis, reduced TAG recycling or a combination of both.

## Introduction

Microalgae encompass a huge diversity in the tree of life and are found in a great variety of environments. Although they represent only 1% of Earth photosynthetic biomass, microalgae are of crucial ecological importance as they are responsible for around half of total oxygen production (Field et al. 1998; Benoiston et al. 2017). Among them, diatoms, belonging to the Stramenopiles, are the most abundant group. They account for 40% of the net primary production in the oceans, where they play key biogeochemical roles for carbon export, silicon and iron cycles (Armbrust 2009; Benoiston et al. 2017). Diatoms have a complex evolutionary history with two endosymbiosis events leading to the emergence of a “complex plastid” surrounded by four membranes (Guéguen et al. 2021; Le Moigne et al. 2022). In natural conditions, diatoms are constantly exposed to environmental fluctuations, in particular of nutrients availability, light intensity and temperature. They respond to such unfavorable conditions (or stress conditions) by accumulating oil in the form of lipid droplets (LD), a feature that raised interest for potential applications (for review: (Sivakumar et al. 2022; Wang et al. 2024)).

LD are intracellular organelles widespread among eukaryotes. In photosynthetic organisms, and in the absence of sufficient amount of nutrients to ensure cell division, the excess of carbon assimilated through photosynthesis is stored in LD. Their primary function is carbon storage via lipid remodeling, although other functions have been identified (for review (Guéguen et al. 2021; Le Moigne et al. 2022)). The main constituents of LD are triacylglycerols (TAG) and steryl-esters (SE), but they can also contain other hydrophobic molecules such as pigments or waxes. This neutral core is surrounded by a monolayer of glycerolipids. The lipid composition of this monolayer reflects the membrane from which the LD emerged. In the diatom *Phaeodactylum tricornutum,* the core is constituted by TAG and the monolayer contains both the plastidial lipid sulfoquinovosyl-diacylglycerol (SQDG) and the betaine lipid 1(3),2-Diacylglyceryl-3(1)-O-2′-(hydroxymethyl)(N,N,N,-trimethyl)-β- alanine (DGTA) in addition to phosphatidylcholine (PC). This suggests a participation of the plastid’s most external membrane, that is continuous with the endoplasmic reticulum (ER), to LD biogenesis (Lupette et al. 2019). Finally, many proteins are found associated to LD through different hydrophobic motifs, as hairpin structures, amphipathic helixes or lipid anchors (Dhiman et al. 2020). These include structural proteins, which are not conserved among organisms, as well as many proteins that vary depending on the studied condition and/or tissue considered. Among these LD proteins are enzymes involved in lipid synthesis or degradation. The diversity of lipids and proteins that form LD reflects their dynamic nature that responds to constant changes in the environment (for review (Le Moigne et al. 2022; Guzha et al. 2023)).

In most organisms, LD biogenesis begins with TAG accumulation and the formation of TAG lenses between the two leaflets of the ER membrane, continuing with the budding of LD into the cytosol (Walther et al. 2017; Kumari et al. 2023). While budding can occur spontaneously when TAG concentrations are sufficiently high, the Seipin protein can greatly facilitate this process (Santinho et al. 2020).

Seipin has initially been identified for its implication in human Bardinelli-Seip congenital lipodystrophy (Magré et al. 2001) and has also been associated with other metabolic and neurologic diseases (Windpassinger et al. 2004; Ito and Suzuki 2007; Li et al. 2022). Seipin proteins have been mostly studied in animal models, particularly mammals, and yeast. They localize to the ER at discrete sites corresponding to LD biogenesis sites (Szymanski et al. 2007; Fei et al. 2008; Wolinski et al. 2011). They contain two transmembrane domains embedded in the ER surrounding a luminal domain (Lundin et al. 2006; Binns et al. 2010). This luminal domain has been structurally resolved by Cryo-EM in *Drosophilia melanogaster* (Sui et al. 2018), *Homo sapiens* (Yan et al. 2018) and *Saccharomyces cerevisiae* (Klug et al. 2021; Arlt et al. 2022) and shows remarkable structural conservation (reviewed in (Walther et al. 2023)). It allows oligomerization in a ring-like structure. Molecular dynamics simulations indicate important roles of the luminal domain to trap and concentrate TAG (Prasanna et al. 2021; Zoni et al. 2021) while transmembrane domains residues are important for LD maturation and budding (Kim et al. 2022). Interaction with LDAF (Lipid Droplet Associated Factor) protein in human and its homologs in other species are crucial for Seipin nucleation of LD (Eisenberg-Bord et al. 2018; Teixeira et al. 2018; Castro et al. 2019; Chung et al. 2019; Chartschenko et al. 2020). In addition to this function in TAG concentration and LD budding, several publications show that Seipin can participate in the control of glycerolipid synthesis, in particular in phosphatidic acid (PA) metabolism, (Fei et al. 2011; Sim et al. 2012, 2020; Han et al. 2015; Talukder et al. 2015; Wolinski et al. 2015; Pagac et al. 2016; Sołtysik et al. 2020), reviewed in (Salo 2023)).

Interest in plant Seipins has emerged only recently and three Seipin homologs have been identified in the model plant *Arabidopsis thaliana* (Cai et al. 2015). These isoforms all localize to the ER but show different tissue expression patterns (Cai et al. 2015; Taurino et al. 2018). Heterologous expression of the three isoforms in *Nicotiana benthamiana* demonstrated that they promote LD formation, with size variations depending on the isoform (larger LD for AtSeipin1 and smaller LD for the other 2 isoforms). The N-terminal region is crucial for LD size control (Cai et al. 2015). Double and triple knock-outs (KO) of Seipins in *Arabidopsis* lead to accumulation of large LD in seeds and pollen, where budding of LD from the nucleus inner membrane is reported, but without any increase of TAG content (Taurino et al. 2018). Few Seipin partners have thus far been identified in plants: LDIP (LD-Associated Protein [LDAP]-Interacting Protein) seems to be the functional homolog of human LDAF (Pyc et al. 2021) and another partner, VAP27-1 (vesicle-associated membrane protein (VAMP)– associated protein 27-1) was proposed to participate in the stabilization of LD-ER junctions (Greer et al. 2020).

In the diatom *Phaeodactylum tricornutum,* a unique gene encoding a Seipin homolog (PtSeipin) has been identified (Lu et al. 2017) like in animals or yeasts, but the origin of this sequence, its cellular role and its molecular function remain elusive.

As Seipins show little sequence conservation, we investigated the similarities and differences of PtSeipin with other Seipins and performed a comprehensive phylogenetic study showing that Stramenopiles Seipins are closer to those of animals and fungi and were most likely inherited from the host than from the algal symbiont following secondary endosymbiosis. We addressed the localization and function of PtSeipin by a functional genomic approach using KO and overexpression (OE) lines. Our findings confirm that PtSeipin localizes to the endomembrane system with spots at the vicinity of LD. Its localization surrounding the plastid supports the hypothesis that the most external plastid membrane, which is continuous with the ER, is involved in LD biogenesis in diatoms. Eventually, we observed that the morphology of LD in KO lines is strongly altered with the formation of large LD, as has been described in other organisms. Additionally, a strong increase in TAG content was observed in both KO and OE lines. TAG accumulation in KO lines was greatly enhanced by high light exposure. These results show that Seipin general function is conserved in *Phaeodactylum*, but that PtSeipin has developed some unique features and may be an interesting target for biotechnological applications such as biofuel production (Le Moigne et al. 2024).

## Results

### PtSeipin *in silico* analysis

PtSeipin was initially identified by BLAST using AtSeipin2 as a query (Lu et al. 2017). However, the sequence conservation of PtSeipin was low: the alignment of PtSeipin with AtSeipin2 showed only 31% of identity over a short region of 82 amino acids out of 435. An alignment of PtSeipin and *Arabidopsis* Seipins is shown on Supplementary Figure S1. As Seipins from various organisms have conserved structural features (Sui et al. 2018; Yan et al. 2018; Klug et al. 2021; Arlt et al. 2022), we examined the structure predictions for PtSeipin. We first used the Phobius algorithm (https://phobius.sbc.su.se/, (Käll et al. 2007)), predicting the presence of a signal peptide with a posterior label probability of 0.60, as well as two transmembrane domains and an internal hydrophobic domain (Figure 1A). The N- and C-terminal ends were predicted as cytoplasmic, while the central region sits in the lumen of the ER. These predictions are consistent with the ER localization found by Lu and collaborators using LocTree3, as well as with the described structure of other Seipins (Sui et al. 2018; Yan et al. 2018). The predicted structure of PtSeipin using AlphaFold (https://alphafold.ebi.ac.uk/, (Jumper et al. 2021; Varadi et al. 2022)) showed a very high prediction value for the two transmembrane helices as well as most of the luminal domain, while predictions for the N- and C-terminal parts and for two loops in the luminal domain were very low (Figures 1B and C). Superposition of the luminal domain of PtSeipin predicted structure with that of human Seipin (CryoEM resolved structure (Yan et al. 2018)) showed nearly perfect overlay, except for these two loops (Figure 1D, red arrows). Figure 1E shows a cartoon of PtSeipin structural features highlighting conservation with human and fly Seipins (the β-barrel and α-helices 1 to 3) and specific features: the two loops mentioned above are located between the first and second and second and third β-sheets respectively, and each contain predicted additional α- helices, termed α1’ and α2’. Interestingly, the presence of such loops is observed in Seipin proteins from other diatoms and from the eustigmatophytes *Nannochloropsis oceanica* and *Microchloropsis gaditana*, suggesting that this feature is conserved within SAR (Supplementary Figure S2).

**Figure 1:**
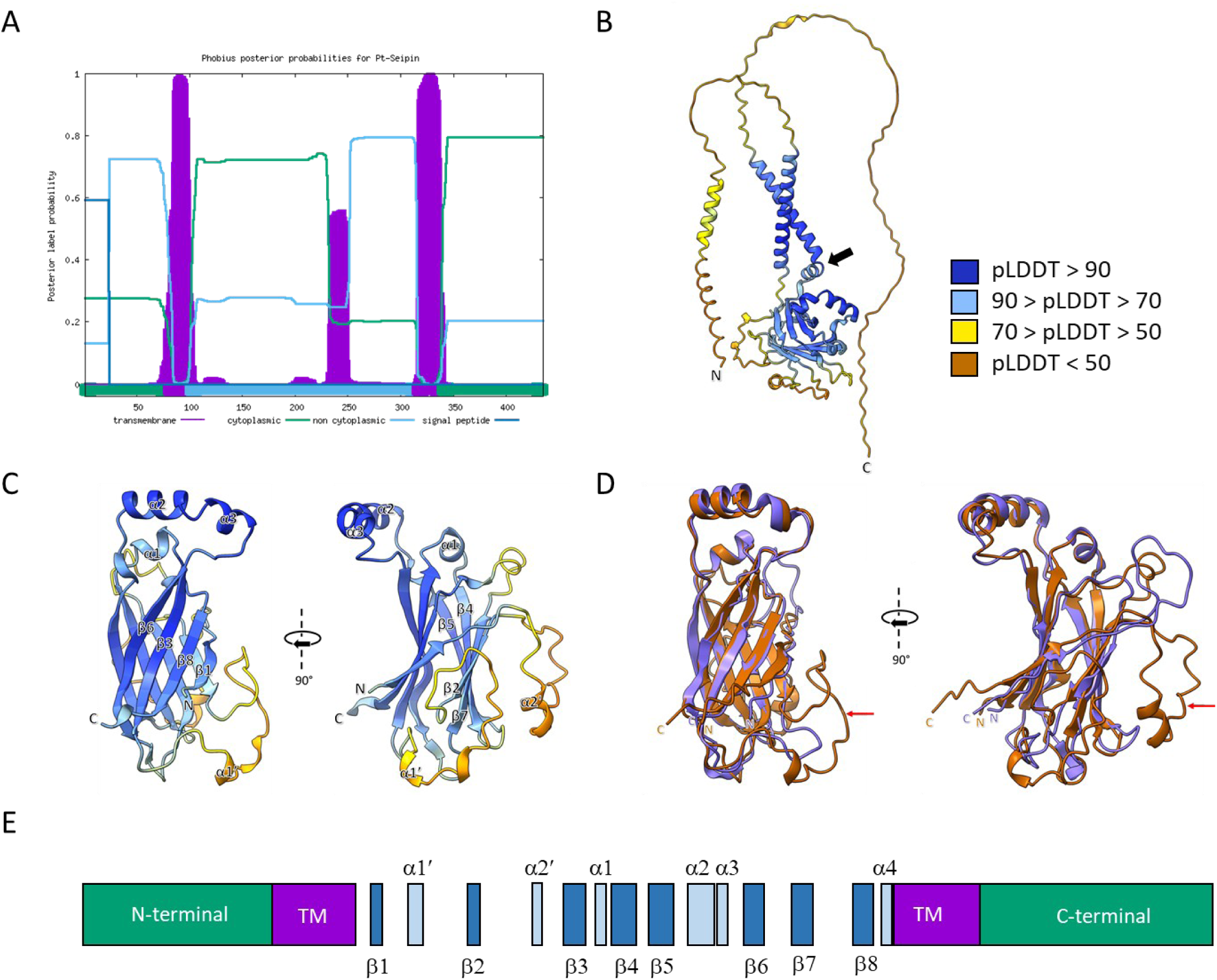
PtSeipin *in silico* analysis. A. Signal peptide and transmembrane domains prediction on PtSeipin complete sequence using Phobius ( https://phobius.sbc.su.se/, (Käll et al., 2007)). Predicted transmembrane domains are represented in purple; green and light blue coloring of the bottom line indicate respectively that a domain is cytoplasmic or non cytoplasmic; signal peptide prediction is represented with a dark blue line. B. PtSeipin structure prediction by AlphaFold (https://alphafold.ebi.ac.uk/entry/B7G3W8). The color code represent the per residue prediction model confidence (pLDDT) according to (Jumper et al., 2021). Dark blue: very high confidence (pLDDT > 90); light blue: high confidence (90 > pLDDT > 70); yellow: low confidence (70 > pLDDT > 50); orange/brown: very low (pLDDT < 50). The black arrow highlights the presence of a “kinked helix” just before the C-terminal transmembrane helix, an important conserved feature (Arlt et al., 2022). C. Zoom on the luminal domain. The α-helices (α1-3) and β-sheets(β1-8) have been labelled to match labelling of published CryoEM Seipin structures from human and flies (Sui et al., 2018; Yan et al., 2018). Additional α-helices that appear on the AlphaFold prediction have been labelled α1’ and α2’. α4 helix corresponds to the “kinked helix” (Arlt et al., 2022). D. Superposition of PtSeipin predicted structure (brown) and human Seipin resolved CryoEM structure (purple). Red arrows highlight extra loops in PtSeipin structure. E. General diagram of PtSeipin organization. The size and spacing of the different features respect their positions in PtSeipin sequence. This diagram highlights the position of the two loops containing the predicted α1’ and α2’ helices, respectively between β1 and β2 and between β2 and β3. Green: N- and C-terminal domains; purple: transmembrane domains (TM); dark blue: β-sheets; light blue: α-helices.

Studies of Seipins from human, fly and yeast have highlighted the importance of several subdomains and residues within the luminal domain. To investigate the conservation of these subdomains in *Phaeodactylum* and more generally throughout the tree of life, we searched for motifs using the MEME tool (https://meme-suite.org/meme/tools/meme, (Bailey and Elkan 1994)) in a subset of sequences from 48 species representing the diversity of Opistokontha (animals and fungi), Archaeplastida (green algae and land plants) and SAR (Oomycota, Eustigmatophytes and Diatoms) (Supplementary Table 1). The complete results are shown in Supplementary Figure S3. Three motifs (A, C and E), corresponding respectively to α-helices 1 to 3 and surrounding regions, appear conserved in nearly all sequences (Supplementary Figure S4). α-helices 2 and 3 correspond to the hydrophobic helices (Sui et al. 2018) or membrane-bound helices (Yan et al. 2018), which are involved in oligomerisation and interact with lipids in the ER membrane. Several amino-acids from α−helix 1 are also involved in the oligomeric interface according to Sui and collaborators (Sui et al. 2018). Of these, only one Serine residue (S117 in the human sequence, position 6 in the motif A) appears to be conserved.

### Phylogeny of Seipins

Seipins that have been studied so far belong to either the Opistokontha (mammals, flies, yeast) or the Archaeplastida (*Arabidopsis thaliana)*. In their initial study of PtSeipin, (Lu et al. 2017) provided an incomplete phylogenetic tree with species belonging to a restricted number of phyla. We aimed to provide a comprehensive phylogeny of Seipins using the dataset used for the motif analysis. Information gained from this motif analysis (presence/absence of motifs, relative position in the sequence, distance between adjacent motifs) was coded as described in Supplementary Table 2 and included in the phylogenetic analysis as described in the material and methods section. Figure 2 shows the phylogenetic tree obtained. Seipins from Archaeplastida (green algae and land plants) and Opistokontha (animals and fungi) form two distinct clades. Only Seipins from nematods appear to have significantly diverged and do not cluster with other animal sequences. Interestingly, Seipin from all SAR members included in the study (4 diatoms, 2 eustigmatophytes and 2 oomycetes) cluster together, and group closer to Opisthokonta than to Archaeplastida. This suggests that Seipin from SAR were inherited from the host rather than from the algal symbiont following secondary endosymbiosis events.

**Figure 2:**
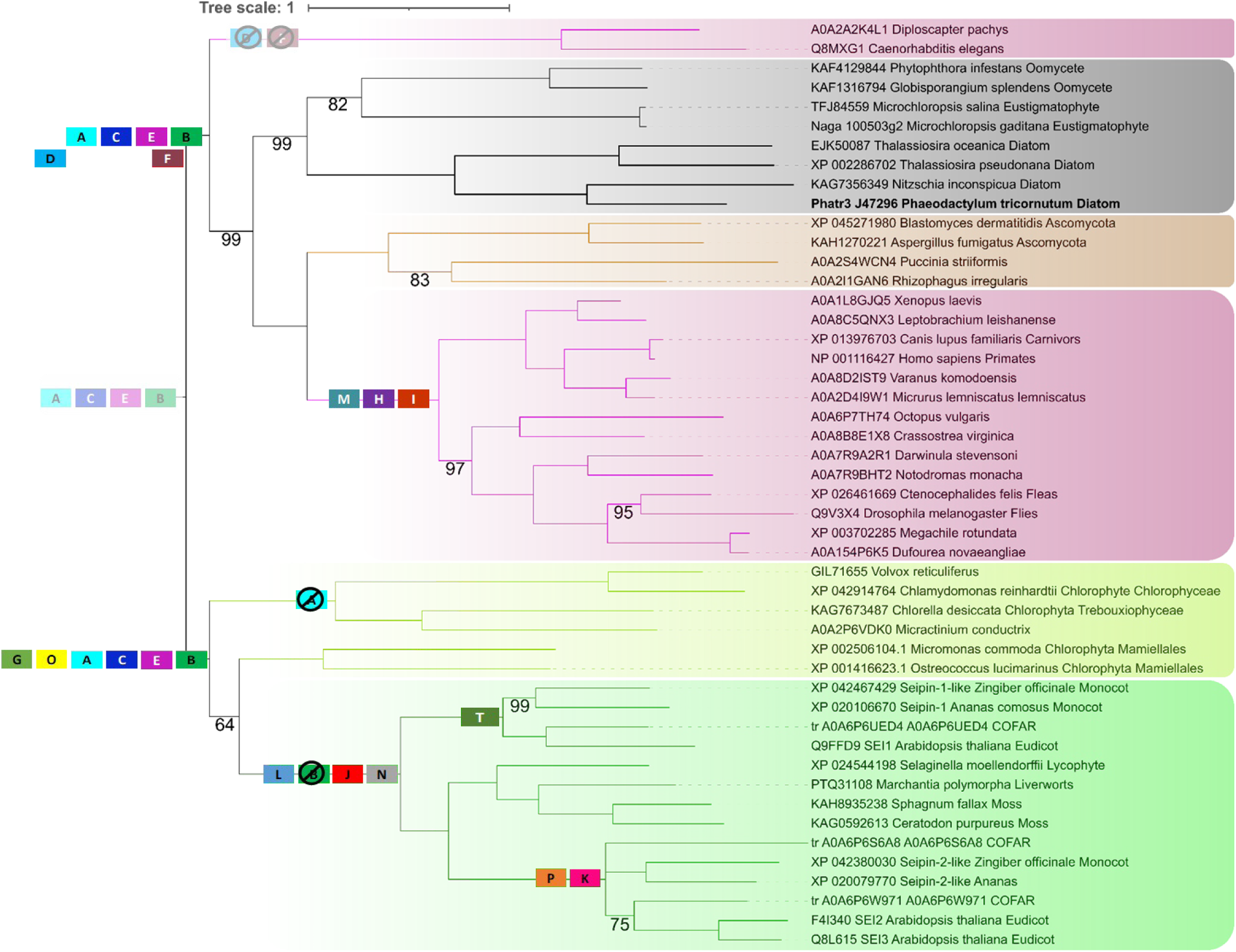
Phylogeny of Seipin proteins. Midpoint rooted tree of Seipins. Sequence identification and phylogenetic analysis are described in the material and methods. Only posterior probabilities different from 100 are shown. Sequences are colored according to the great groups to which they belong: Archaeplastida (Land Plants and Green algae, in green and yellow green respectively), Animals (in pink), Fungi (in brown) and SAR (in grey). Diatom sequences are highlighted with larger lines and *Phaeodactylum tricornutum* is in bold font. Presumed domains acquisition and loss based on the ancestral states analysis are shown on the picture. Domains A, C, E and B that are common to both the SAR/Opistokontha and Archaeplastida are shown as probable in the common ancestor (shaded colors). Domains D and F are found both in the ancestor of SAR and in that of Opistokontha but are not detected in nematoda sequences, which appear to be very divergent and emerge at the basis of the SAR/Opistokontha clade. As it is more parsimonious to suppose that they were present in the common ancestor and subsequently lost in nematode, they were indicated at this position but placed below the other domains.

Based on this tree, we analyzed the ancestral states for the two major clades (SAR/Opistokontha and Archaeplastida), and of subdivisions of these groups (SAR, SAR/Diatoms, Opistokontha, Opistokontha/Fungi, Opistokontha/Animals on the one hand and Green Algae, Land Plants, Land Plants/Seipin1 and Land Plants(Seipin2/3) on the other (Supplementary Tables 3 and 4). Four motifs are found to be ancestral to both major clades (Supplementary Table 3 and Figure 2), suggesting that they were present in the most ancestral form of Seipin. Motifs A, C and E have already been presented above; the fourth one, motif B, is located just upstream of the C-terminal transmembrane domain (Supplementary Figures S1, and S4B), and includes the so-called “kinked helix” (α4 on Figure 1E). Functionally important conformational changes in this region have been highlighted in yeast (Arlt et al. 2022). Motif B has been lost in several species and seems to have evolved into motif J in Land Plants, as these two motifs are detected at the same position in some species (Supplementary Table 2). Interestingly, both motifs B and J can be found in diatoms (B in *P. tricornutum* and *Thalassiosira pseudonana* and J in *Thalassiosira oceanica* and *Nitzschia inconspicua*), suggesting that the evolutionary history of Seipins might be more complex and that gene recombinations between the red alga and the host of the secondary endosymbiosis have probably occurred. Indeed, other domains found in Archaeplastida but not in Opistokontha can be identified in SAR species, such as the N domain in *T. pseudonana,* the O and L domains in *P. tricornutum* (with a non significant p-value), and the O domain in *Microchloropsis gaditana* and *Microchloropsis salina* (significant p-value but modified position) (Supplementary Table 2).

### PtSeipin expression in yeast

To investigate whether PtSeipin function was similar to that of yeast Seipin, we expressed PtSeipin in the *S. cerevisiae* Seipin (FLD1/SEI1) deletion mutant strain *ylr404w*Δ (Szymanski et al. 2007; Fei et al. 2008; Cartwright et al. 2015). We observed the complemented strain by confocal laser scanning microscopy. As controls, we included the yeast wild type (WT) strain, and the *ylr404w*Δ strain complemented by expression of the yeast SEI1 protein. Expression of the different Seipin proteins was further confirmed by Western Blot using an anti-HA antibody (Supplementary Figure S5B). Expression of PtSeipin appeared to be mildly deleterious for the yeast, as the *ylr404w*Δ*/PtSeipin* strain showed altered growth compared to all other strains (Supplementary Figure S5A). As described previously (Szymanski et al. 2007; Fei et al. 2008), the mutant yeast strain showed fewer larger LD when compared to the WT, and expression of SEI1 in the mutant restored the size and number of LD (Figure 3, (Cai et al. 2015)). Interestingly, PtSeipin expression restored the size and number of LD in a similar way to SEI1 (Figure 3), confirming that its general function in LD biogenesis is conserved.

**Figure 3:**
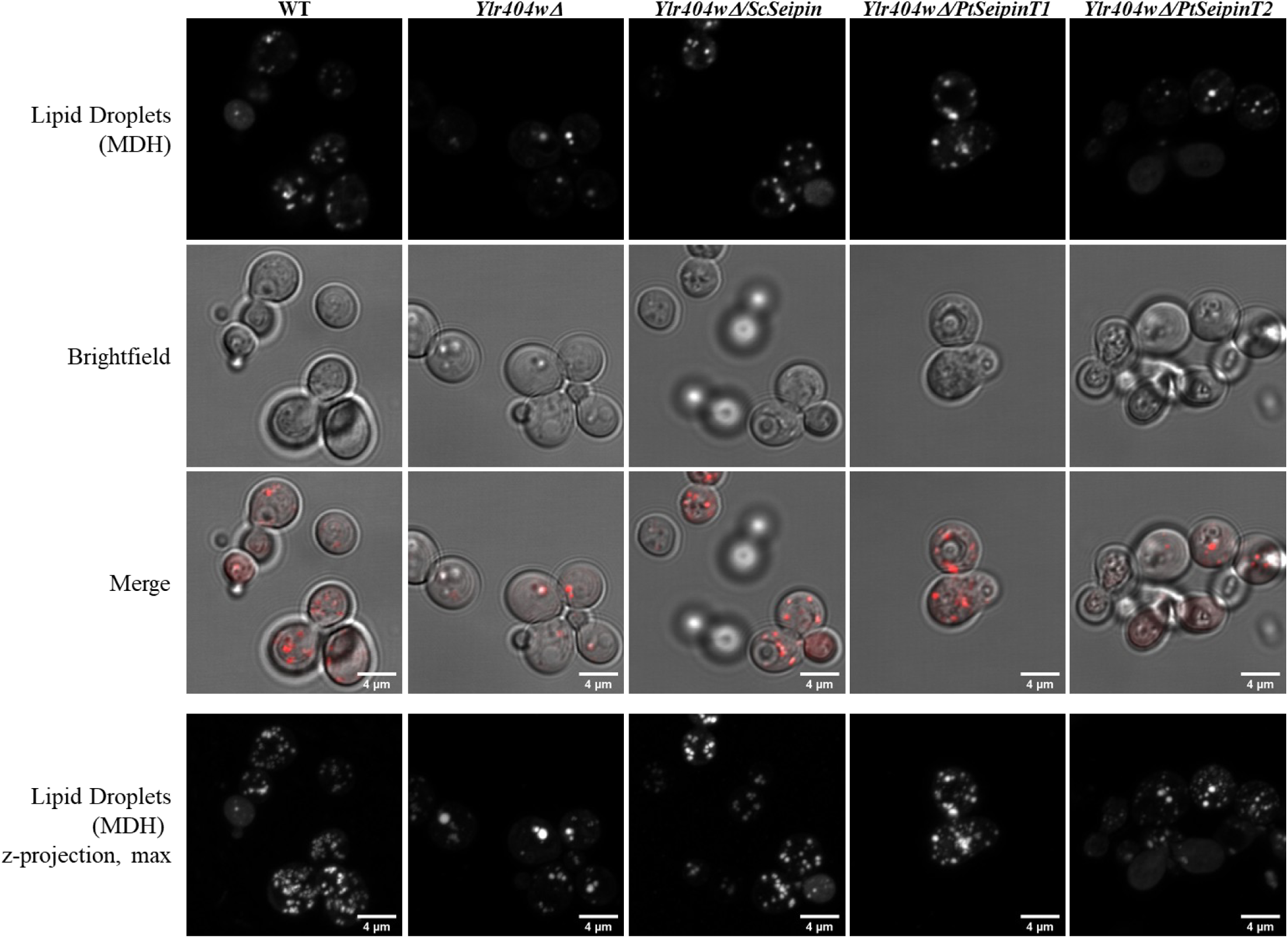
Complementation of Δ*Fld1 (Ylr404wΔ)* yeasts. Confocal imaging of Lipid Droplets stained with Nile Red in yeasts. Wild-type (WT) and Seipin deficient (*ylr404wΔ*) strains were transformed with the empty plasmid YCplac33. *Ylr404wΔ* was either complemented by yeast Seipin (*Ylr404wΔ/ScSeipin*) or by *Phaeodactylum tricornutum* Seipin (*Ylr404wΔ/PtSeipin*). As the latter led to growth delay, two different time points are shown. *Ylr404wΔ/PtSeipin*T1 cells were harvested at the same time but at a lower OD_600_ while *Ylr404wΔ/PtSeipin*T2 were harvested at a later time but at similar OD_600_. The three upper lines show one z-scan while the lowest line show a z-projection of the maximal intensity of lipid droplets fluorescence. Scale bar: 4μm.

### PtSeipin localization

In yeast and animal cells, Seipin proteins localize to discrete areas of the endoplasmic reticulum (ER), forming LD nucleation sites (Szymanski et al. 2007; Fei et al. 2008; Salo et al. 2016; Wang et al. 2016). Two major lines of evidence suggest that LD biogenesis in *P. tricornutum* could also occur at the outer membrane of the plastid, also called Epiplastidial membrane (EpM) or chloroplastic endoplasmic reticulum membrane (cERM); first, many contact sites are observed between the LD and the EpM; second, the lipid monolayer surrounding LD contains lipids with plastidial origin: the sulfolipid SQDG, and specific species of the betain lipid DGTA containing fatty acids that are produced in the plastid (Lupette et al. 2019; Guéguen et al. 2021). In order to determine the localization of PtSeipin in *Phaeodactylum*, we expressed fusion proteins with eGFP in several independent lines, and lines with different expression levels were obtained (Supplementary Figure 6A). The overexpressing lines were observed using confocal laser scanning microscopy. Figure 4 shows representative images of two independent lines expressing PtSeipin-GFP in exponential growth condition, with additional staining of LD with monodansylpentane (MDH). Representative images of independently obtained PtSeipin-GFP overexpressing lines (See methods and Supplementary methods 1) in exponential phase and in starvation conditions are shown on Supplementary Figure S6B and Supplementary Figure S7 respectively. Fluorescence was always observed in the plastid but comparison of higher (PtSeipin GFPa on Figure 4, PtSeipin-GFP8 on Supplementary Figure S6) and lower expressing lines (PtSeipin GFPb on Figure 4, PtSeipin-GFP5 and 9 on Supplementary Figure S6) highlights that it is likely due to chlorophyll autofluorescence that is detected in the GFP channel. In cells grown in exponential phase, where only small LD are observed, brighter foci are detected in close vicinity to these LD (Figure 4 and Supplementary Figure S6B), while in starved cells, where LD are much bigger, the GFP signal is detected in large areas around the LD (Supplementary Figure 7A). Weaker GFP signal was also visible in the ER and all around the plastid (Figure 4 and Supplementary Figure 7A), as illustrated on a 3D reconstruction (Supplementary Figure 7B), but it was barely or not detectable in the lower expressing lines (PtSeipin-GFP5 and 9, Supplementary Figure S6).Interestingly, we often observed that LD were associated with more than one foci of PtSeipin-GFP, with part of the foci located at the plastid surface and others outside of the plastid (Figure 4 close-ups), suggesting that both ER and cERM may contribute together to LD biogenesis.

**Figure 4:**
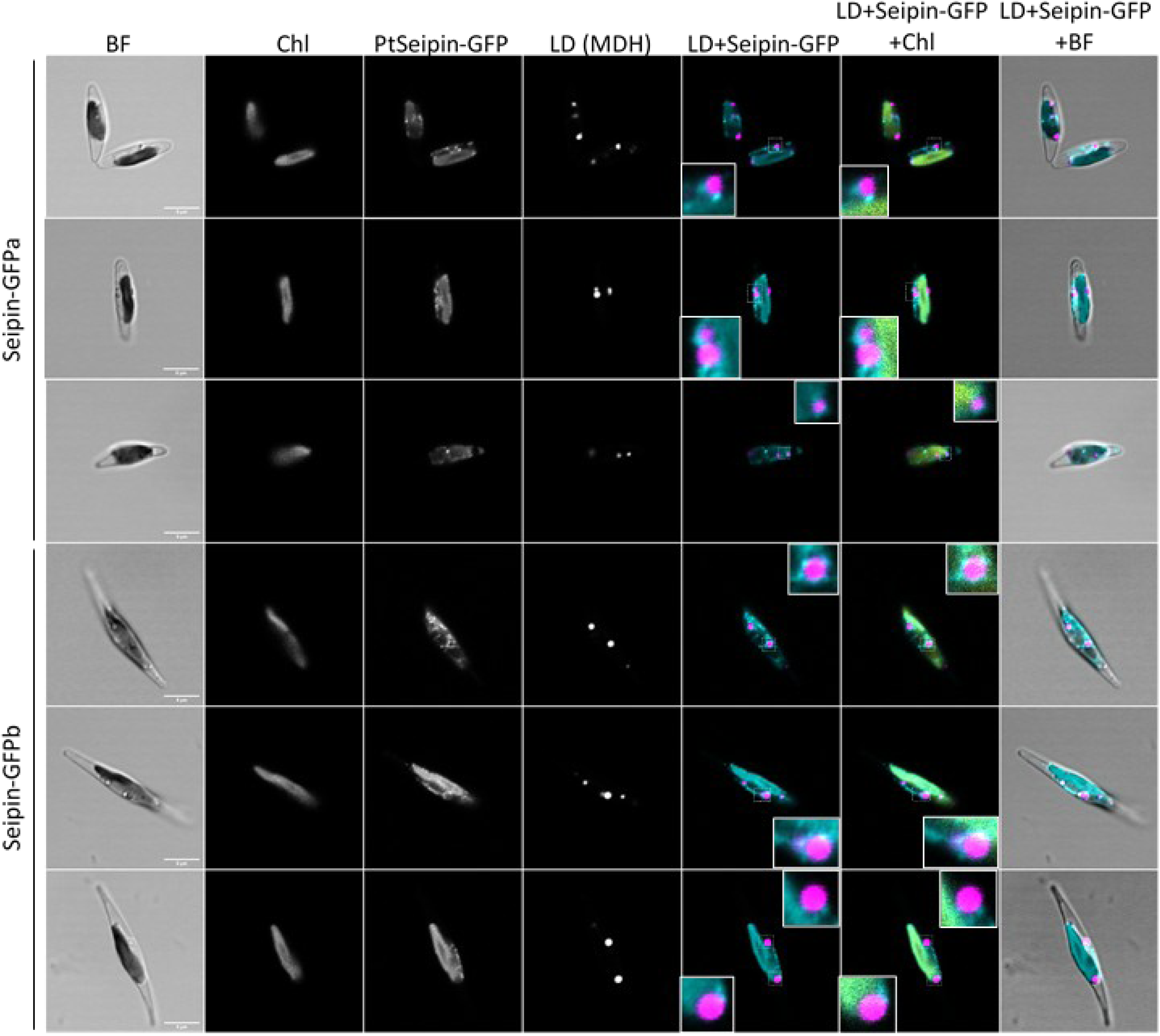
Laser Scanning Confocal Microscopy (LSCM) observation of the *in vivo* localization of PtSeipin *in Phaeodactylum tricornutum* in control culture conditions in exponential growth. Representative images of two PtSeipin-GFP (a and b) cell lines. From left to right: pseudo-brightfield (BF: greyscale); Chlorophyll autofluorescence (Chl: greyscale), PtSeipin-GFP signal (greyscale), LD stained with MDH (greyscale), overlay of LD and GFP signals (respectively magenta and cyan), overlay of LD, GFP and chlorophyll signals (respectively magenta, cyan and yellow), overlay of pseudo-bright field with LD and GFP signals (respectively grey, magenta and cyan). Scale bar: 5 µm. Close-ups on some LD showing contacts with PtSeipin signal are shown.

### Generation of PtSeipin Knock-Out mutants

In order to characterize the function of PtSeipin in *P. tricornutum*, we used three PtSeipin-eGFP overexpression lines with different expression levels (PtSeipin-GFP5, 8 and 9, *cf.* Supplementary Figure 6) and generated Knock-Out (KO) lines, using the CRISPR-Cas9 system previously implemented in *P. tricornutum* (Nymark et al. 2016; Allorent et al. 2018). As the use of the CRISPR-Cas9 system can lead to off-target effects, two guide RNAs targeting the *PtSeipin* gene upstream of the region coding for the first transmembrane domain were chosen (Supplementary Figure 8). For each of these guides, three independent mutant lines were selected to rule out random effects of different genome insertions linked to the biolistic genetic transformation. The details of the mutations are shown on Supplementary Figure 8. For the sake of clarity, we only show results for two of these mutants, one for each guide, in the main figures, while results obtained with other mutants are included as supplementary information. We only considered results that were consistent among all the mutant lines across all experimental replicates. Similarly, we selected two of the PtSeipin-eGFP overexpressing lines, the lowest and the highest, to be presented in the main results section.

### Assessment of physiological parameters in PtSeipin mutants

We first examined whether alterations in PtSeipin content affected cell physiology. For this, we followed the growth of both KO and OE lines over a period of 8 days (Figure 5A), and measured Fv/Fm ratio for these lines as an estimate of photosynthesis efficiency (Figure 5B), and Nile Red fluorescence as a marker of neutral lipids accumulation (Figure 5C). We only detected a mild alteration of the growth of the PtSeipin-GFP OE8 line at later time points (Figure 5A right), which is consistent with results obtained by Lu and collaborators (Lu et al. 2017) with PtSeipin-cMyc lines. The Fv/Fm values in the PtSeipin-GFP OE8 line also appeared slightly decreased (Figure 5B), but even though the difference with the WT is statistically significant, the variation observed is too small to reflect a real impact on photosynthetic activity. However, by contrast with past reports, Nile Red fluorescence in both OE lines was significantly lower than in the WT at day 1, and only showed a mild non statistically significant increase at day 8 (Figure 5C). PtSeipin KO also did not alter growth: a small growth delay was observed in the ΔSeipin1.3 line (Figure 5A left) but such delay was not consistently observed across all KO lines nor across all experiments (Supplementary Figure 9). A small but significant decrease of the Fv/Fm ratio was also observed in both KO lines at all time-points(Figure 5B). As for PtSeipin-GFP OE8, the difference between ΔSeipin8.3 and WT was very small, but the stronger difference observed for this line at day 8, and for the ΔSeipin1.3 line at all considered time points suggest that photosynthetic activity was slightly altered in the Seipin KO, especially when cells had entered the stationary phase. Finally, a small but significant increase of Nile Red fluorescence was also observed in the PtSeipin KO lines, at day 5 and 8 for ΔSeipin8.3 and at day 7 and 8 for ΔSeipin1.3 (Figure 5C). These results suggest that PtSeipin KO but not PtSeipin OE lines have a higher neutral lipids content. This experiment was used to select reference time points used for all subsequent experiments: 4 days, corresponding to the end of the exponential phase, and 8 days, corresponding to the stationary phase and to the first signs of nutrient deficiency. Data corresponding to day 4 are shown in the main figures, while data obtained at day 8 are shown as supplementary figures.

**Figure 5:**
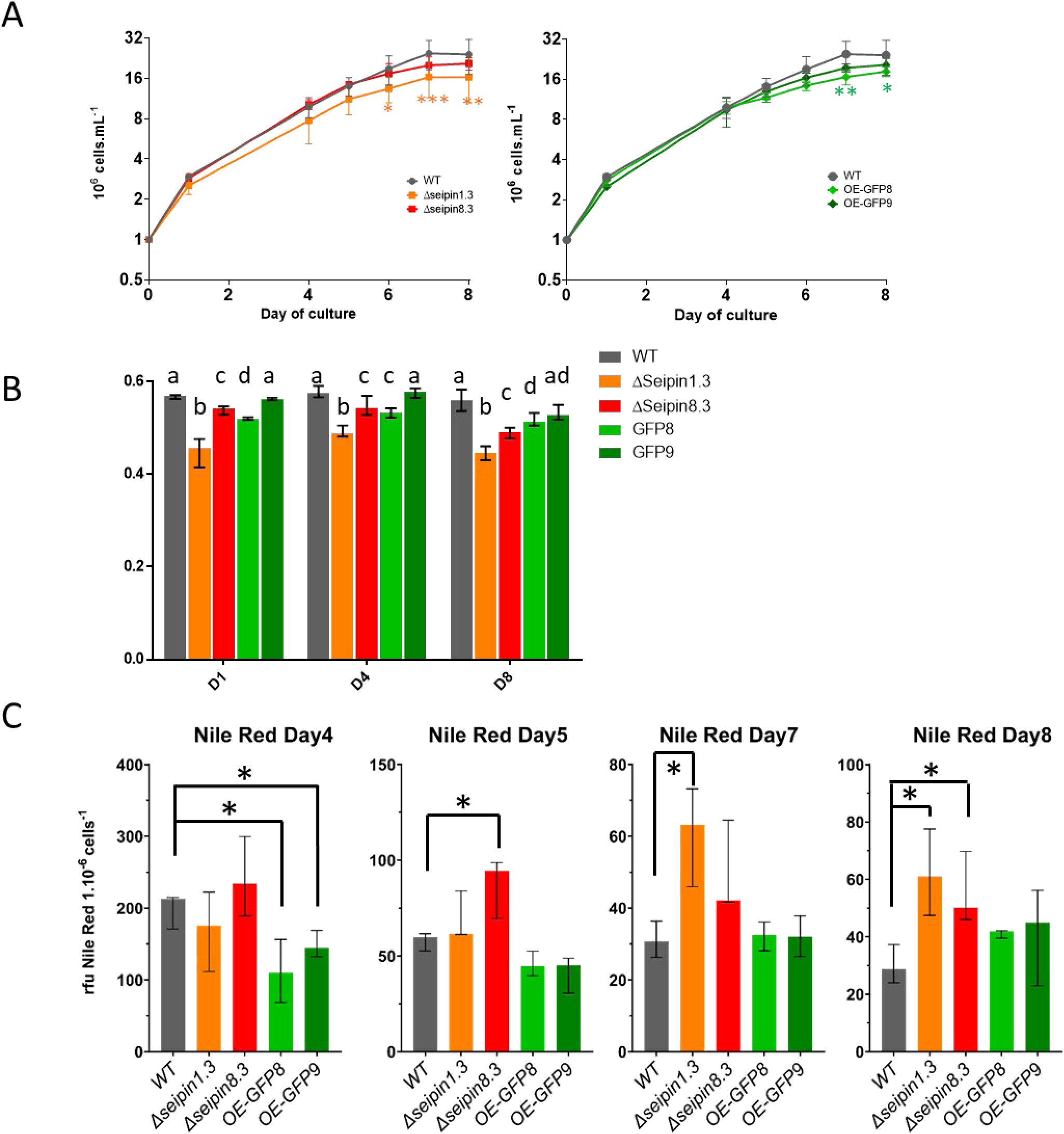
Cell physiology parameters in PtSeipin mutants under control culture conditions. A) Growth curves of WT (grey), PtSeipin KO (orange and red, left panel) and PtSeipin-GFP overexpressing lines (light and dark green, right panel). Cell concentrations (in million cells.mL^-1^) were calculated based on absorbance at 730 nm, which was measured during 8 days. Median, min and max of triplicates are shown. Statistically significant differences between mutants and WT were evaluated by 2- way ANOVA with Dunett’s multiple comparison test and are indicated as follows: *: p-value<0.05; **: p-value<0.01; *** p-value <0.001. B) Photosynthesis efficiency of WT (grey), PtSeipin KO (orange and red) and PtSeipin-GFP overexpressing lines (light and dark green) was evaluated by measures of Fv/Fm at three points of the curve. Day 1: exponential phase; day 4: end of exponential phase; day 8: stationary phase. Statistically significant differences between the different cell lines were evaluated using multiple t-tests. Different letters indicate significant differences. C) Neutral lipids accumulation during the stationary phase was evaluated by fluorescence intensity measures following Nile Red staining. WT is in grey, PtSeipin KO lines in orange and red and PtSeipin-GFP overexpressing lines in light and dark green. Statistically significant differences between mutants and WT were evaluated using multiple t-tests and are indicated as follows: *: p-value<0.05.

### The number and size of LD are altered in the PtSeipin KO

In many species, Seipin depletion leads to changes in the morphology of LD (Szymanski et al. 2007; Fei et al. 2008; Wang et al. 2016; Taurino et al. 2018). We thus wondered whether this was also the case in PtSeipin mutants and whether changes in Nile Red fluorescence reflected changes in LD size and number. PtSeipin KO and OE lines were observed after 4 and 8 days of culture in replete conditions by confocal microscopy (Figure 6A and Supplementary Figure S10A), and the number and size of LD were measured using ImageJ (Figure 6B and C and Supplementary Figure S10B and C). At both time points and in all of the PtSeipin KO lines, we observed that the cells had a tendency to contain less LD than the WT (Figure 6B and Supplementary Figure S10B), and that LD size was highly variable and could reach several microns in diameter (Figure 6C and Supplementary Figure S10C), which is normally only observed after severe nutrient starvation (Jaussaud et al. 2020). In addition, small alterations of cell shape were observed in most of the mutant lines, as cells appeared shorter and wider (Figure 6A). By contrast, the PtSeipin OE lines showed a tendency to have more LD and the size of their LD was similar to that in the WT, or even slightly smaller (Figure 6B and C and Supplementary Figure S10B and C).

**Figure 6:**
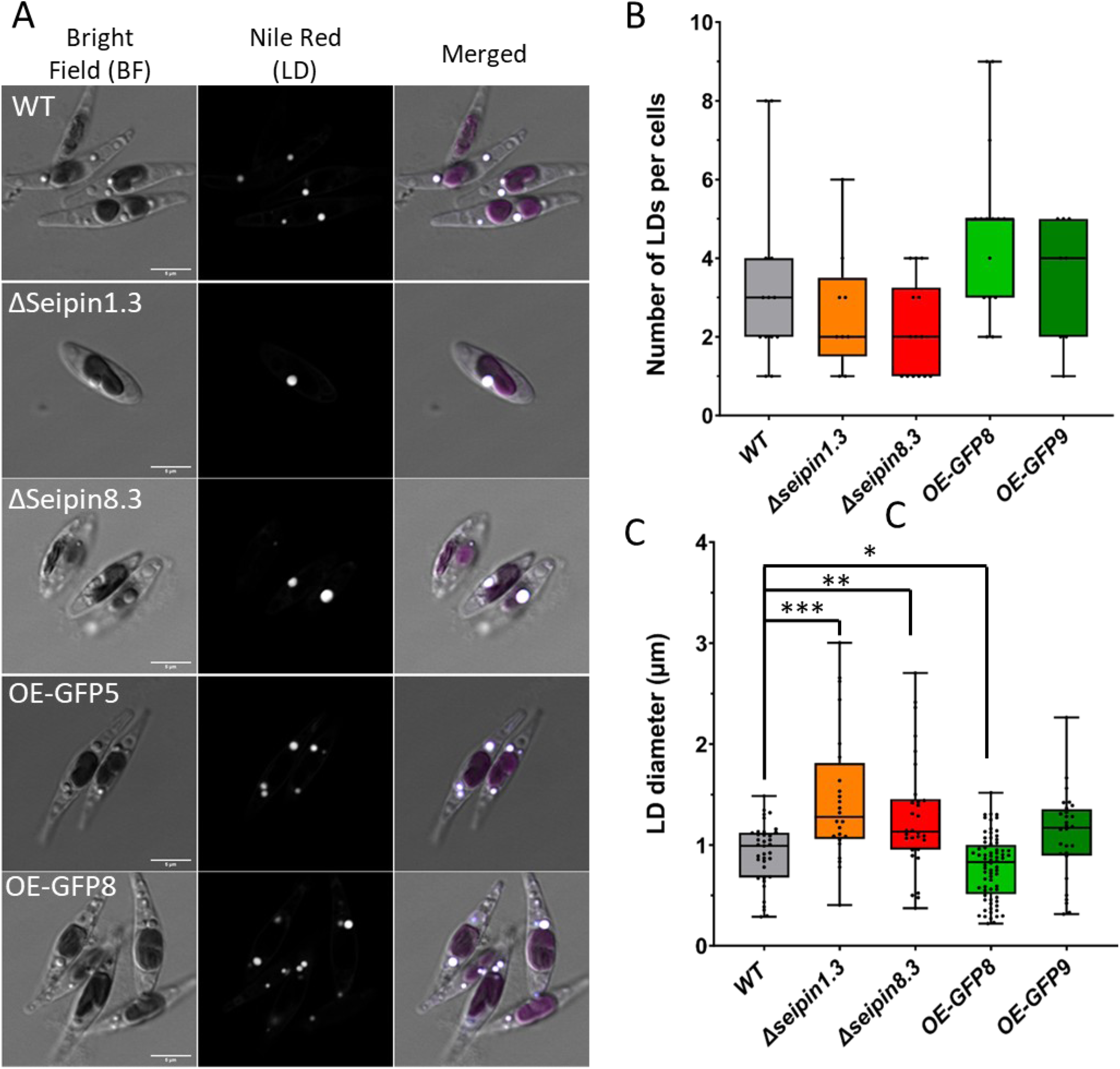
Observation and quantification of LD in PtSeipin mutants after 4 days of culture in control (CT) culture conditions. A) Laser Scanning Confocal Microscopy (LSCM) images of WT, PtSeipin KO (ΔSeipin1.3 and ΔSeipin8.3) and PtSeipin-GFP overexpressing lines (OE-GFP5 and OE-GFP8) after staining with Nile Red. From left to right: pseudo-bright field (grey), Nile Red fluorescence (greyscale) and overlay of pseudo-bright field with Nile Red and chlorophyll signals (respectively grey, cyan and magenta). Scale bar: 5 µm. B) Quantification of the number of LD per cell in all the cell lines: WT (grey), PtSeipin KO (orange and red) and PtSeipin-GFP overexpressing lines (light and dark green). Results are presented as boxplots and each dot corresponds to an individual cell. *WT*: n=13, *Δseipin1.3*: n=9, *Δseipin8.3*: n=14, *OE-GFP8*: n=15, *OE- GFP9*: n=8. Statistically significant differences between mutants and WT were evaluated using multiple t- tests but none were found. C) Quantification of the size of LD in all the cell lines: WT (grey), PtSeipin KO (orange and red) and PtSeipin-GFP overexpressing lines (light and dark green). Results are presented as boxplots and each dot corresponds to an individual LD. *WT*: n=43, *Δseipin1.3*: n=24, *Δseipin8.3*: n=30, *OE-GFP8*: n=72, *OE-GFP9*: n=28. Statistically significant differences between mutants and WT were evaluated using multiple t-tests and are indicated as follows: *: p-value<0.05. **: p-value<0.01; *** p-value<0.001.

### High light exposure exacerbates the impact of PtSeipin KO

As LD formation in microalgae is highly induced by abiotic and biotic stresses, we next wondered whether this could be altered in PtSeipin mutant lines. Nutrient starvation stresses have been well characterized in *P. tricornutum*: nitrogen starvation leads to the rapid formation of large LD (Abida et al. 2015; Jaussaud et al. 2020), and a similar process occurs upon phosphate starvation, albeit at a slower pace (Abida et al. 2015). High light stress also leads to LD accumulation in microalgae (Napolitano 1994; Brown et al. 1996; Khotimchenko and Yakovleva 2005; Goold et al. 2016) but, at the best of our knowledge, the dynamics of LD formation in *Phaeodactylum* in this condition was not investigated. Unlike nitrogen and phosphorus starvation, high light exposure leads to transient accumulation of many small LD (supplementary Figure S11). The morphology of LD in PtSeipin mutants following nitrogen or phosphorus deprivation was not altered compared to the WT (Supplementary Figure S12). By contrast, the morphology of LD in PtSeipin KO lines upon high light (HL: 200 µmol photons.m^-2^.s^-1^ vs 75 µmol photons.m^-2^.s^-1^ in control conditions) was strongly altered, with a very strong increase in the size of LD and decrease of their number (Figure 7 and Supplementary Figure S13). Similarly to what was observed in control conditions, Seipin overexpression had no visible effect on LD morphology or cell morphology (Figure 7 and Supplementary Figure S13). In agreement with these data, no change of growth or Nile Red fluorescence levels were observed in the OE lines, while growth was slightly delayed in the PtSeipin KO lines, with a significantly higher Nile Red fluorescence at day 4 and 5 (Supplementary Figure S14). Finally, the PtSeipin KO cells morphology was more severely altered than in control condition, with shorter and larger cells (Figure 7A).

**Figure 7:**
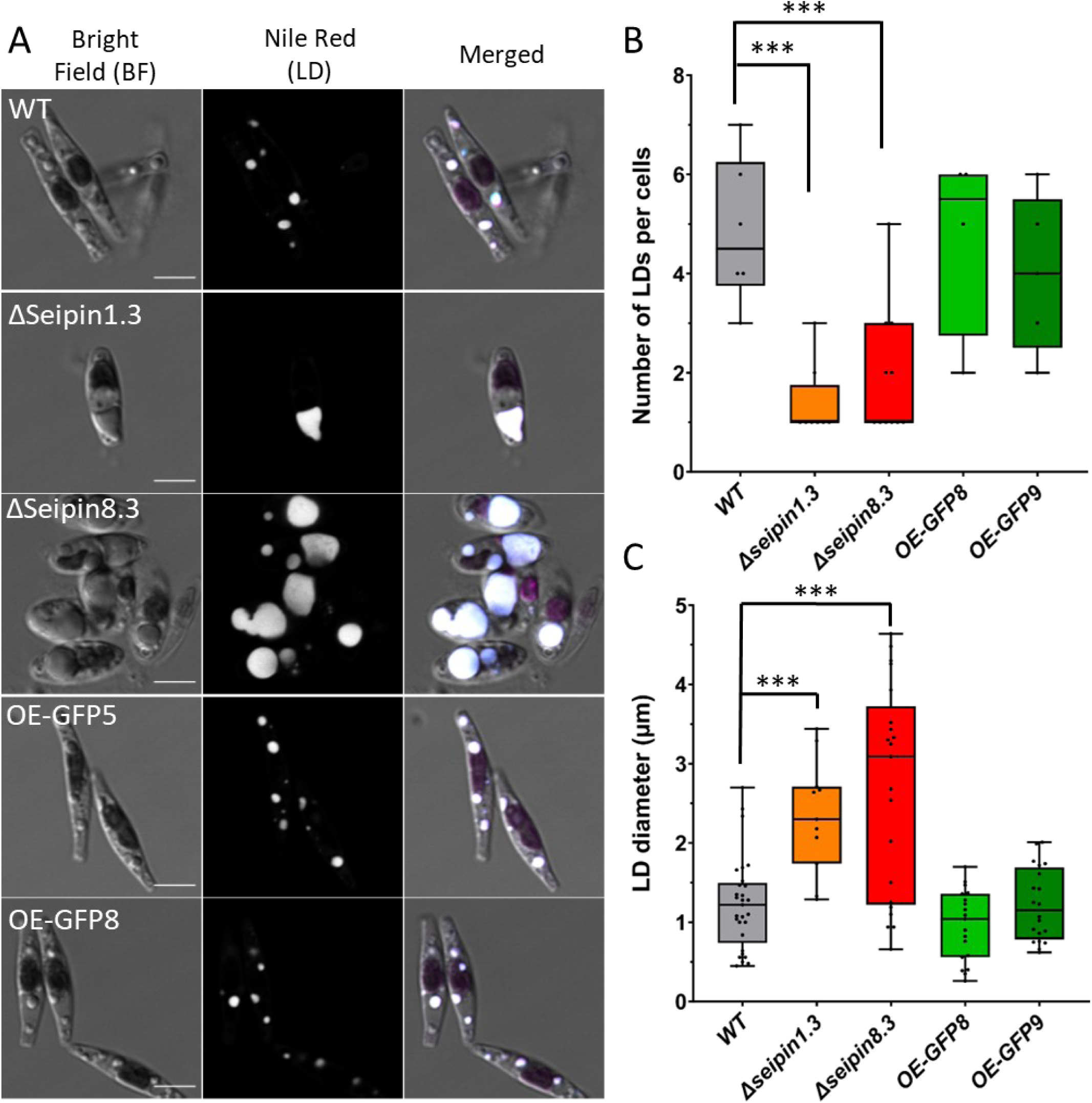
Observation and quantification of LD in PtSeipin mutants after 4 days of culture in high light (HL) culture conditions. A) Laser Scanning Confocal Microscopy (LSCM) images of WT, PtSeipin KO (ΔSeipin1.3 and ΔSeipin8.3) and PtSeipin-GFP overexpressing lines (OE-GFP5 and OE-GFP8) after staining with Nile Red. From left to right: pseudo-bright field (grey), Nile Red fluorescence (greyscale) and overlay of pseudo-bright field with Nile Red and chlorophyll signals (respectively grey, cyan and magenta). Scale bar: 5 µm. B) Quantification of the number of LD per cell in all the cell lines: WT (grey), PtSeipin KO (orange and red) and PtSeipin-GFP overexpressing lines (light and dark green). Results are presented as boxplots and each dot corresponds to an individual cell. *WT*: n=6, *Δseipin1.3*: n=8, *Δseipin8.3*: n=11, *OE-GFP8*: n=10, *OE-GFP9*: n=5. Statistically significant differences between mutants and WT were evaluated using multiple t-tests and are indicated as follows: *: p-value<0.05; **: p-value<0.01; *** p-value<0.001. C) Quantification of the size of LD in all the cell lines: WT (grey), PtSeipin KO (orange and red) and PtSeipin-GFP overexpressing lines (light and dark green). Results are presented as boxplots and each dot corresponds to an individual LD. *WT*: n=29, *Δseipin1.3*: n=11, *Δseipin8.3*: n=21, *OE-GFP8*: n=37, *OE-GFP9*: n=20. Statistically significant differences between mutants and WT were evaluated using multiple t-tests and are indicated as follows: *: p-value<0.05; **: p-value<0.01; *** p-value<0.001.

In addition to alterations of LD size and numbers, in both yeast and *Arabidopsis thaliana*, Seipin depletion leads to the formation of LD in the nucleus, originating from the nuclear inner envelope (Cartwright et al. 2015; Taurino et al. 2018). In order to verify whether similar alterations of LD biogenesis occurred in *Phaeodactylum*, we observed the cells using Transmission Electron Microscopy (TEM) after cryo-fixation. Figure 8 shows representative images of the WT and PtSeipin KO after high light exposure. The images confirm the alteration of the cell morphology observed by confocal microscopy, as well as the differences in the number and size of LD, but we did not observe the formation of any nuclear LD. Interestingly, as the use of cryo-fixation allows the preservation of the LD organization and unveils its heterogeneity, we observed the presence of darker regions, probably corresponding to the presence of a liquid phase of carotenoids (Lupette et al. 2019; Ezzedine et al. 2023) inside LD. The distribution of this spots appears different between the WT and the KO. While most of the WT LD contain one large dense region, the KO LD show a more variable distribution; some also contain one large dark region but others show a high number of very small dark areas, or even an absence of such dark areas (Figure 8).

**Figure 8:**
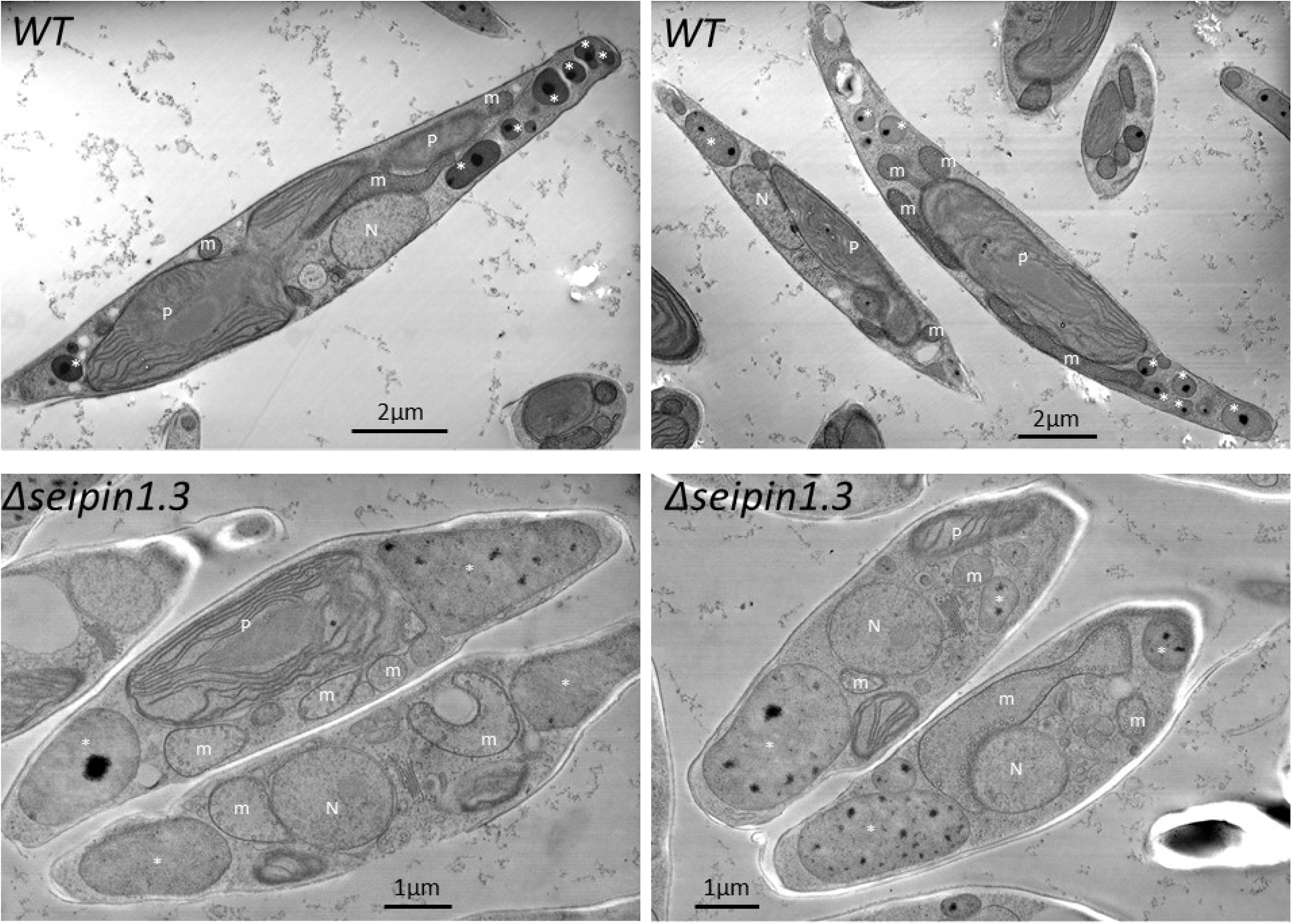
Transmission Electron Microscopy (TEM) observation of cryo-fixed WT and PtSeipin KO after 4 days of culture in high light (HL) culture conditions. N: nucleus; m: mitochondrion; P: plastid; *: Lipid droplets

We next wanted to observe whether the volumes of the various organelles were affected in PtSeipin KO. To this aim, we used focused ion bean scanning electron microscopy (FIB-SEM) to generate 3D electron images (Supplementary Movies 1 to 4). The obtained images were then segmented (Supplementary movies 5 to 8, Supplementary Figure S16) to determine the volume of the cells and of the internal organelles: LD but also nucleus, mitochondria and plastid (Figure 9A and Supplementary Figure S15A). Given the technical difficulties, we could only analyze two cells for the WT and two cells for the KO, and we selected one KO cell line, ΔSeipin1.3. As shown on Figure 9B and Supplementary Table 5, the two WT had very different cell sizes while the two KO had the same volume. Yet in both KO and WT we observed that the portion of the cell volume occupied by the nucleus was very stable, and similar between KO and WT. The volume occupied by mitochondria appeared more variable but no difference was observed between KO and WT. In agreement with what was observed using confocal imaging (Figure 7) the total volume of LD was higher in the KO than in the WT and we observed two very large LD in each KO cell (Figure 9A, Supplementary Figure S15A and Supplementary Table 5). One of the KO also contained a very high number of very small LD (Supplementary Figure S15 and Supplementary Table 5); the smallest ones (<0.1µm^3^) were probably too small to be seen by confocal microscopy, which is why we never observed such a high number of LD in the KO (Figure 7B). The other major difference was the volume occupied by the plastid, which was reduced in the two KO lines compared to the WT. Inside the plastid, the volume occupied by the pyrenoid was also reduced (Figure 9B, Supplementary Table 5).

**Figure 9:**
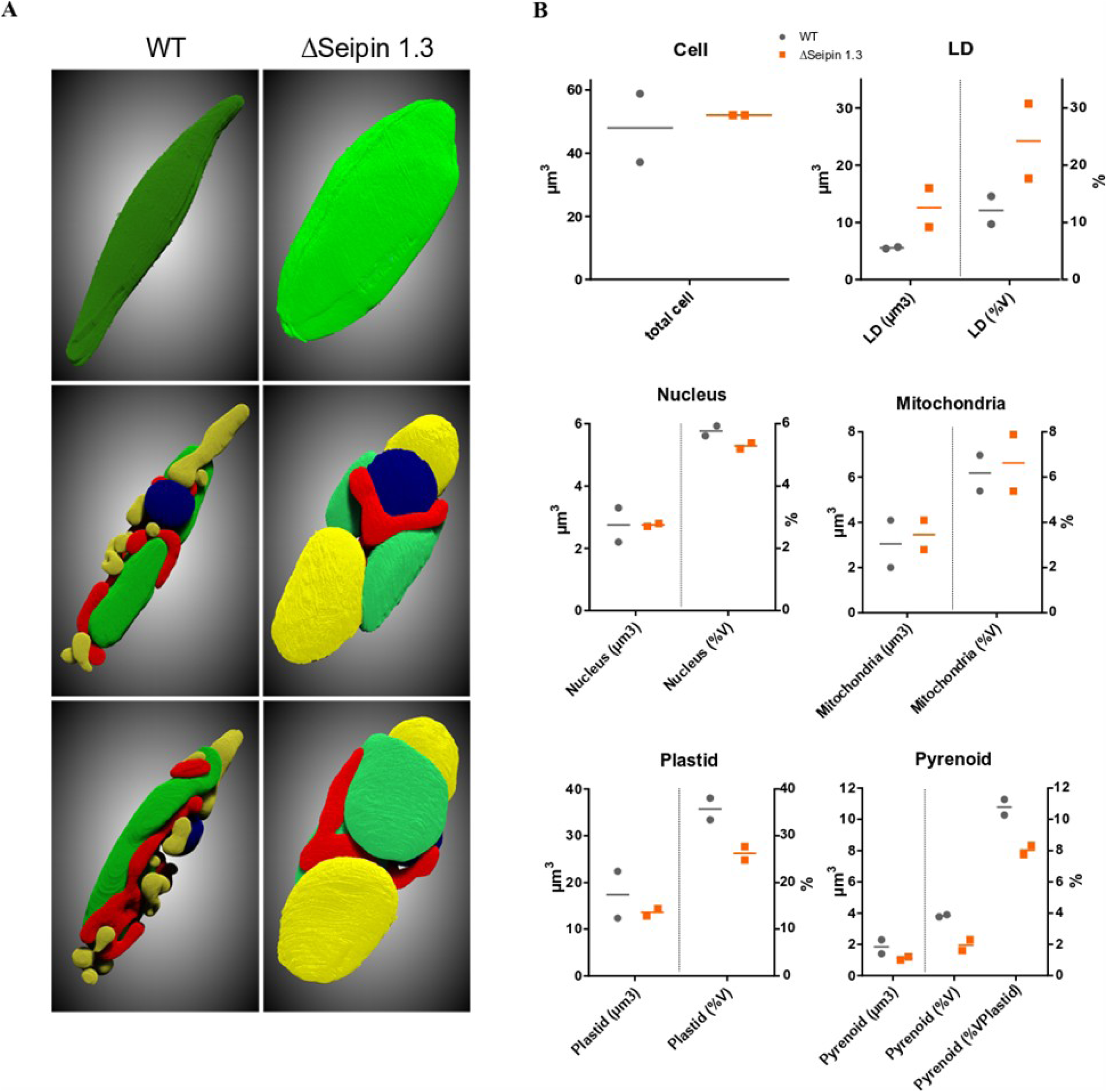
Segmentation of WT and PtSeipin KO following focused ion beam-scanning electron microscopy (FIB-SEM) and volumes of the different organelles. A. Segmentation of a WT (left) and a PtSeipin KO (right) cells. Upper panel shows the segmentation of the total volume while the two lower panels show two 3D views of the segmentation of the different organelles. Yellow: LD; red: mitochondria; blue: nucleus; green: plastid B. Measures of the volumes occupied by the total cell and the different organelle compartments within the cell. For all organelles, the raw volume (in µm^3^) and the relative occupancy of the cell as a percentage of the cell volume (%V) are shown. For the pyrenoid, the % of the plastid volume occupied (%VPlastid) is also shown. WT are shown as grey circles and ΔPtSeipin1.3 as orange squares. For each cell line, the two values, corresponding to the two segmented cells are shown.

### PtSeipin KO leads to enhanced TAG accumulation with only minor changes to TAG composition

The changes in LD size and number as well as higher Nile Red fluorescence signal, in both control and HL conditions, as well as the modification of the cell volume occupied by the plastid suggested that the lipid composition of PtSeipin KO was strongly affected. Moreover, higher TAG accumulation had also been described in PtSeipin OE lines (Lu et al. 2017). We thus addressed the lipid composition of PtSeipin mutants in both control and HL conditions. The glycerolipid profile (expressed in nmol/mg of dry weight) of PtSeipin KO and OE lines after 4 days of culture in control and HL conditions is presented on Figure 10, while the data obtained after 8 days of culture are presented on Supplementary Figure 17. As high variations in lipid content could sometimes be observed, results at day 4 are also shown in mol% in control and HL condition (Supplementary Figures S18C and D and S19 C and D respectively). Results combining all experiments on all KO lines are also shown as Supplementary Figure S20. No changes in glycerolipids composition are observed in the PtSeipin overexpressing lines at day 4 in control or HL conditions, apart from a small increase of MGDG in HL (Figure 10A and B, Supplementary Figures S18C and S19C). After 8 days of culture, a small but significant increase of TAG was observed in the OE lines in control conditions (Supplementary Figure 17A) while the other glycerolipids did not change. This TAG increase was not observed in HL conditions (Supplementary Figure 17B). In contrast to the minor changes observed in the OE lines, the Seipin KO lines showed a remarkable raise in TAG content in both conditions and at both time points (Figure 10, Supplementary Figures S17 A and B, S18C and S19C and S20A, B, E and F). In addition, PtSeipin KO lines showed small alterations of other glycerolipids (Figure 10, Supplementary Figures S17 A and B, S18D, S19D). These changes were not entirely consistent across experiments but some general trends can be drawn when considering all experiments together (Supplementary Figure S20). In control condition, the changes are mild but a relative decrease of DGTA at day 4 and 8 and of some phospholipids at day 8 can be observed when considering the relative percentage of glycerolipids with the exclusion of TAG (Supplementary Figure S20 C and D). However, alterations are much stronger in HL conditions and decrease of most phospholipids is observed at both time points, while surprisingly, galactoglycerolipids, which are the major lipids in the plastid, seem preserved (Supplementary Figure S20 E to H) in spite of the relative plastid size reduction (Figure 9, Supplementary Table 5).

**Figure 10:**
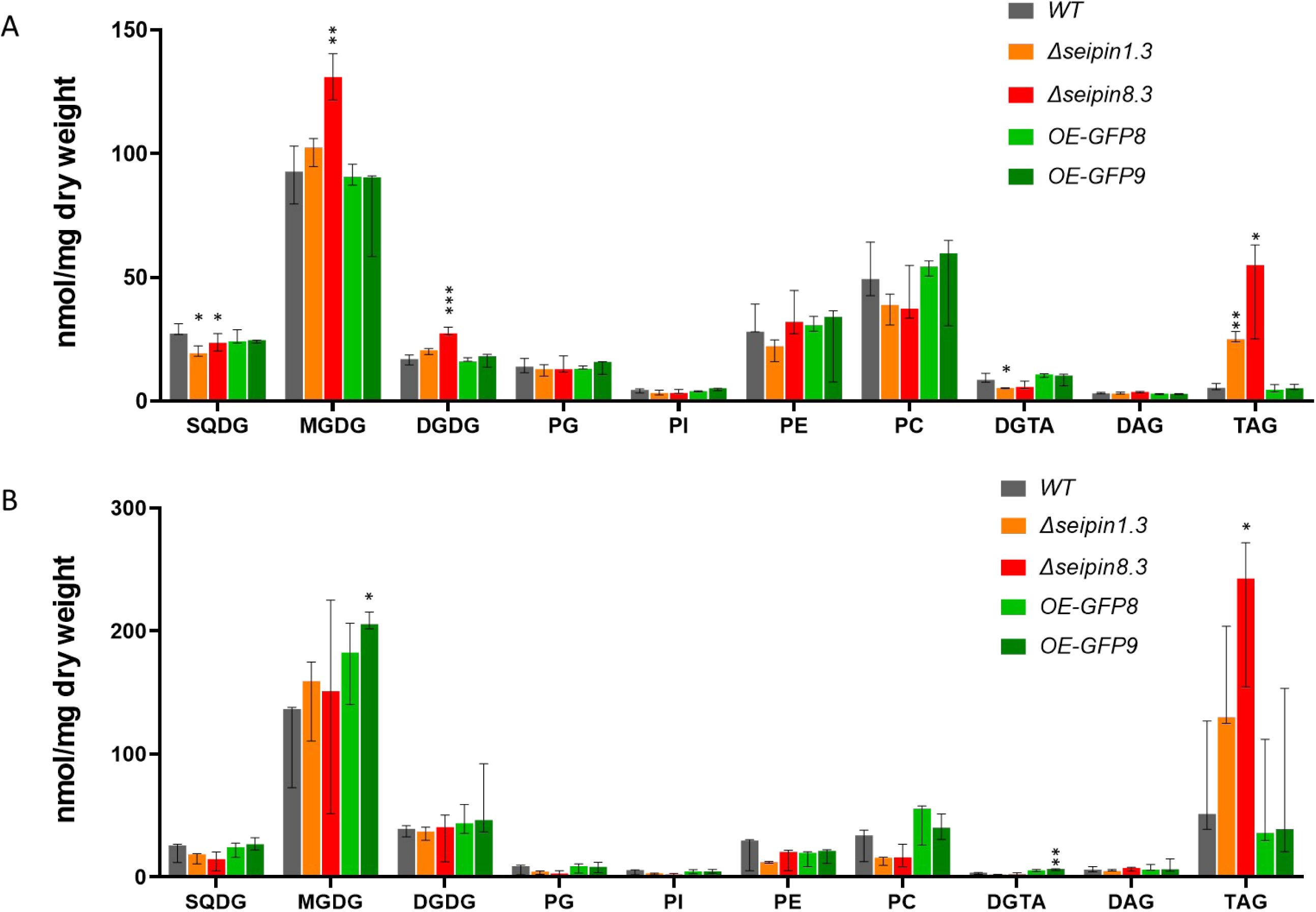
Glycerolipid profiles of WT and PtSeipin mutants after 4 days of culture in control condition (A) and high light condition (B). Glycerolipid classes were quantified following liquid chromatography and tandem mass spectrometry (LC-MS/MS) as described in the material and methods, and are presented in nmol/mg of dry weight. Median, min and max values of biological triplicates are shown. SQDG: sulfoquinovosyl-diacylglycerol; MGDG: monogalactosyl-diacylglycerol; DGDG: digalactosyl-diacylglycerol; PG: phosphatidylglycerol; PI: phosphatidylinositol, PE: phosphatidylethanolamine, PC: phosphatidylcholine, DGTA: ; DAG: diacylglycerol; TAG: triacylglycerol. Statistically significant differences between mutants and WT were evaluated using multiple t-tests and are indicated as follows: *: p-value<0.05; **: p-value<0.01; *** p-value<0.001.

We next looked whether changes in TAG content led to modifications in their fatty acids profiles. Figure 11 shows the distribution of all TAG species at day 4 in control (A) and HL (B) conditions, while the TAG distributions at day 8 are shown on Supplementary Figure S21. We did not observe any major changes in the TAG distribution of PtSeipin OE lines and PtSeipin KO compared to WT in both control and HL conditions (Figure 11 and Supplementary Figure S21). Major TAG species (TAG16:0_16:0_16_1 and 16:0_16:1_16:1) show only minor changes in a subset of cell lines and experiments, while the only consistent changes are observed in PtSeipin KO lines, with an increase in very minor species, that noticeably all contain C18:1 or C18:4.

**Figure 11:**
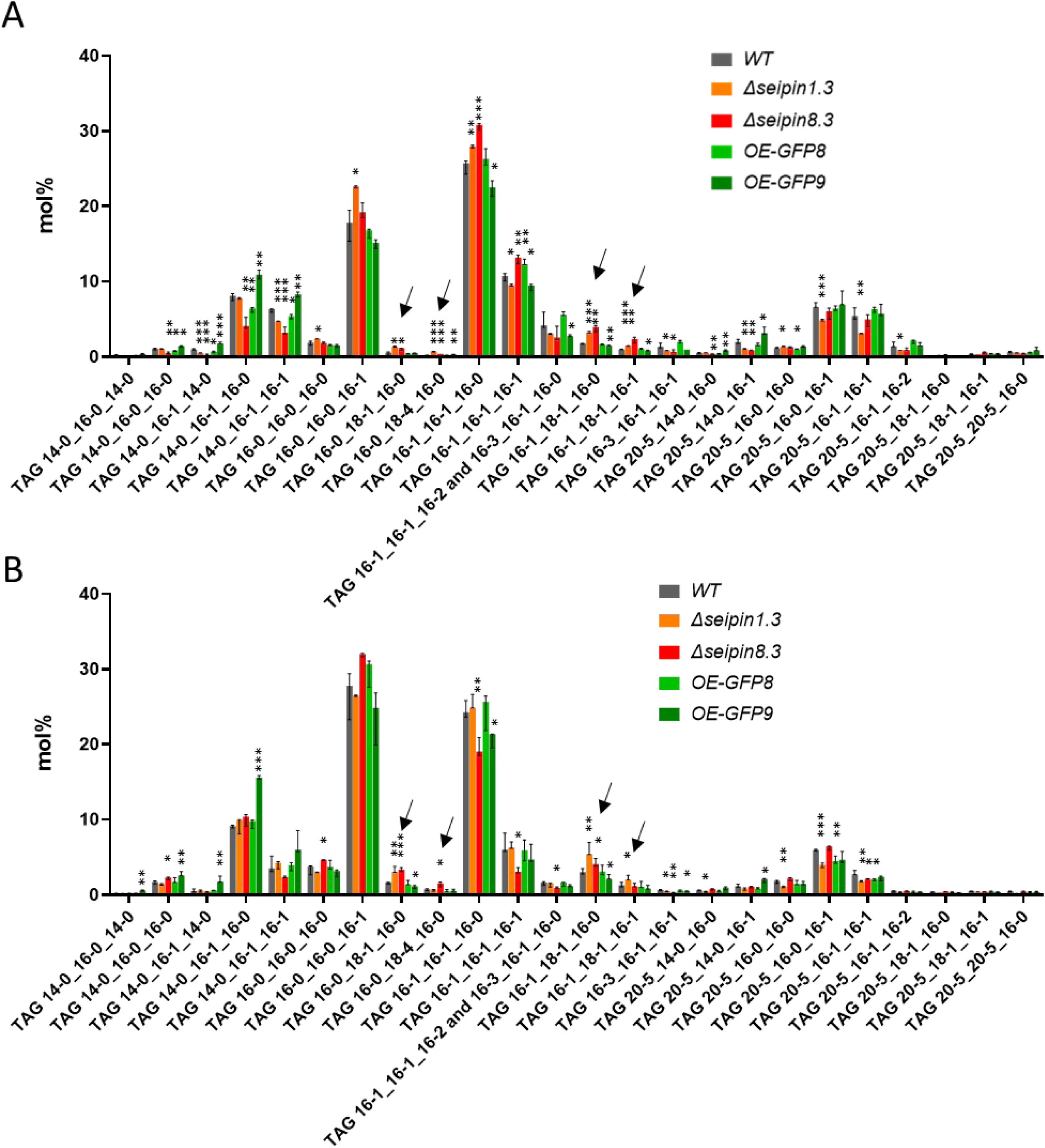
Triacylglycerol (TAG) profiles of WT and PtSeipin mutants after 4 days of culture in control condition (A) and high light condition (B). TAG species were quantified following liquid chromatography and tandem mass spectrometry (LC-MS/MS) as described in the material and methods, and are presented in mol%. Median, min and max values of biological triplicates are shown. Arrows indicate relative increase in species containing C18:1 and C18:4. Statistically significant differences between mutants and WT were evaluated using multiple t-tests and are indicated as follows: *: p-value<0.05; **: p-value<0.01; *** p-value<0.001.

### Differences in membrane glycerolipids composition

TAG accumulation can result from direct synthesis, from reduced degradation or from recycling of membrane glycerolipids. We investigated whether the composition of membrane glycerolipids was modified in PtSeipin mutants in control and HL conditions (respectively Supplementary Figure S18E to L and S19 E to L). The composition of the galactoglycerolipids MGDG and DGDG is not affected in mutants lines (KO and OE) compared to WT, in both control and HL conditions. On the other hand, the total quantity of phospholipids decreased in PtSeipin KO lines under HL conditions (Supplementary Figure S20 E and F) and most of the consistent differences are observed in PC and PE composition (Supplementary Figure S18 I and J and S19 I and J). In particular, PtSeipin KO lines show an increase in PE and PC species with a combination of 16:X and 18:X fatty acids (16-1_16-0, 16-1_16-1, 16-1_16-2, 16-1_16-3, 16-1_18-1, 16-1_18-2, 16-1_18-3) at the expense of classes with at least one C20:5 acyl group (20-5_16-0, 20-5_16-1, 20-5_16-2, 20-5_20-4, 20- 5_20-5 and 20-5_22-6). A similar trend is observed for the betaine lipid DGTA (increase of 16:0_16:1, 16:1_16:1, 16:1_18:1 and 16:1_18:2, and decrease of 14:0_20:5, 20:5_20:5, 20:5_22:5 and 20:5_22:6) (Supplementary Figures S18K and S19K). By contrast, the PC, PE and DGTA species containing combinations of 20:5 and 18:X do not seem affected. These results are coherent with the relative increase of C18:1, and decrease of C20:5 in the total FA pool (Supplementary Figures S18B and S19B). We observed little alteration of membrane lipids profiles in the OE lines, with only a slight increase in some species containing C20:5 in MGDG (MGDG 20:5_16:4 in control condition, MGDG 20:5_16:3 in HL) and PC (PC 20:5_20:4 and 20:5_20:5 in HL).

## Discussion

Lipid droplets in microalgae have been the object of many studies over the past years, mostly for their potential industrial applications. Nevertheless, basic knowledge regarding the mechanisms of LD biogenesis, and lipid metabolism in general, is still poor, especially in secondary endosymbionts. While often relevant, the reference to model plants has its limitations. In diatoms such as *Phaeodactylum tricornutum,* the complex intracellular architecture (Flori et al. 2016) and division mode (De Martino et al. 2009; Tanaka et al. 2015), the specificities of its major lipid droplets, raise specific questions regarding LD biogenesis. In the present paper, we investigated the function of *Phaeodactylum*’s Seipin homolog (PtSeipin) and we highlight conserved function as well as unique specificities.

### Evolution of Seipin sequences and interest for subsequent studies

Our study provides the first phylogenetic analysis of Seipins including a wide set of organisms resulting from primary and secondary endosymbiosis as well as animals and fungi (Opistokontha). Previous papers on plant and algae Seipins also included phylogenetic analyses that were either partial (Lu et al. 2017) or limited to the green lineage (Cai et al. 2015). Our results show that Seipins from organisms belonging to three clades of the SAR group (Oomycota, Eustigmatophyceae and Bacillariophyceae) cluster with those from Opistokontha, rather than with those of green algae and plants. These results suggest that Seipin from SAR were mostly inherited from the heterotrophic host rather than from the symbiotic algae, as already shown for *Nannochloropsis oceanica* PDAT (Yang et al. 2022). Nevertheless, the presence of rare Archaeplastida-specific motifs in some SAR species highlights that some recombination of both isoforms probably occurred independently in several species (Supplementary Table 2). This is in contradiction with the results of Lu and collaborators (Lu et al. 2017), which showed SAR Seipins as a sister group to plant Seipins. This difference can be at least partially explained by methodology, as the study from Lu et al. used the neighbor joining method while in our study we used a Bayesian analysis, including both information from the primary sequence and information based on the presence/absence and relative positions of motifs identified using MEME, thus providing more robust phylogenetic analyses. Our Bayesian inference was very robust and supported by very high posterior probabilities (Figure 2), whereas the previous phylogeny showed non-supported clades especially for the animal and fungi-plants-diatoms clades (Lu et al. 2017). Regarding the green lineage, our results are in agreement with those of Cai and collaborators (Cai et al. 2015), which showed the split between Seipin 1 on the one hand and Seipin 2 and 3 on the other. Moreover, our study suggests that the emergence of two and more isoforms of Seipin coincides with the apparition of seed plants, and that the Seipin from Bryophytes and non-seed vascular plants are closer to the Seipin2/3 isoforms found in seed plants. This result should be confirmed using a larger dataset of Bryophytes and vascular plants; though interesting, this is beyond the scope of this manuscript.

The analysis of motifs using MEME has also highlighted some very interesting features (Supplementary Figure S3 and Tables 2 and 3). Several general observations can be made. 1) The most conserved motifs, identified as ancestral (A, B, C and E), are located in the luminal domain. 2) Motifs identified in the transmembrane helixes are generally conserved within smaller groups of species (D in the N-terminal transmembrane helix of most SAR/Opistokontha, I and N in the C-terminal transmembrane helix respectively of most animals and of land plants, L in the N-terminal helix of vertebrates and plants and C-terminal helix of some green algae). 3) Conservation of motifs in the N-terminal part is even more restrained: domain Q is found only in vertebrate sequences and domain P is common to land plants, yet absent from Angiosperm Seipin 1 sequences. In both cases, the motifs are part of predicted α-helixes. 4) No conserved motif was detected in the C-terminal domain.

The localization of the motifs thus highlights some important features regarding conservation of structure as well as biophysical properties (in particular hydrophobicity), but their identification also hints at some new research lines. Indeed, the highly conserved motifs A, C and E correspond to regions that include the α −helices imbedded in the β-barrel (Supplementary Figure S4). α-helices 2 and 3 correspond to the hydrophobic helices (Sui et al. 2018) or membrane-bound helices (Yan et al. 2018), which are involved in Seipin oligomerisation and interact with lipids in the ER membrane. Several amino acids from α−helix 1 are also involved in the oligomeric interface according to Sui and collaborators (Sui et al. 2018), but only the serine (S117 in the human sequence, position 6 in motif A) appears highly conserved, while the other strongly conserved residues do not have identified functions. Investigating the function of this α-helix, as well as the function of Seipins in organisms in which it has been lost (*e.g. Chlamydomonas rheinardtii)* might thus be of high interest. Motif B is also located at a crucial point, the “kinked helix” located at the start of the C-terminal transmembrane domain and which shows functionally important conformational changes in yeast (Arlt et al. 2022). This motif appears to be ancestral (Supplementary Table 3 and Figure 2) but has evolved in the green lineage into motif J, as evidenced by the simultaneous identification of both motifs at the same position in some sequences (Supplementary Table 2). The functional significance of this shift, if any, remains to be investigated.

The development of *in silico* structure prediction tools, in particular AlphaFold, has allowed the comparison of PtSeipin structure with those of human, drosophila and yeast, which have been resolved by CryoEM (Sui et al. 2018; Yan et al. 2018; Klug et al. 2021; Arlt et al. 2022). This has highlighted the presence of extra-loops, with undetermined structures, surrounding the second luminal β-sheet in the sequences of PtSeipin, but also of other diatoms’ and Eustigmatophytes’ Seipins (Figure 1D and Supplementary Figure S2). Interestingly, while the links between β-sheets are unstructured, both the β1- β2 and the β2- β3 regions contain motifs that are conserved between animal sequences (motifs M and H) and between Archaeplastida sequences (motifs G and O). Yet the functional significance of these regions, and of the extra-loops identified in SAR remain to be investigated.

### PtSeipin localization supports a plastid contribution to LD biogenesis

In plant cells (Cai et al. 2015), yeast (Szymanski et al. 2007; Fei et al. 2008) and human cells (Ito and Suzuki 2007), Seipin has been shown to localize at discrete points in the ER, corresponding to LD biogenesis sites. Consistently with these results, in growing *P. tricornutum* cells, we observed that PtSeipin-GFP formed foci at the vicinity of LD and partially surrounded LD (Figure 4 and Supplementary Figure S6B). In starved cells, where LD are much larger, this surrounding of LD was even more pronounced (Supplementary Figure S7). In the strong expressing lines, a fainter signal was also observed in elongated structures, that expand to the cell extremities and correspond to ER-like staining (Liu et al. 2016), as well as around the plastid (Figure 4 and Supplementary Figures S6B and S7B). As this signal was not observed in the lines where PtSeipin-eGFP expression was lower (OE lines 5 and 9, Supplementary Figure S6), we cannot exclude that it is an artifact linked with overexpression of a tagged protein, but PtSeipin seems able to localize to the plastid external membrane as well as in the ER. Moreover, we observed that foci at the vicinity of LD, which probably correspond to LD biogenesis sites, can be located both at the plastid surface and in the ER (Figure 4). This corroborates previous observations suggesting that the plastid most external membrane, the cERM, contributes to LD formation (reviewed in (Le Moigne et al. 2022)) and may be functionally equivalent to the ER. Finally, as LD can be connected to more than one of this foci at a time, it seems that both cERM and ER can contribute together to LD formation, using TAG from different origins.

### PtSeipin and LD biogenesis

Modifications of PtSeipin expression lead to alterations of LD biogenesis. PtSeipin overexpression has only a minor impact: we do not observe any change in LD size, but LD numbers per cell tend to increase in the PtSeipin-eGFP OE lines at day 8, especially in the strongest OE line (OE-GFP8, Supplementary Figure S10). This trend is already visible at the end of the exponential phase (day 4, Figure 6), even though the differences were not statistically supported. This increase in LD number is also reported by Lu *et al*. (Lu et al. 2017), together with an increase in LD size, but the authors do not provide quantification data. Interestingly, there is no increase in LD number or size when OE lines are exposed to high light condition, (Figure 7 and Supplementary Figures S13), but this condition itself induces the formation of many small droplets in the WT (Figure 7, Supplementary Figures S11, S12 and S13).

By contrast, striking phenotypes are observed following PtSeipin depletion. In non-stressed conditions, we observed that the size of LD was heterogeneous among cells, with LD that could reach up to 2 µm in diameter (Figure 6, Supplementary Figure S10), a size that is normally observed only following stress such as nitrogen starvation (Jaussaud et al. 2020). This phenotype is strongly exacerbated after culture in high light condition (Figures 7 and 9, Supplementary Figures S13 and S15), with the formation of LD that can reach up to 4 µm, a giant size that is permitted by changes in the cell morphology, as cells appear shorter and larger (Figures 7 and 9, Supplementary Figure S15). Formation of very small LD is also observed in some cells, aside with the very large ones (Supplementary Figure S15, Supplementary Table 5). The alteration of LD morphology is a conserved feature following Seipin depletion. Indeed, in yeast, KO of either FLD1 or LDB16, the two proteins that associate to fill Seipin function, results in the formation of aberrant LD, with either giant LD or a clustering of very small LD (Szymanski et al. 2007; Fei et al. 2008; Wolinski et al. 2011). Similarly, in human, loss of Seipin results in the ectopic formation of giant LD in non adipogenic cell lines, and to LD absence in adipogenic cell lines (Fei et al. 2011). Finally, in *Arabidopsis thaliana*, loss of two or three Seipin isoforms results in the formation of giant LD in the seeds and developing embryos (Taurino et al. 2018). Moreover, in both yeast and plants, ectopic formation of LD budding from the nuclear inner envelop is observed (Cartwright et al. 2015; Taurino et al. 2018). Yet, our observations by TEM (transmission electron microscopy) and FIB-SEM (focused ion beam - scanning electron microscopy) following cryo-fixation of PtSeipin KO did not show any similar phenotype, even with the huge droplets obtained in HL condition (Figure 8, Supplementary Movies 1-4 and Supplementary Figure S16). This difference may come from the LD biogenesis sites, with a very probable involvement of the plastid most external membrane, as mentioned above, but we did not observe abnormal LD budding between the two most external plastidial membranes. Other mechanisms controlling proper LD budding thus likely exist in *P. tricornutum*.

Interestingly, the preservation of LD internal structure following cryo-fixation allowed the observation of dark structures inside the LD following HL stress. HL stress triggers the formation of reactive oxygen species (ROS) and many microalgae respond by the synthesis of antioxidant pigments, which can be stored in LD (Pick et al. 2019; Procházková et al. 2019). We previously observed the presence of beta-carotene in LD following nitrogen stress (Lupette et al. 2019); moreover, the presence of similar dark structures in LD was also observed in *Haematococcus pluvialis* (Kim et al. 2015) and in the snow alga *Sanguina nivaloides,* where it was attributed to the accumulation of another pigment, Astaxanthin (Ezzedine et al. 2023). The authors observed different stages of pigment accumulation in the LD, indicating that pigment loading is a controlled process (Ezzedine et al. 2023). In the WT, we observe that a large majority of LD contain only one large dark structure, while the KO LD show heterogeneity, with variations in the size and number of these dark structures. It is thus likely that the absence of PtSeipin and the formation of extra-large LD that ensues disturbs the pigments loading.

### TAG accumulation in PtSeipin mutants

Increased TAG accumulation is observed in parallel with LD biogenesis defects in both OE and KO lines. In the OE lines, TAG increase is only observed in control condition at day 8, when cells are in the stationary phase (Supplementary Figure S17A). This increase is moderate (×1.5 to ×3 depending on the cell line and experiment considered) and in a similar range to what was observed previously (×1.5 in (Lu et al. 2017)). However, we do not observe any consistent change in the TAG content at day 4, when cells are at the end of the exponential phase (Figure 11A). Finally, we did not observe any significant differences in TAG distribution between the WT and PtSeipin-eGFP OE lines at either time points (Figure 11), suggesting that there is no change in the TAG synthesis pathways. As LD number is higher in the PtSeipin OE lines (see above), it seems that PtSeipin overexpression in control conditions leads to an increase in LD nucleation sites, and that this in turn leads to faster TAG accumulation when cells are submitted to moderate nutrient deficiency but not when they are in replete conditions. Such an increase in LD numbers and TAG content is also observed with overexpression of *Arabidopsis thaliana* Seipins (Cai et al. 2015). By contrast, a reduction in Seipin levels is expected to lead to a reduction of LD nucleation sites, and thus to a reduction in TAG accumulation. This is what was observed in a recently published study showing that knock-down (KD) of the Seipin homolog in the eustigmatophyte *Nannochloropsis oceanica* leads to a reduction of TAG accumulation (Liu et al. 2024).

By contrast, a striking increase in TAG accumulation is observed following PtSeipin depletion, at all time-points and in both tested conditions (Figure 10 and supplementary Figures S17 to 20). In the absence of stress, TAG quantity show an 11-fold and a 6-fold increase at day 4 and day 8, respectively (Figure 10A, supplementary Figure S17A and Supplementary Figure S20A and B). Culture in high light condition affects TAG content in all cell lines with higher variability. Indeed, as shown on Supplementary Figure S11, LD accumulation in the WT is transient, as cells become acclimated to changes in light intensity. However, we observed that the dynamics of increase/decrease in LD and TAG content varied depending on experiments, hence a high variability in the WT, especially at later time points. In spite of this variability, TAG content in the KO lines was always higher (Figure 10B, supplementary Figure S17B and Supplementary Figure S20E and F), reaching up to 64% of total glycerolipids, while it stayed below 50% in the WT (Supplementary Figure 19C). Very interestingly, PtSeipin KO lines show nearly no alteration in growth in control condition (Figure 5A) and only very mild delay in HL condition compared to the WT (Supplementary Figure S14), and they appear as promising candidates for biofuels applications (Patent: (Le Moigne et al. 2024)). Although they were observed in different microalgae (respectively *N. oceanica* (Liu et al. 2024) and *P. tricornutum*), the difference between Seipin KD and KO suggests that Seipin plays a key regulatory role in TAG homeostasis, that is retained when Seipin levels are simply reduced but lost when Seipin is absent.

### Pull, Push or Protect?

The TAG accumulation observed in non-stressed conditions can result from three broad mechanisms that are summed up in the “Push-Pull-Protect” model (Vanhercke et al. 2014). “Push” refers to a global increase in lipid synthesis, “Pull” to modifications of the downstream uses of fatty acids, towards TAG rather than towards other lipids, and “Protect” to a decrease in TAG degradation. We did not observe any consistent increase in total lipid content, either in the OE or in the KO line, indicating that we have no “push” effect linked to variations in PtSeipin expression (Supplementary Figure S20 A, B, E and F).

Consistently with a “pulling” effect, we observe a relative decrease of the betain lipid DGTA and some phospholipids in control condition and a decrease of DGTA and all phospholipids in HL condition in the KO lines, while galactolipids synthesis appears unchanged or even slightly increased (Figure 10, Supplementary Figures S17, S18 C and D, S19 C and D and S20). Increase of TAG to the detriment of other lipid classes may result from increased activity of Diacylglycerol acyl-transferases (DGAT) enzymes, or from increased phospholipid recycling by the phospholipid:diacylglycerol acyl-transferase (PDAT). *P tricornutum* has one identified PDAT, and it has recently been shown that it is localized in plastid membranes and can use PC but also MGDG as its substrate (Pan et al. 2024). The lipid class profile is thus not consistent with an increased PDAT activity. Moreover, while we do observe some changes in the composition of PC, these changes are not correlated with changes in TAG composition (Figure 11, Supplementary Figures S18 and S19). By contrast with the one PDAT, several DGAT enzymes exist in *P tricornutum* (reviewed in (Guéguen et al. 2021)), and they have been shown to have different localizations, either in the plastid (DGAT2A and 2D) or on the most external plastidial membrane, that is continuous with the ER (DGAT1, DGAT2B, DGAT2C and DGAT3) (Huang et al. 2024), indicating that they are probably active on different DAG pools, inside and outside of the plastid. The preservation of MGDG and DGDG content, while other lipid classes are decreased, could thus reflect increased activity of specific DGATs upon PtSeipin depletion, most likely among those that are located in the cERM, where PtSeipin also localizes (Figure 4, Supplementary Figures S6B and S7), directing neo-synthetized FA towards TAG rather than towards phospholipids and DGTA synthesis. Alternatively, PtSeipin depletion could result in a general activation of DGATs, and relative preservation of galactolopids synthesis could result from additional regulatory mechanisms that aim to preserve the plastid integrity. MGDG synthase γ (MGDγ), which is localized in the cERM, could play a crucial role to ensure that MGDG (and DGDG) pools are preserved, possibliy to the detriment of other lipids synthesis (Guéguen et al. 2024). Finally, as PtSeipin localization suggests that it can coordinate LD biogenesis from the cERM and ER (Figure 4), loss of PtSeipin could unbalance the relative contributions of both compartments to LD biogenesis, and subsequently the DAG/TAG equilibrium in these compartments. Indeed, in the absence of Seipin, LD budding depends on TAG concentration but also on membrane composition and curvature (Thiam and Ikonen 2021). While their composition is unknown, cERM and ER could have different properties and ability to promote LD nucleation. Investigating how PtSeipin loss affects LD nucleation sites, and TAG synthesis enzymes will be of great interest. Thus far, while interactions between Seipins and various enzymes involved in lipid synthesis (GPATs, LPAATs, PAPs) have been identified (Sim et al. 2012, 2020; Talukder et al. 2015; Pagac et al. 2016), but no interaction of Seipins with DGATs have been described.

The third (non-exclusive) hypothesis is that PtSeipin depletion leads to reduced LD degradation. In *P tricornutum,* LD formation follows circadian rhythms as TAG are produced during the day, when photosynthesis is active, and degraded during the night to sustain cell division and general metabolism (Chauton et al. 2013; Jallet et al. 2016). TAGs are thus hypothesized to serve as a transitory storage of energy (Becker et al. 2018) but could also serve as a temporary storage of fatty acids that can be used for the formation of other glycerolipids upon cell division. As LD have contacts with many other organelles, including ER, plastid and mitochondria (Lupette et al. 2019), a function of LD in fatty acids and lipids trafficking can also be proposed as it exists in many other organisms (reviewed in (Le Moigne et al. 2022)). A failure to remobilize TAGs could thus result in a decrease of other lipids, as we observe here for betaine and phospho-lipids. The absence of changes in galactolipid content could indicate either that their production does not depend on TAG remobilization or, as suggested above, that additional regulatory mechanisms favor the formation of galactolipid above other glycerolipids classes. A recent report has highlighted the effects of depletion of the acyl-CoA binding protein PtACBP, that is involved in LD degradation, and show effects on all lipid classes including galactolipids (Leyland et al. 2024), but conditions were very different (*i.*e recovery following nitrogen starvation) from those used in our study. Pt Seipin depletion could lead to changes in the proteome of LD that impair LD degradation but this remains to be investigated.

## Methods

Unless specified otherwise, all chemical reagents were obtained from Sigma Aldrich.

### In silico analysis

*Phaeodactylum tricornutum* Seipin (*Pt*Seipin) coding gene (Phatr3_J47296) was recovered from Ensembl Protists database and the protein sequence B7G3W8_PHATC from UniProt database. Other protein sequences used for Seipin structure prediction in *Thalassiosira pseudonana*, *Microchloropsis gaditana* and *Microchloropsis salina* were recovered as described in the following part (Phylogenetic analysis). Transit peptide and transmembrane domains prediction was performed using the Phobius software (https://phobius.sbc.su.se; (Käll et al. 2007)). All predicted structures were retrieved from the AphaFold database or generated with ColabFold v1.5.5: AlphaFold2 (Mirdita et al. 2022) based on the AlphaFold pipeline (Jumper et al. 2021; Varadi et al. 2022). The resolved structure of Human Seipin (*Hs*Seipin) was recovered from Uniprot (PDB identifier, 6DS5) (Yan et al. 2018). All PDB files were processed with the UCSF ChimeraX software (version 1.5) (Pettersen et al. 2021).

### Phylogenenetic analysis

To delineate a sequence dataset for phylogenetic analyses, we employed PSI-BLAST (Altschul et al. 1997, 2005) and DELTA-BLAST (Altschul et al. 1997; Schäffer et al. 2001; Boratyn et al. 2012) using *Arabidopsis thaliana* (AT5G16460 SEIPIN1, AT2G34380 SEIPIN2, AT1G29760 SEIPIN3), *Chlamydomonas reinhardtii* (XP_042914764), *Phaeodactylum tricornutum* (Phatr3_J47296), *Homo sapiens* (NP_001116427), and *Drosophila melanogaster* (NP_570012) as queries. The initial dataset, primarily comprising eudicots, monocots, mosses, ferns, and green algae, resulted from intersecting PSI- and DELTA-BLAST outcomes for the first two species. The second dataset, encompassing stramenopiles, fungi, mammals, and yeasts, was populated through the intersection of results using the latter three species. A subset, featuring up to three representatives per major phylogenetic group, was established. For Rhodophyte sequences, a targeted DELTA-BLAST was executed. Selected sequences underwent analysis using various online tools to confirm the presence of transmembrane domains (an essential Seipin protein feature) and the characteristic Seipin domain. Notably, the *Saccharomyces cerevisiae* sequence was excluded due to the absence of the characteristic Seipin domain. BioEdit Sequence Alignment Editor (Hall 1999) was employed for sequence handling, and alignment was carried out using MAFFT version 7.407_1 (Katoh and Standley 2013) implemented in NGphylogeny.fr (Lemoine et al. 2019). The BLOcks SUbstitution Matrix (BLOSUM 80, (Henikoff and Henikoff 1992)) was utilized for multiple alignment. Phylogenetic analyses were conducted in MEGA X (Kumar et al. 2018). The substitution model for Maximum Likelihood (ML) was selected based on the Bayesian Information Criterion (BIC score), calculated using the best model finder tool in MEGA X, considering the corrected Akaike Information Criterion (AICc) and log Maximum Likelihood value (lnL). The Le-Gascuel (Le and Gascuel 2008) substitution model was applied, incorporating a discrete Gamma distribution to model evolutionary rate differences among sites (+G). The rate variation model allowed for some sites to be evolutionarily invariable (+I). The tree with the highest log likelihood was retained. Initial trees for the heuristic search via the Nearest-Neighbor-Interchange (NNI) method (DasGupta et al. 2008) were obtained automatically using Neighbor-Joining (Saitou and Nei 1987) and BioNJ algorithms (Gascuel 1997) based on pairwise distances estimated using a JTT model (Jones et al. 1992), and the topology with superior log likelihood value was selected. The ML analysis was supported by 5000 pseudoreplicates. Positions with less than 90% site coverage were removed, allowing for less than 10% alignment gaps and missing data at any position (partial deletion option). The same dataset used for ML was used for Bayesian analyses. The amino acid sequence dataset was complemented with the outcome of a motif search analysis. The Multiple Em for Motif Elicitation (MEME suite) Version 5.5.5 (Bailey et al. 2009) was used to find 20 motifs with length between 6 and 20 aa in the whole dataset. The motifs have been named and ranked by statistical significance, then annotated onto each sequence (Supplementary Figure S3). The distance between each ensuing motives was ranked as well (Supplementary Table2). The information was mathematically formalized and used as Partition 1 (the aa sequence alignment was set as Partition 2) to be used in MrBayes v3.2.7 (Ronquist and Huelsenbeck 2003; Ronquist et al. 2012) software. Two independent Metropolis Coupled Markov Chain Monte Carlo (MC3) analyses were run. Four chains (one cold and three heated, temperature parameter set at 0.10) per run drove the analyses. 2,000,000 generations were set with sampling every 100 generations and a burn-in period of 25 %. For the Partition 2, the model selection was done during the analysis by estimating the posterior probabilities of the different models together with their parameters. The average standard deviation of split frequencies at the end of the two parallel analyses reached 1.64×10^-3^ with an average Potential Scale Reduction Factor (PSRF) of 1.000 indicating excellent convergence of the chains (Gelman and Rubin 1992). The ancestral state of each domain and distance in main nodes of the resulting tree were calculated.

### Microalgae strain and culture conditions

The diatom *Phaeodactylum tricornutum* (ecotype Pt1 strain CCMP2561) named as Wild Type (WT) was used in this study for PtSeipin functional study *in vivo*.

*Phaeodactylum tricornutum* cells were cultivated in Enriched Sea Artificial Water (ESAW) medium [NaCl (326.7 mM), Na_2_SO_4_ (25 mM), KCl (8.03 mM), KBr (0.725), H_3_BO_3_ (0.372 mM), NaF (0.0657 mM), MgCl_2_-6H2O (47.18 mM), CaCl_2_-2H_2_O (9.134 mM), SrCl_2_-6H_2_O (0.082 mM), Fe-EDTA (8.17 μM), Na_2_EDTA-2H_2_O (8.3 μM), ZnSO_4_-7H_2_O (0.254 μM), CoCl_2_-6H_2_O (0.0672 μM), MnCl_2_-4H_2_O (2.73 μM), Na_2_MoO_4_-2H_2_O (6.12 nM), Na_2_SeO_3_ (1 nM), NiCl_2_-6H_2_O (6.27 nM), CuSO_4_-5H_2_O (0.039 nM), vitamine B8 (4.09 nM), vitamin B12 (0.738 nM), vitamine B1 (0.593 μM)], enriched with NaNO_3_ (0.549 mM) as the nitrogen source (except for nitrogen starvation) and NaH_2_PO_4_-H_2_O (0.022 mM) as the phosphorus source (except for phosphorus starvation). The medium was designated as ESAW 10N10P (Abida et al. 2015). Cells were cultivated in INFORS Multitron Pro incubators at 20°C with 12/12 light/dark cycles and 100rpm agitation. For control condition (CT), illumination at 75 µmol photons.m^-2^.s^-1^ was used and for High Light condition (HL), light intensity was raised to 200 µmol photons.m^-2^.s^-1^.

For growth kinetics, the cells were cultured in 20 mL volumes in glass Erlenmeyer flasks of 100 mL capacity to ensure good oxygenation. Cells were inoculated at an initial concentration of 1.10^6^ cells.mL^-1^ in triplicates. Daily samplings were performed during 8 days. Cell concentration was calculated by measuring the absorbance at 730 nm on a TECAN infinite M1000 pro and using a regression curve previously established in the lab (Conte et al. 2018). Neutral lipids accumulation was estimated after staining with Nile Red (9-diethylamino-5- benzo[a]- phenoxazinone, Thermofisher) for 20 minuts at a final concentration of 0.5 mg.mL^-1^, 20% Dimethylsulfoxide (DMSO). Nile Red emission (580 nm) was measured after excitation at 530 nm using a TECAN infinite M1000 pro (TECAN, Austria). Results were expressed in relative fluorescence unit per million cells.

For lipid analyses, total cultures were harvested after 4 or 8 days of culture, centrifuged for 10 minutes at 3500×g, the cell pellets were flash frozen in liquid nitrogen and kept at −80°C until lipid extraction.

### Generation of PtSeipin KO lines

Knock out (KO) lines for PtSeipin were generated using the CRISPR-Cas9 system (Nymark et al. 2017). Two independent single guide RNAs (sgRNA) targeting the Seipin coding region were designed using the PhytoCrispex tool (https://www.phytocrispex.bio.ens.psl.eu/CRISP-Ex/) and *P. tricornutum*’s reference genome. NGG was chosen as the Protospacer Adjacent Motif (PAM) sequence. Targets were selected with respect to their position in the *PtSeipin* (Phatr3_J47296) gene to be as close as possible to the start codon of the gene (Supplemental Figure S8). An additional BLAST was performed on *P. tricornutum* genome using the preselected CRISPR targets sequence to insure that no similar sequences were found elsewhere in the genome. Primers pairs corresponding to the chosen sgRNA sequences (Table 1) were annealed by heating 2,5 µl of each primer at 100 µM in 45 µl of annealing buffer (1 mM EDTA, 50 mM NaCl, 10 mM Tris pH 7.6) at 95°C for 4 minuts then allowing the mix to cool down at room temperature for 45 minuts.

**Table 1:**
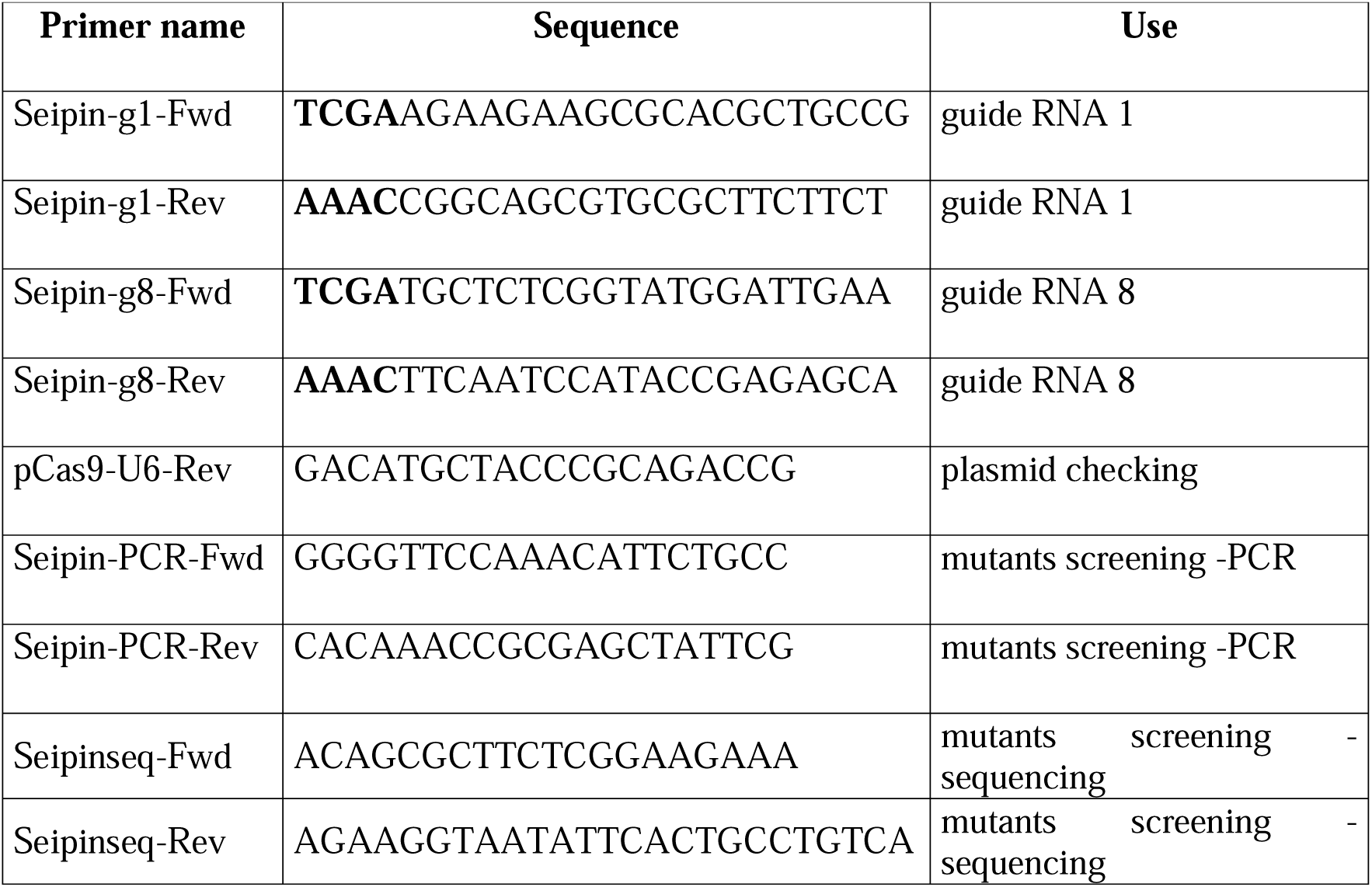
PtSeipin KO Primers table Added restriction sites are indicated in bold font.

The pKSdiaCAs9_sgRNA_ZeoR ((Seydoux et al. 2022), Addgene #74923) was digested by BsaI (New England BioLabs) for 1 hour at 37 °C. Annealing was performed with 1 µl of annealed sgRNA and 100 ng of linearized vector using T4 DNA Ligase (New England BioLabs). The ligation product was transformed in *E. coli* NEB5α (New England Biolabs). PCR screening with Phire Plant Direct PCR Master Mix (F160, ThermoFisher Scientific) was performed on bacteria colonies to assess the presence of the insert using the forward sgRNA primer (Seipin-g1-Fwd or Seipin-g8-Fwd) and the pCas9-U6-Rev primer (Table 1). Plasmids from positive colonies were extracted using the NucleoSpin Plasmid kit (Macherey-Nagel) and sent for sequencing to Macrogen (The Netherlands) using the M13-Rev primer. Plasmid midipreps (NucleoBond Xtra Midi kit, Macherey Nagel) were performed to obtain vectors with correct sgRNA insertions at a minimal concentration of 1µg/µl.

The single resulting vector (pKS diaCas9_sgRNA, Addgene #74923) was integrated into the genome of *Phaeodactylum tricornutum* following biolistic transformation as described in (Allorent et al. 2018) (adapted from (Nymark et al. 2017)). Briefly, 100.10^6^ *P. tricornutum* cells were spread on ESAW 10N10P/1% agar plates. Tungsten beads were coated with 5 µg of plasmid shot on the plates using the PDS-100/He system (Bio-Rad,Hercules, CA, USA; 1672257) with 1550 psi rupture disks. After 72 hours of recovery, the cells were plated on selective plates (ESAW10N10P/1 % Agar/Zeocin 0.07 μM) and after 4 to 6 weeks transformed colonies were recovered and screened. The sequence corresponding to PtSeipin was directly amplified from algae colonies using the Phire polymerase (ThermoFisher) with Seipin-PCR-Fwd and Seipin-PCR-Rev primer and sent for sequencing to Macrogen (The Netherlands) using Seipinseq-Fwd and Seipinseq-Rev (Table 1). Sequencing results were analyzed using the Tracking of Indels by DEcomposition (TIDE, (Brinkman et al. 2014)) software. Large deletions were identified using the alignment tool of the Snapgene software. Mutants containing at least 30 % of an interesting mutation were further purified as described in (Guéguen et al. 2024) by spreading 200 cells on a new selection plate and screening the obtained colonies then repeating the purification step until 100% pure mutant colonies were obtained. For each of the 2 selected guides, 3 independent pure KO lines were obtained and used in the present study (Supplemental Figure 8). For clarity purpose, results obtained with 2 lines, one for each guide, are shown in the main text.

### Generation of PtSeipin overexpressor lines

In order to overexpress PtSeipin in *Phaeodactylum*, both exons of PtSeipin coding sequence were inserted in the pPha-eGFP (Seydoux 2022) vector using Gibson assembly. For this, fragments corresponding to both coding exons of PtSeipin were amplified by PCR using High fidelity Phusion polymerase (ThermoFisher) with the following primers (Table 2): exon 1 was amplified with primer Seip-Ex1-pOx-F and Seip-Ex1-Ex2-R, exon 2 with SeipEx2-Ex1-F and SeipEx2-Linker-R. In parallel, the vector pPhaT1-linker-eGFP (Seydoux et al. 2022) was amplified and linearized by a single PCR with primers Linker-pOXGFP–F and pOX-SeipEx1–R. Amplified fragments were assembled by a single Gibson reaction (Gibson et al. 2009) before transformation into Neb5α bacteria. Plasmids were extracted using the NucleoSpin Plasmid kit (Macherey-Nagel) and sent for sequencing to Macrogen (The Netherlands) using primers S-pOX-1F, S-pOX-2R, S-pOX-3F and S-pOX-4R (Table 3). Plasmid midipreps (NucleoBond Xtra Midi kit, Macherey Nagel) were performed to obtain the transformation vector at a minimal concentration of 1 µg/µl. Transformation of algae was performed as described above. After 48h of recovery, cells were transferred to selection plates (ESAW10N10P/1 % Agar/Blasticidin 0.028 μM). After 4 to 6 weeks, transformed colonies were recovered and GFP expression was assessed by fluorescence microscopy and western blotting (see below).

**Table 2:**
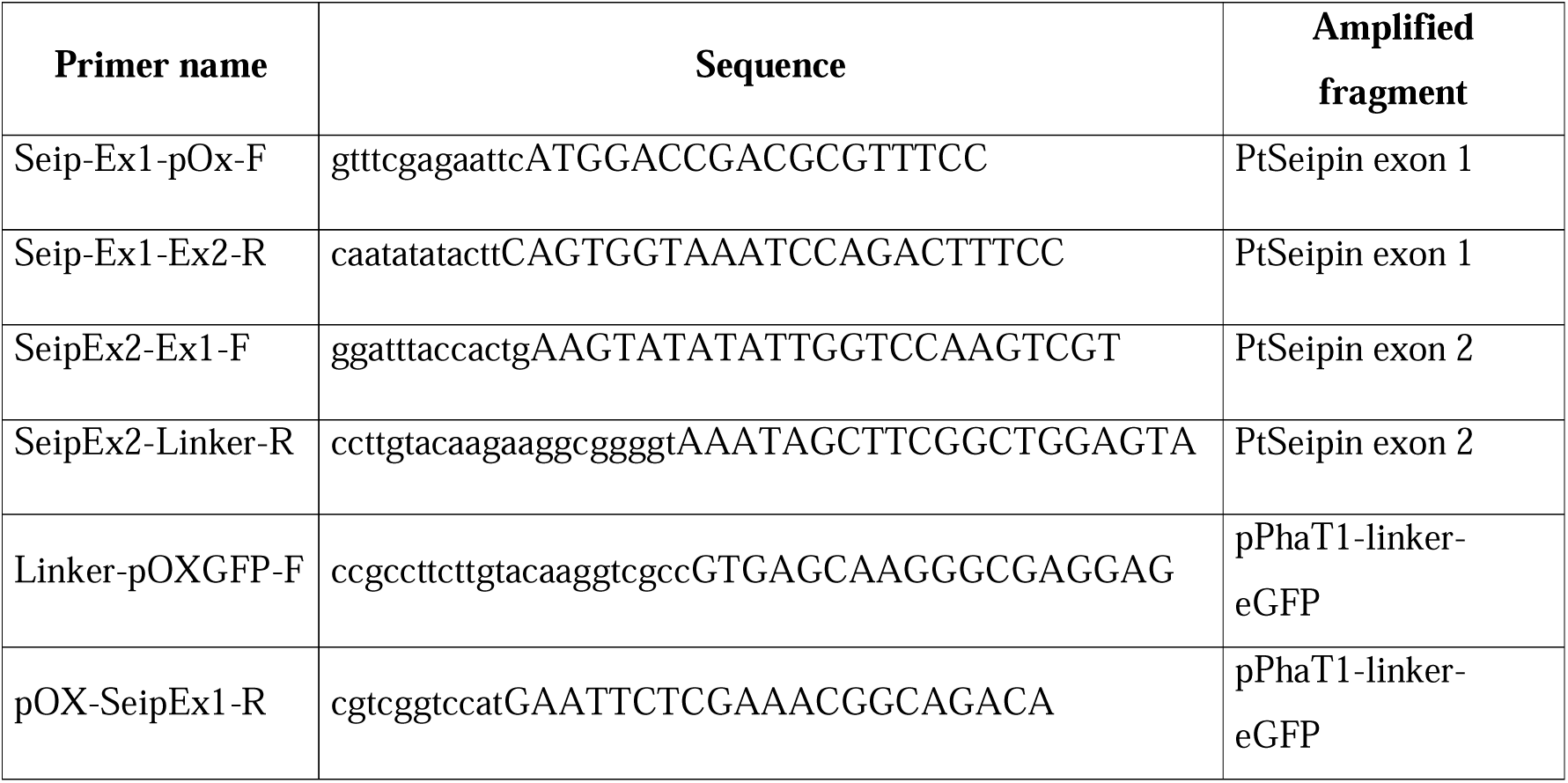
Primers used for generation of the PtSeipin overexpression plasmid. Sequences in caps correspond to the sequence of the amplified fragment and sequences in small font correspond to the part that were added for Gibson Assembly purpose

**Table 3:**
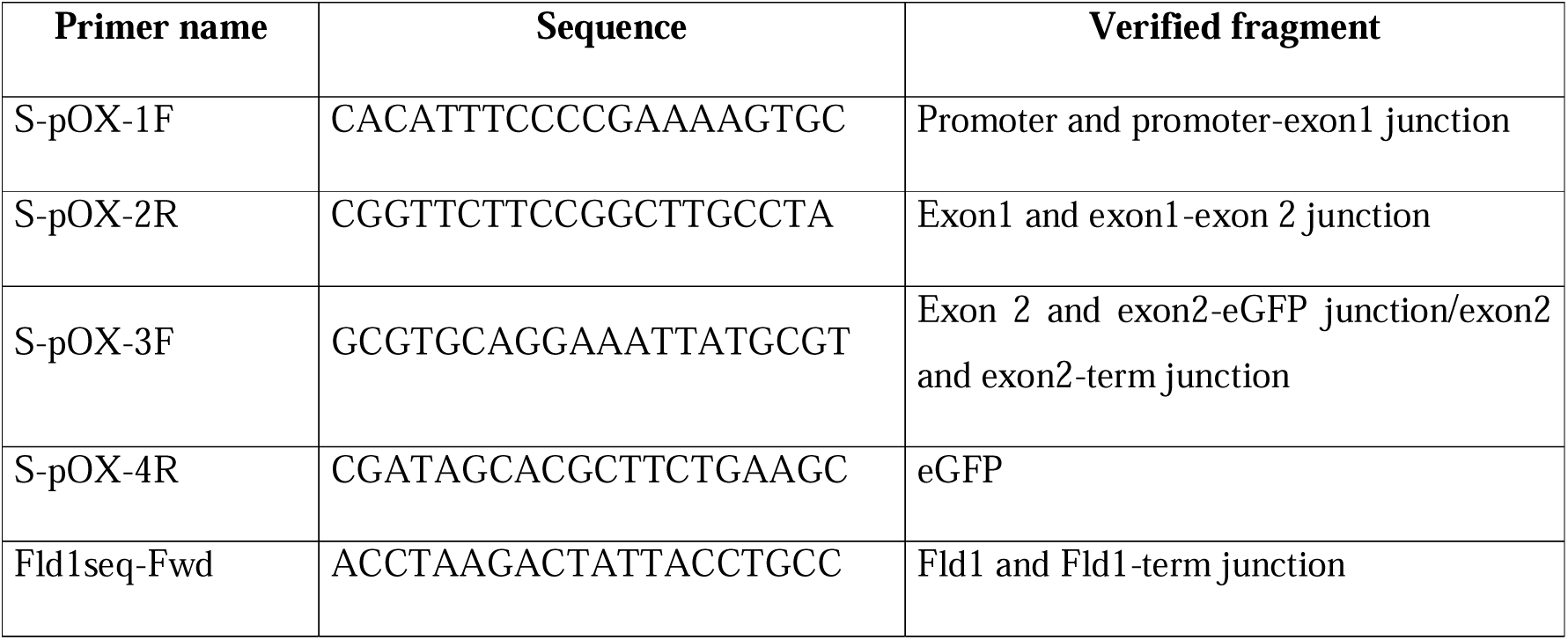
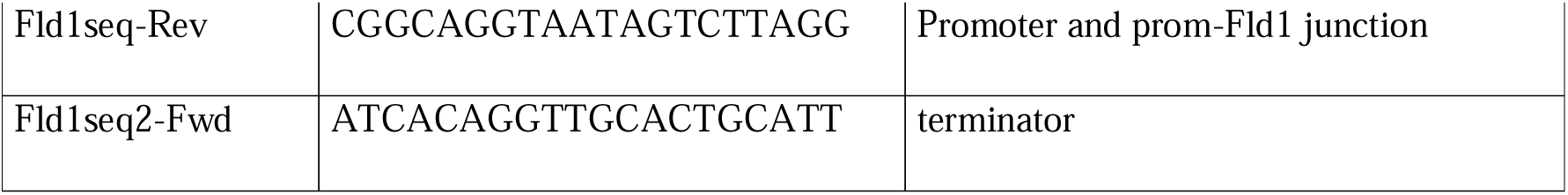
Primers used for sequencing of various plasmids.

Two independent series of mutant lines were generated using those constructions (PtSeipin-GFP5, 8 and 9 (*cf*. Supplementary Figure S6) and PtSeipin-GFP a and b (*cf.* Figure 4)) due to loss of the transgene expression in the original lines. Production of additional PtSeipin-GFP lines using different construction was later performed and is presented in Supplementary Material.

### Yeast complementation

#### Plasmids construction

Plasmids allowing the expression of PtSeipin and yeast Seipi1/Fld1 under control of the phosphoglycerate kinase (PGK) constitutive promoter were generated using the YCplac33 backbone. The PGK promoter and the Seipin1/Fld1 coding sequence were amplified from yeast genomic DNA; the ADH1 terminator coupled to 3xHA tag was amplified from plasmid pFA6a-3HA-His3MX6-n9 (Addgene Reference #41600); PtSeipin coding sequence was amplified from the pPha-PtSeipin-eGFP previously obtained. All fragments were amplified by PCR using the high fidelity Phusion enzyme (ThermoFisher) with the primers listed in Table 4: PGK was amplified using either primers 1+2 (YCplac33-PtSeipin) or 1+3 (YCplac33- Fld1), PtSeipin was amplified using primers 4+5, Fld1 using primers 6+7 and the terminator using either primers 8+9 (YCplac33-PtSeipin) or 8+10 (YCplac33-Fld1). YCplac33 was linearized by digestion with SmaI (New England Biolabs) for 1 hour at 25°C.

**Table 4:**
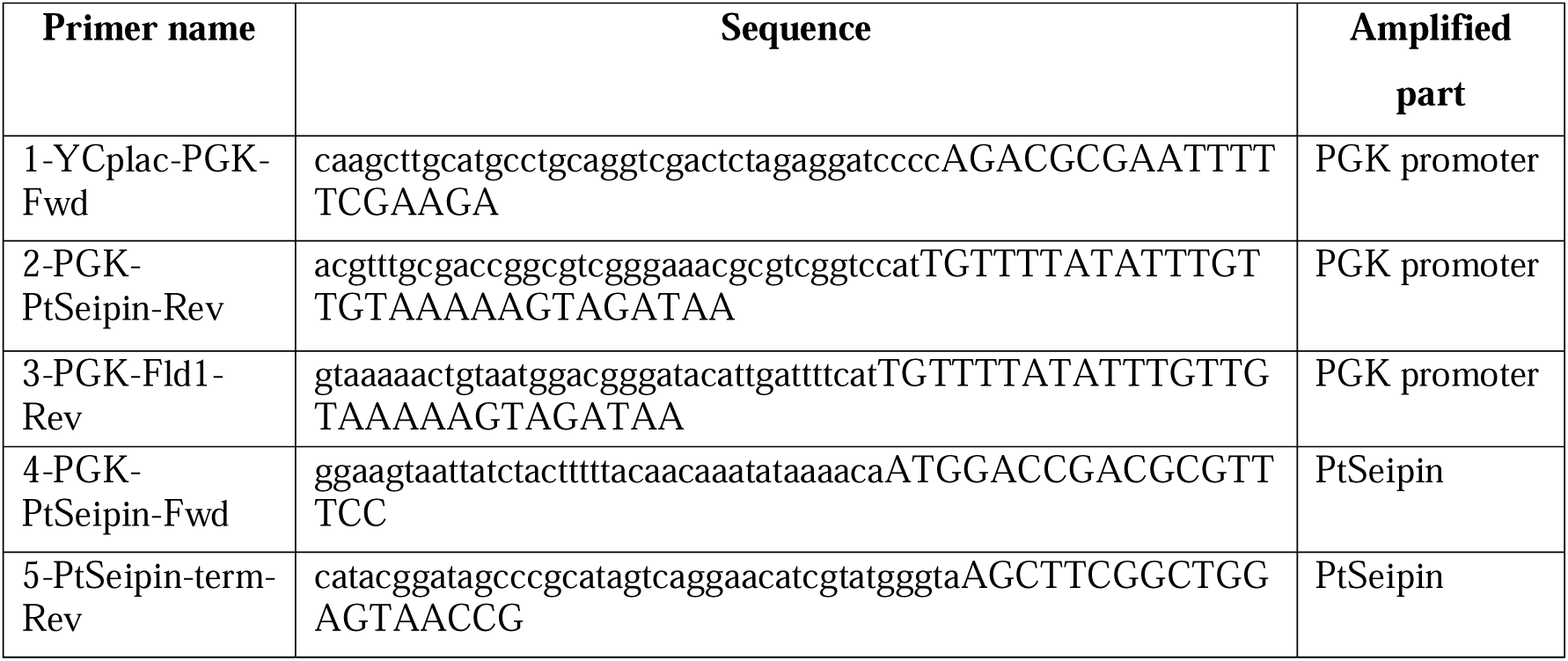
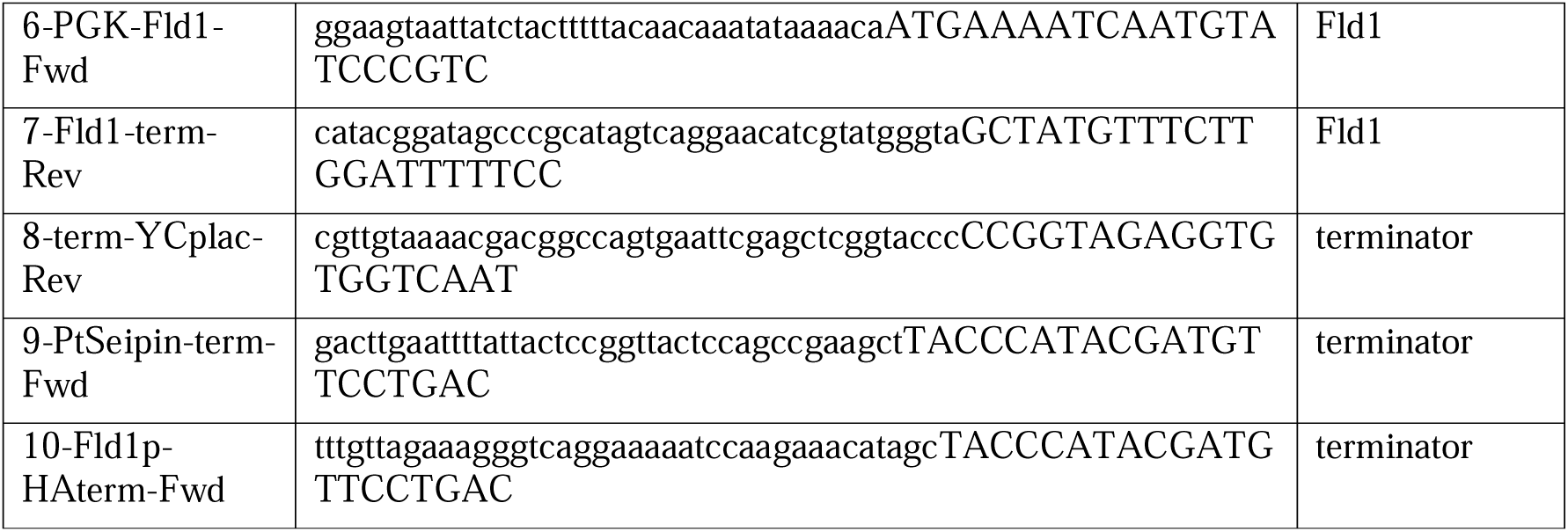
Primers used for generation of the yeast complementation plasmids. Sequences in caps correspond to the sequence of the amplified fragment and sequences in small font correspond to the parts that were added for plasmid assembly in yeast.

#### Yeast strains, culture and transformation

The yeast *Saccharomyces cerevisiae* Wild Type (WT, strain BY4741: MATa; his3Δ1; leu2 Δ0; met15 Δ0; ura3 Δ0) and KO for Seipin1/Fld1 (strain Ylr404w, kindly provided by Dr. Marine Froissard, Institut Jean-Pierre Bourgin, INRAE-Versailles, France) were used for functional complementation assays. Non-transformed yeast cells were cultivated in Yeast Peptone Dextrose complete medium (yeast extract 1%, peptone 2% and glucose D 2% w/v, YPD Broth Powder, MP Biomedicals). Transformed strains were grown in Yeast Nitrogen Base (YNB 6.7 g L^-1^, MP Biomedicals) supplemented with 0.77 g L^-1^ CSM-URA and 2% Glucose.

For complementation, the different fragments allowing reconstitution of an expression vector (YCplac33+PGK+(Fld1 or PtSeipin)+terminator) were transformed into the strain Ylr404w while the YCplac33 plasmid was transformed into the WT as a control. For transformation, cells in exponential phase (2<OD_600_<3) were suspended in LiAc-Sorbitol solution [1 M D-Sorbitol in TE Buffer (TE buffer: 100 mM Lithium acetate, 10mM Tris-acetate, 1 mM EDTA, pH8)] to make them competent. 50 µL competent cells are added to a mix containing 300 µL LiAc- PEG (40% m/v PEG 3350 in TE buffer), 5 µL denaturated salmon sperm DNA (10mg/mL)and DNA fragments (15ng linearized vector and 5 µL of each of the purified fragments). The cells were then successively incubated for 30 minutes at 30 °C, 15 minutes at 42 °C and plated on selective plates (Yeast Nitrogen Base (YNB) supplemented with CSM-Ura (MP biomédicals) and 2% Glucose, 2% agar). After 72 hours at 30°C, resistant colonies were recovered and screened: plasmid DNA was extracted using Zymoprep Yeast Plasmid Miniprep Kit (Zymo Research); the recovered plasmids were transformed into NEB5α bacteria and plasmid DNA was again extracted as described previously and sent for sequencing using primers S-pOX-2R and S-pOX-3F (for YCplac33-PtSeipin) and Fld1seq-Fwd, Fld1seq-Rev and Fld1seq2-Fwd (for YCplac33-Fld1) (Table 3).

To establish growth curves, a previously established protocol was used (Billey et al. 2021). Briefly, cells were inoculated at an optical density (OD_600_) of 0.05 in 300 µL volume in a transparent 96-well microplate (Thermo Scientific Nunc MicroWell 96-Well, Nunclon Delta-Treated, Flat-Bottom Microplate). Cells were incubated at 30 ± 0.5°C in an Infinite M1000 PRO microplate reader (Tecan) set for plate orbital shaking for 15 s every 20 minutes and absorbance measurements (OD_600_) every hour for 82 hours, with four measurements per well. For microscopy observations, cells were inoculated at low density (OD_600_=0.01) in a final volume of 30mL (100mL Erlenmeyer flask) and observed when OD_600_ was between 2 and 3. As cells complemented with PtSeipin showed slower growth, cells were observed both at the same time as the other lines with a lower OD_600_ (PtSeipinT1) or at a later time point with a OD_600_ (PtSeipinT2). This experiment was repeated 3 times.

### Lipid Analysis

All analyses were realized on the LIPANG platform and performed on biological triplicates except for a preliminary experiment, which was done on the six KO lines in two different conditions on biological duplicates. Additional experiments including the two KO lines and the two OE lines shown in the main text were repeated 3 times. Statistical analyses were performed using GraphPad Prism (v6.07).

#### Lipid extraction

Cell pellets stored at −80 °C were freeze-dried overnight (Christ Alpha 2-4 LDplus). Pellets were weighted and lipid extraction were performed according to the Folch method (Folch et al. 1957; Jouhet et al. 2017). Briefly, frozen pellets were ground in a Potter-Elvehjem mortar and resuspended in boiling ethanol. 2 mL of methanol and 8 mL of chloroform were sequentially added and mixed using argon bubbling. After 1 hour at room temperature, the samples were filtrated through glass wool which was rinsed with Chloroform/Methanol (2:1, v/v). Biphase formation was initiated with NaCl 1% (w/v) and centrifugation at 2000 rpm for 10 minutes. The lower organic phase was collected and dried at 40°C under argon flux and stored at −20°C.

#### Methanolysis and Gas chromatography coupled to fire induced detector (GD-FID)

Lipids were resuspended in methanol and 10% were sampled for methanolysis. 5 µg of 15:0 internal standard (SIGMA) was added to each sample for absolute quantification. Methanolysis medium composed of methanol:sulfuric acid (40:1, v/v) was added and samples were heated for 1 hour at 100 °C. The reaction was stopped by addition of 3 mL of water and formation of a biphase was obtained after addition of 3mL of hexane and vortexing. The upper phase containing fatty acids methyl-esters (FAMEs) was collected, evaporated under argon and resuspended in 100µL of hexane. FAME samples were then run on a gas chromatographer coupled to fire ionisation detector (Perkin Elmer Clarus 580) with nitrogen as a vector gas and a SGE BPX70 column. FAME were identified by comparison of their retention times with those of standards (Sigma) and quantified by the surface peak method using 15:0 for calibration thanks to a home-developed software (available upon request). The total quantity of fatty acids was used to determine the amount needed for the mass spectrometry analysis.

#### Mass spectrometry analysis

Lipid fractions were then analyzed by high-pressure liquid chromatography coupled to tandem mass spectrometry (HPLC-MS/MS) using appropriate standard lipids as described in (Jouhet et al. 2017). Quantities corresponding to 25 nmol of total fatty acids were sampled and resuspended in 100 μL of chloroform/methanol [2/1,v/v] containing 125 pmol of each internal standard.(phosphatidylethanolamine (PE) 18:0–18:0 and diacylglycerol (DAG) 18:0– 22:6 from Avanti Polar Lipid, and sulfoquinovosyldiacylglycerol (SQDG) 16:0–18:0 extracted from spinach thylakoid (Demé 2014) and hydrogenated (Buseman 2007)). 20 µL of lipids were then injected and separated by HPLC using an Agilent 1200 HPLC system with a 150mm×3mm (length×internal diameter) 5 μm diol column (Macherey-Nagel), at 40 °C. The mobile phases were hexane/isopropanol/water/1M ammonium acetate, pH 5.3 [625/350/24/1, v/v] (A) and isopropanol/water/1M ammonium acetate, pH 5.3 [850/149/1, v/v] (B). For elution, the percentage of B was originally set at 0 for 5 minutes and was increased linearly to 100% during 30 minutes then kept at 100% for 15 minutes. This was followed by a rapid return to 100% A in 5 minutes and the column was equilibrated for 20 minutes with 100% A before the next injection, hence a total runtime of 70 minutes with a flow rate of the mobile phase at 200 μL/minute. The distinct glycerophospholipid classes were eluted sequentially depending on their polar head group. Mass spectrometric analysis was performed on a 6460 triple quadrupole mass spectrometer (Agilent) equipped with a Jet stream electrospray ion source. The following settings were used: drying gas heater at 260 °C, drying gas flow at 13 L·minute^−1^, sheath gas heater at 300 °C, sheath gas flow at 11 L·minute^−1^, nebulizer pressure at 25 psi, capillary voltage at ±5000 V and nozzle voltage at ±1000 V. Nitrogen was used as collision gas. The quadrupoles Q1 and Q3 were operated respectively at widest and unit resolution. Phosphatidylcholine (PC) and diacylglyceryl hydroxymethyltrimethylCβCalanine (DGTA) analyses were carried out in positive ion mode by scanning for precursors of m/z 184 and 236 respectively at a collision energy (CE) of 34 and 52 eV. SQDG analysis was carried out in negative ion mode by scanning for precursors of m/z –225 at a CE of –56 eV. Phosphatidylethanolamine (PE), phosphatidylinositol (PI), phosphatidylglycerol (PG), monogalactosyldiacylglycerol (MGDG) and digalactosyldiacylglycerol (DGDG) measurements were performed in positive ion mode by scanning for neutral losses of 141 Da, 277 Da, 189 Da, 179 Da and 341 Da at CEs of 20 eV, 12 eV, 16 eV, 8 eV and 8 eV, respectively. DAG and triacylglycerol(TAG) species were identified and quantified by multiple reaction monitoring (MRM) as singly charged ions [M+NH4]+at a CE of 16 and 22 eV respectively. Quantification was done for each lipid species by multiple reaction monitoring (MRM) with 50 ms dwell time with the various transitions previously recorded (Abida et al. 2015). Mass spectra were processed using the MassHunter Workstation software (Agilent) for identification and quantification of lipids. Lipid quantities (pmol) were corrected for response differences between internal standards and endogenous lipids and by comparison with a qualified control (QC), thanks to an in-house software developed by the LIPANG platform. QC extracts correspond to a known *P. tricornutum* lipid extract previously qualified and quantified by thin-layer chromatography and gas-chromatography coupled to ion flame detection as described in (Abida et al. 2015).

### Protein analysis

#### Protein extraction

*S.cerevisiae* cell pellets were freeze-dried (Christ Alpha 2-4 LDplus) overnight. Dried pellets were resuspended in 200 µL lysis buffer (8 M Urea, 1 mM EDTA, 50 mM Tris-HCl pH 6,8) and about 50 µL of 0.5 mm glass beads (Sigma, USA) was added before heating 5 minutes at 70°C. Cell lysis was performed on a PreCellys 24 Touch (Bertin Technologies, France) set up on 3 cycles of 20 seconds at 10000 rpm. Heating and lysis steps were performed twice. The lysate was centrifuged 5 minutes at 5000×g and at 4 °C and the supernatant was collected.

*P.tricornutum* cell pellets were freeze-dried (Christ Alpha 2-4 LDplus) overnight. Dried pellets were resuspended in 700 µL “TANAKA” buffer (sucrose 0.7 M, Tris 0.5 M, HCl 30 mM, EDTA 50 mM, KCl 0.1 M, PMSF 2 mM, β-mercaptoethanol 2 mM, protease inhibitor cocktail pellet). 750 µL Phenol was added and the mix was vortexed for 30 seconds then centrifuged (20000×g, 5 minutes, 4°C) to create a biphase. About 500 µL of the upper pigmented phase containing protein was recovered. Proteins were precipitated at −20 °C overnight after addition of 1 mL 0.1 M ammonium acetate / ice-cold methanol (−20 °C). Proteins were pelleted by centrifugation at 20,000×g for 30 minutes at 4 °C, then washed with 1mL of 0.1M ammonium acetate / ice-cold methanol. A final wash with 100% ice-cold acetone was performed and pellets were dried in a fume hood for 30 minutes and resuspended in lysis buffer (8 M urea, 1 mM EDTA, 50 mM Tris-HCl pH 6.8) supplemented with 5% β- mercaptoethanol.

Protein concentration was measured using Protein Assay Dye Reagent (BioRad). A standard curve was established using Bovine Serum Albumin (BSA) with concentrations between 0 and 10 μg/mL. Samples were diluted, homogenized by vortexing and incubated at room temperature with Protein Assay Dye Reagent for 5 minutes before absorbance reading at 595 nm (Eppendorf Biophotometer D30).

#### SDS-Page and Western blotting

Samples were mixed with denaturing buffer (LDS (4X) NuPAGE™, Life technologies) and reducing agent (Sample reducing agent (10X) NuPAGE™, Life technologies). The mixture was heated to 95 °C for 5 minutes. Samples were loaded onto a polyacrylamide gel (NuPAGE™ 4 at 12% Bis-Tris minigel, Life Technologies) and migration was performed in a Mini Gel Tank (Life Technologies) at 100 V for 10 minutes, then 120 V for approximately 90 minutes.

Protein transfer from polyacrylamide gels onto nitrocellulose membranes (Amersham Protran 0.2 μm) was performed a Mini Gel Tank (Life Technologies) filled with a transfer buffer (Laemmli 1×: Glycine 192 mM, Tris-HCl 25 mM, pH 8.3, 10% ethanol) at 90 V for 90 minutes. Following transfer, membranes were washed in distilled water, incubated for 2 minutes in ponceau red [ponceau red 0.2% (w/v), TCA 3% (v/v)] and pictures were taken using a GelDoc XR imaging system (Biorad). The membranes were then washed in Tris-buffered saline (TBS: Tris-HCl 20 mM, NaCl 150 mM, pH 7.6).

For immunolabelling, the membranes were incubated at room temperature with stirring for at least 2 hours in 5% (w/v) TBS-milk buffer. Anti GFP primary antibodies [coupled with HRP, Ref 130-091-833 Miltenei (Biotech)] were used at 1:5000 dilutions in TBS1X, 0.5% milk, Tween 0.1%. Revelation was made with Clarity™ and Max Clarity™ Western ECL (BioRad, USA) according to the manufacturer’s recommendations. The luminescence was revealed at 428nm (Amersham ImageQuant 800).

### Microscopy

#### Epifluorescence microscopy

Cells were observed using an epifluorescence microscope (Zeiss AxioScope A1) equipped with a Zeiss AxioCam MRc. Images were captured using a Zeiss EC Plan Neofluar 100x/1.3 oil immersion objective. Chlorophyll autofluorescence and Nile Red fluorescence in lipid droplets were visualized with Zeiss filter set 16 (BP 485/20, FT510, LP515) and eGFP was visualized with Zeiss filter set 38 (BP 470/40, FT495, BP525/50).

#### Laser Scanning Confocal microscopy

Confocal microscopy observations were carried out using a Zeiss LSM880 microscope equipped with a 63x/1.4 oil-immersed Plan-Apochromat objective, running Zen 2.3 SP1 acquisition software (Platform µLife, IRIG, LPCV). Chlorophyll autofluorescence, eGFP fluorescence and Nile Red fluorescence were excited using the 488 nm ray of an Argon Multiline laser, and were detected at 650-700 nm, 500-530 nm, respectively and 580-640 nm respectively, using a Fast AiryScan detector. ‘Pseudo brightfield’ images were acquired in parallel by differential interference contrast (DIC), using the 488 nm laser ray as light source and a photomultiplier tube detector for transmitted light (T-PMT). Z-stacks containing consecutive images with a distance of 0.15 µm were obtained for each cell. All images were processed using the freeware ImageJ 1.53t version Fiji.

#### Cell cryo-fixation, and sample preparation for electron microscopy

The detailed protocol is described in (Gallet et al. 2024). Briefly, a fraction of sample was centrifuged at 1,000×g for 5 min, and pelleted cells were cryo-fixed using high-pressure freezing (EM HPM100, Leica, Germany) in which cells were subjected to a pressure of 210 MPa at 196□°C, followed by freeze substitution (EM ASF2, Leica, Germany). For the freeze substitution, a mixture 2% (w/v) osmium tetroxide in dried acetone was used. The freeze-substitution system was programmed as follows: 60-80□hours at −90□°C, heating rate of 2□°C hour^−1^ to −60□°C (15□h), 10-12□hours at −60□°C, heating rate of 2□°C□hour^−1^ to −30□°C (15□h), and 10-12□hours at −30□°C, quickly heated to 0□°C for 1 hour to enhance the staining efficiency of osmium tetroxide and then back at −30□°C. The cells were then washed four times in anhydrous acetone for 15□minutes each at −30□°C and gradually embedded in Epon resin (Embed812). A graded resin/acetone (v/v) series was used (30, 50 and 70% resin) with each step lasting 2□hours at increased temperature: 30% resin/acetone bath from −30□°C to −10□°C, 50% resin/acetone bath from −10□°C to 10□°C, 70% resin/acetone bath from 10□°C to 20□°C. Samples were then placed in 100% resin for 8-10□hours and in 100% resin with the accelerator benzyldimethylaminefor 8□hours at room temperature. Resin polymerization finally occurred at 65 °C for 48□hours. Ultrathin sections (70–80 nm) were prepared with a diamond knife on a PowerTome ultramicrotome (RMC products, Tucson, AZ, USA) and collected on 200-mesh copper grids. Samples were visualized by scanning transmission electron microscopy (STEM) using a MERLIN microscope (Zeiss, Oberkochen, Germany) set up at 30 kV and 240 pA.

#### FIB-SEM Imaging

Focused ion beam (FIB) tomography was performed with a Zeiss CrossBeam 550 microscope (Zeiss, Germany), equipped with Fibics Atlas 3D software for tomography described previously (Uwizeye et al. 2021). The resin block containing the cells was fixed on a stub with silver paste, and surface-abraded with a diamond knife in a microtome to obtain a perfectly flat and clean surface. The entire sample was metallized with 4 nm of platinum to avoid charging during the observations. Inside the FIB-SEM, a second platinum layer (1-2 µm) was deposited locally on the analyzed area to mitigate possible curtaining artefacts. The sample was then abraded slice by slice with the Ga^+^ ion beam (generally with a current of 700 nA at 30 kV). Each freshly exposed surface was imaged by scanning electron microscopy (SEM) at 1.5 kV and with a current of ∼1 nA using the in-column EsB backscatter detector. The simultaneous milling and imaging mode was used for better stability, with an hourly automatic correction of focus and astigmatism. For each slice, a thickness of 8 nm was removed, and the SEM images were recorded with a pixel size of 8 nm, providing an isotropic voxel size of 8 × 8 × 8 nm^3^.

#### Image segmentation and metric geometry computations

From the aligned FIB-SEM stack, the region of interest denoting the cell was cropped to remove unnecessary background using the Fiji software (https://imagej.net/Fiji). The 3D segmentation, reconstructions and visualization was achieved using Dragonfly software (www.theobjects.com/dragonfly). The surface areas and volumes were directly calculated from Dragonfly software.

#### Accession numbers

Species and associated Seipin protein sequences used for phylogeny (NCBI/GenBank accession numbers unless otherwise specified) are listed here and can also be found in Supplementary Table 2: *Thalassiosira oceanica* (EJK50087), *Thalassiosira pseudonana* (XP_002286702), *Nitzschia inconspicua* (KAG7356349), *Phaeodactylum tricornutum* (PHATRDRAFT_47296: NCBI, Phatr3_J47296: Ensembl Protist), *Globisporangium splendens* (KAF1316794), *Phytophtora infestans* (KAF4129844), *Microchloropsis gaditana* (Naga_100503g2), *Microchloropsis salina* (TFJ84559), *Aspergillus fumigatus* (KAH1270221), *Blastomyces dermatitis* (XP_045271980), *Puccinia striiformis* (PSHT_04528), *Rhizophagus irregularis* (RhiirA4_399035), *Leptobrachium leishanense* (A0A8C5QNX3: Uniprot), *Xenopus laevis* (XP_041445307.1), *Micrurus lemniscatus* (A0A2D4I9W1: Uniprot), *Varanus komodoensis* (A0A8D2IST9: Uniprot), *Homo sapiens* (NP_001116427), *Canis lupus* (XP_013976703), *Notrodomas monacha* (CAG0914701.1), *Darwinula stevensoni* (CAG0880472.1), *Dufourea novangliae* (KZC07497.1), *Megachile rotundata* (XP_003702285), *Ctenocephalides felis* (XP_026461669), *Drosophila melanogaster* (NP_570012.1), *Crassostrea virginica* (XP_022334537.1), *Octopus vulgaris* (XP_029649550.1), *Caenorhabditis elegans* (NP_741547.1), *Diploscapter pachys* (PAV68820.1), *Ostreococcus lucimarinus* (XP_001416623.1), *Micromonas commoda* (XP_002506104.1), *Chlamydomonas reinhardtii* (XP_042914764), *Volvox reticuliferus* (GIL71655), *Micractinium conductrix* (PSC72173.1), *Chlorella desiccata* (KAG7673487), *Marchantia polymorpha* (P108), *Selaginella moellendorfii* (XP_024544198), *Ceratodon purpureus* (KAG0592613), *Sphagnum fallax* (KAH8935238), *Ananas comosus* (Seipin1: XP_020106670), *Ananas comosus* (Seipin2: XP_020079770), *Zingiber officinalis* (Seipin1: XP_042467429), *Zingiber officinalis* (Seipin2: XP_042380030), *Arabidopsis thaliana* (Seipin1: AED92296.1), *Arabidopsis thaliana* (Seipin2: AEE31126.1), *Arabidopsis thaliana* (Seipin3: AEC08966.1), *Coffea arabica* (Seipin1: XP_027088813.1), *Coffea arabica* (Seipin2: XP_027111988, *Coffea arabica* (Seipin2-like: XP_027060997.1).

## Supplementary Data files

**Supplementary Table 1:** List of protein sequences used for motifs search and phylogeny

**Supplementary Table 2:** Encoding of MEME motifs used for phylogeny and ancestral states determination.

**Supplementary Table 3:** Ancestral states determination

**Supplementary Table 4:** Ancestral states determination raw results

**Supplementary Table 5**: Volumes of WT and KO cells and inside organelles

**Supplementary Figure S1:** Sequence alignment of PtSeipin and Arabidopsis Seipins

**Supplementary Figure S2:** Comparisons of structure predictions of the luminal domains of Seipins from diatoms (*Phaeodactylum tricornutum* and *Thalassiosira pseudonana*) and Eustimatophyceae (*Microchloropsis gaditana* and *Microchloropsis salina*).

**Supplementary Figure S3:** Motifs identified using MEME

**Supplementary Figure S4:** Positioning of the A, B, C and E motifs on the PtSeipin luminal domain

**Supplementary Figure S5**: Heterologous expression in yeast

**Supplementary Figure S6**: PtSeipin-eGFP overexpressing lines

**Supplementary Figure S7**: Laser Scanning Confocal Microscopy (LSCM) observation of the in vivo localization of PtSeipin in Phaeodactylum tricornutum in control culture conditions in stationary phase.

**Supplementary Figure S8**: Generation of PtSeipin KO mutants

**Supplementary Figure S9:** Growth curves of PtSeipin KO mutants

**Supplementary Figure S10**: Observation and quantification of LD in PtSeipin mutants after 8 days of culture in control (CT) culture conditions.

**Supplementary Figure S11**: Observation of Phaeodactylum tricornutum WT after exposure to different light intensities

**Supplementary Figure S12**: Observation of Phaeodactylum tricornutum WT and KO following phosphorus and nitrogen starvation

**Supplementary Figure S13**: Observation and quantification of LD in PtSeipin mutants after 8 days of culture in high light (HL) culture conditions.

**Supplementary Figure S14**: Cell physiology parameters in PtSeipin mutants under high light conditions

**Supplementary Figure S15**: LD numbers and distributions in cells segmented after focused ion beam scanning electron microsopy (FIB-SEM) imaging

**Supplementary Figure S16**: Images extracted from the FIB-SEM stacks

**Supplementary Figure S17:** Glycerolipid profiles of WT and PtSeipin mutants after 8 days of culture in control condition (A) and high light condition (B).

**Supplementary Figure S18:** Lipids profiles of WT and PtSeipin mutants after 4 days of culture in control condition.

**Supplementary Figure S19:** Lipids profiles of WT and PtSeipin mutants after 4 days of culture in high light condition.

**Supplementary Figure S20:** Changes in glycerolipids distribution across all KO lines and all experiments.

**Supplementary Figure S21:** Triacylglycerol (TAG) profiles of WT and PtSeipin mutants after 8 days of culture in control condition (A) and high light condition (B).

**Supplementary Material**: Generation of independent Pt-GFP overexpressing lines

**Supplementary Movie 1**: WT cell 1 stack obtained after focused ion beam scanning electron microsopy (FIB-SEM)

**Supplementary Movie 2**: WT cell 2 stack obtained after focused ion beam scanning electron microsopy (FIB-SEM)

**Supplementary Movie 3**: PtSeipin KO cell 1 stack obtained after focused ion beam scanning electron microsopy (FIB-SEM)

**Supplementary Movie 4**: PtSeipin KO cell 2 stack obtained after focused ion beam scanning electron microsopy (FIB-SEM)

**Supplementary Movie 5**: WT cell 1 segmentation after focused ion beam scanning electron microsopy (FIB-SEM)

**Supplementary Movie 6**: WT cell 2 segmentation after focused ion beam scanning electron microsopy (FIB-SEM)

**Supplementary Movie 7**: PtSeipin KO cell 1 segmentation after focused ion beam scanning electron microsopy (FIB-SEM)

**Supplementary Movie 8**: PtSeipin KO cell 2 segmentation after focused ion beam scanning electron microsopy (FIB-SEM)

## Acknowledgements

D.L.M. was supported by a PhD grant from the CEA (Focus program ECC). D.L.M.,C.A., G.S.L., V.G., M.L., S.R., C.A., M.C., O.B., J.J., A.A., E.M. and J.S. were supported by the French National Research Agency (GRAL Labex ANR-10-LABEX-04, EUR CBS ANR-17- EURE-0003, PEPR Algadvance A-22-PEBB-0002) and IDEX Université Grenoble-Alpes (DefiCO2 Cross-disciplinary Program Grant ANR-15-IDEX-02). D.L.M., M.C. and J.S. were supported by INRAE BAP division (S-PHAEO project). H.H., Y.P. and Y.G. were supported by the National Natural Science Foundation of China (41976119, 91751117). A.A., E.M., H.H. and Y.G. were supported by a CEA-CAS bilateral program. The LIPANG (Lipid analysis in Grenoble) platform is supported by the Rhône-Alpes Region, the fonds FEDER, and GRAL, financed within the University Grenoble Alpes graduate school (Ecoles Universitaires de Recherche) CBH-EUR-GS (ANR-17-EURE-0003). The microscopy facility µLife of IRIG/DBSCI, funded by CEA Nanobio and GRAL LabEX (ANR-10-LABX-49-01), is financed within the University Grenoble Alpes graduate school CBH-EUR-GS (ANR-17- EURE-0003). The IBS/ISBG EM platform is part of the Grenoble Instruct-ERIC center (ISBG; UMS 3518 CNRS-CEA-UGAEMBL) within the Grenoble Partnership for Structural Biology (PSB), supported by FRISBI (ANR-10-INBS-05-02), and GRAL, financed within the University Grenoble Alpes graduate school (Ecoles Universitaires de Recherche) CBH-EUR-GS (ANR-17-EURE-0003). The IBS/ISBG EM facility is supported by the AuvergneRhône-Alpes Region, the Fondation Recherche Medicale (FRM), the fonds FEDER and the GIS-Infrastructures en Biologie Sante et Agronomie (IBISA).The authors wish to thank Marine Froissard for providing the *Ylr404w* yeast mutant line, Guillaume Allorent, Cécile Giustini, Myriam Sassia Hadjadji, Dimitris Petroutsos, Morgane Michaud, Audrée Lebon and Eugenio Reale for technical assistance and Morgane Michaud, Dimitris Petroutsos, Florence Corellou, Fabrice Rébeillé and Denis Falconet for useful discussion on the project.

## Author’s contribution

D.L.M. has performed most experimental work with the technical help of C.Al. F.R. has screened KO lines and F.R. and M.M. contributed to initial characterization of the KO lines. M.A., R.B., C.Am and Y.G. have developed overexpressing lines. H.H., Y.P., J.S., R.B. and C.Am. contributed to confocal imaging. M.C. contributed to microalgae culture and transformation. V.G. and M.L. have provided technical assistance on lipidomics analyses. G.S.L. and P.H.J. have provided technical assistance on STEM and FIB-SEM respectively and G.S.L. has performed computational analyses of the results. O.B. and A.A. have performed phylogenetic analyses. S.R. contributed bioinformatics tools for data analysis. J.J. and E.M. provided guidance in the analysis of lipidomics profiles. J.S. contributed to the design of experiments and their analyses. J.S. and D.L.M. wrote the first draft of the manuscript. All authors have contributed to the writing of the manuscript.

