## Supplementary data for "Functional study of *Phaeodactylum tricornutum* Seipin homolog highlights unique features of lipid droplets biogenesis in diatoms"

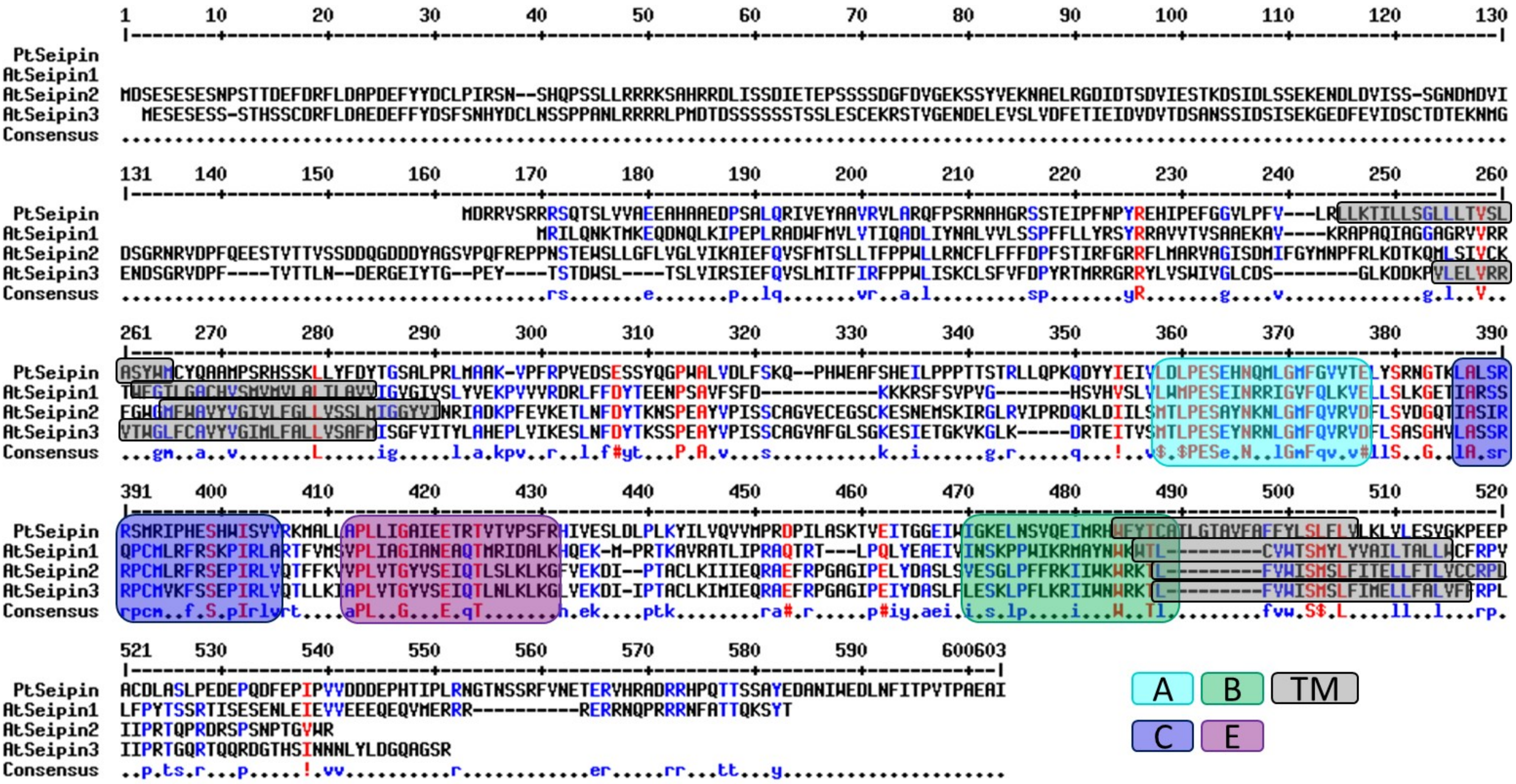


**Supplementary Figure S1: Sequence alignment of PtSeipin and *Arabidopsis* Seipins.**

The four Seipin sequences were retrieved from Uniprot and aligned using Multalin (http://multalin.toulouse.inra.fr/multalin/; (Corpet, 1988)). Conserved motives A, B, C and E identified using MEME (see Supplementary Table2 and Supplementary Figures 3 and 4) are indicated by colored boxes. Predicted transmembrane domains (TM) are indicated by grey boxes.

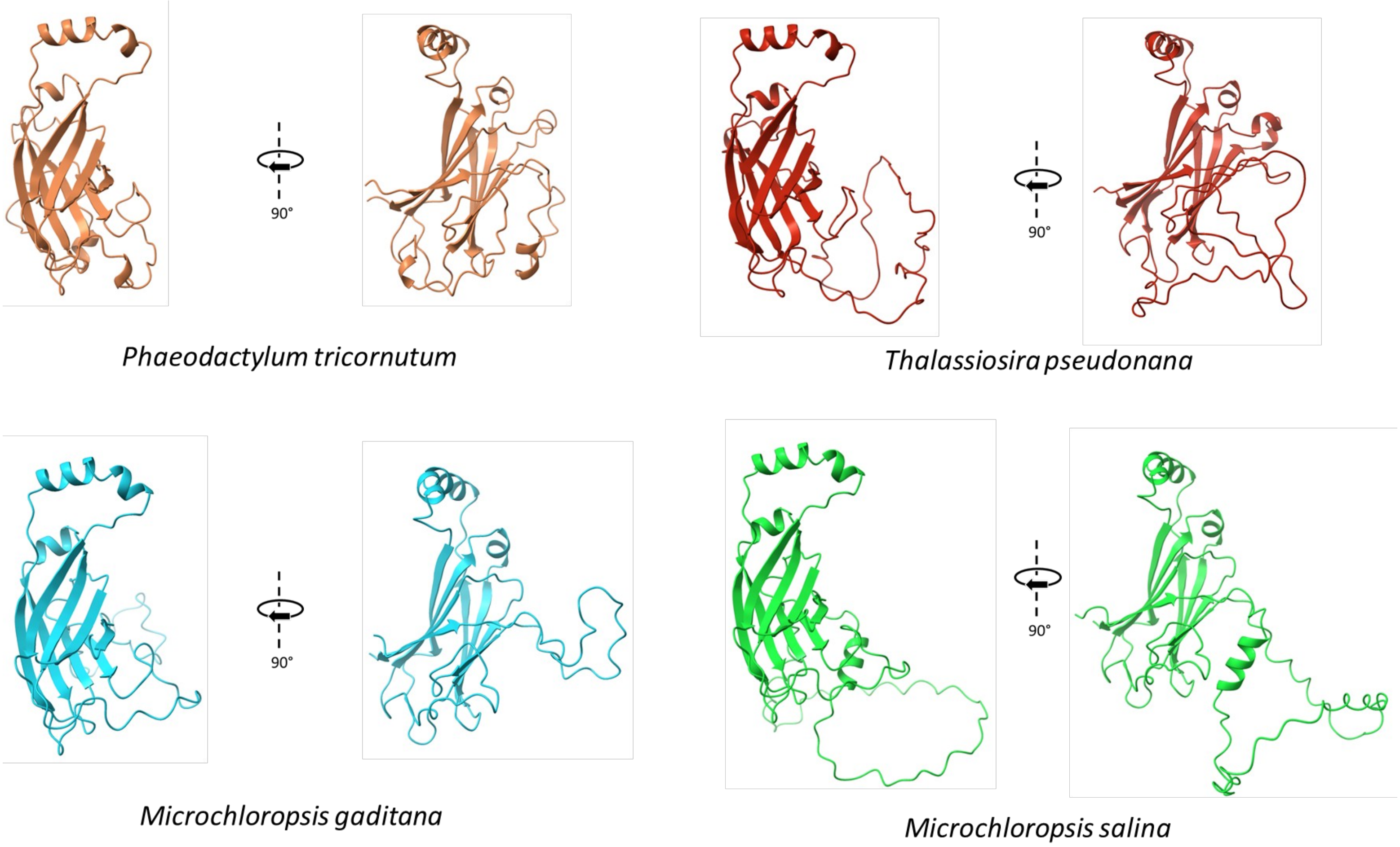


**Supplementary Figure S2: Comparisons of structure predictions of the luminal domains of Seipins from diatoms (*Phaeodactylum tricornutum* and *Thalassiosira pseudonana*) and Eustimatophyceae (*Microchloropsis gaditana* and *Microchloropsis salina*).**
Structure predictions were retrieved from the AlphaFold Database (Varadi et al., 2022) and visualization was performed using ChimeraX (Pettersen et al., 2021).

### Supplementary Figure S3: Motifs identified using MEME

20 motifs (6 to 20 amino acids) have been identified using MEME and a dataset of 48 Seipin sequences.
Each slide corresponds to a motif and the motifs are ranked and named from A to T according to their e-value. For each motif, we show:

- The logo.
- The table containing information on the sequences corresponding to the motifs identified with a significant p-value: species, position in the gene, associated p-value, sequence corresponding to the motif and surrounding regions, repectively in colours and in grey)
- The statistical informations associated with the motif: e-value (estimated number of a particular motif that would appear in a similarly sized set of random sequences), number of species in which the motif has been identified with a significant p-value, width of the motif (between 6 and 20 amino acids). Color code for amino-acids within motifs: Blue: Mostly hydrophobic residues (A, C, F, I, L, V, W and M); Green: Polar, non-charged, non-aliphatic residues (N, Q, S and T); Dark Pink: Acidic residues (D and E); Red: Positively charged residues (K and R); Light Pink: H; Orange: G; Yellow: P; Turquoise: Y.
- The description of the other statistical indicators is given below as provided on the MEME website (https://meme-suite.org/meme/tools/meme).

The log likelihood ratio of the motif. The log likelihood ratio is the logarithm of the ratio of the probability of the occurrences of the motif given the motif model (likelihood given the motif) versus their probability given the background model (likelihood given the null model). (Normally the background model is a 0-order Markov model using the background letter frequencies, but higher order Markov models may be specified via the -bfile option to MEME.).

The information content of the motif in bits. It is equal to the sum of the uncorrected information content, R(), in the columns of the motif. This is equal relative entropy of the motif relative to a uniform background frequency model.

The relative entropy of the motif.

```
re = llr / (sites * ln(2))
```

### Motif A

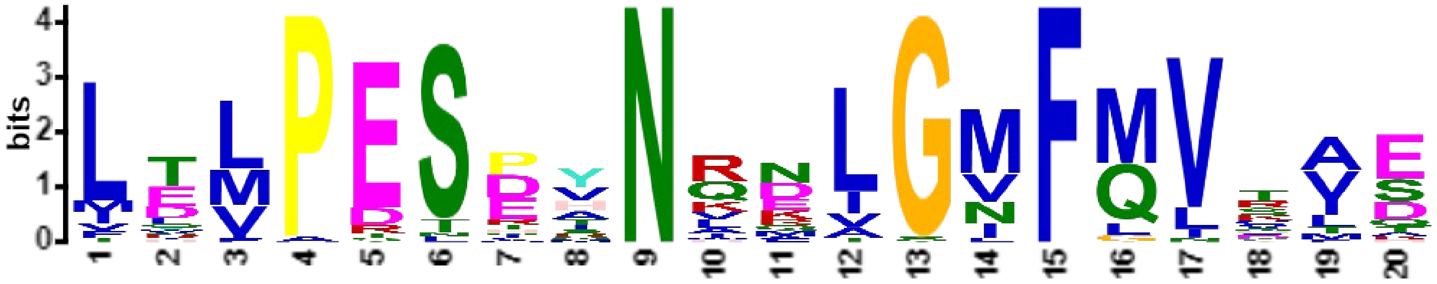

***E*-value:** 2.1e-354
**Site Count:** 43
**Width:** 20

**Log Likelihood Ratio:** 1506
**Information Content:** 53.3
**Relative Entropy:** 50.5
**Bayes Threshold:** 8.8254

| Species | Position in the sequence | P-value | surrounding | Motif associated sequence | surrounding |
|---|---|---|---|---|---|
| 46. A0A6P6S6A8_COFAR | 337 | 4.45e-21 | YNHKLQLTVL | LTMPESEYNRNLGIFQVRVE | CLSSEGKVTA |
| 27. XP_020079770_Seipin-2-like_Ananas | 324 | 5.47e-21 | ANHKLQLTIS | LTLPESDYNRNLGVFQVRVE | FLSAEGKVIF |
| 14. XP_026461669_Ctenocephalides_felis_Fleas | 136 | 6.70e-21 | IGQPYKVYLY | IEMPESDANKNLGMFMVCAE | MRDTASKLRG |
| 25. Q8L615_SEI3_Arabidopsis_thaliana_Eudicot | 326 | 1.47e-20 | LKDRTEITVS | MTLPESEYNRNLGMFQVRVD | FLSASGHVLA |
| 12. XP_003702285_Megachile_rotundata | 112 | 2.15e-20 | VGQPYKVNLH | LEMPESPANRELGMFMVCAQ | LHSRDGFLVE |
| 45. A0A6P6W971_COFAR | 367 | 2.59e-20 | LDHKLQVTVS | LTLPESDYNRNLGIFQVRVD | FLDPDGKAVA |
| 35. PTQ31108_Marchantia_polymorpha_Liverworts | 623 | 4.47e-20 | AGHYFHVNVI | LTMPESEYNIRLGMFQVTAE | MLSARNQVLM |
| 11. A0A154P6K5_Dufourea_novaeangliae | 112 | 4.47e-20 | VGQPYKVNLH | LEMPESPANKELGMFMVCAQ | LHSRDGFLVE |
| 32. KAG0592613_Ceratodon_purpureus_Moss | 421 | 8.99e-20 | PSHKLQVTVF | LTLPESHYNRDLGNFQVSAE | LLSVRGQYLK |
| 16. A0A7R9A2R1_Darwinula_stevensoni | 112 | 1.26e-19 | RGQQYHVLLD | LDMPESPMNQQLGMFMVKID | LKDYDGNVRK |
| 4. A0A8D2IST9_Varanus_komodoensis | 111 | 1.26e-19 | YGQPYRISLE | LELPESPVNQNLGMFMVVIS | CYTKGGRIIS |
| 3. A0A2D4I9W1_Micrurus_lemniscatus_lemniscatus | 111 | 1.26e-19 | YGQPYRMSLE | LELPESPVNQNLGMFMVVIS | CYTKGGRIIS |
| 33. KAH8935238_Sphagnum_fallax_Moss | 442 | 1.75e-19 | KSHKFHVTVF | LTLPESDHNRKLGIFQVSAE | LLSIRGQVIT |
| 15. A0A7R9BHT2_Notodromas_monacha | 110 | 2.43e-19 | RGQPYRVVLS | LDVPESHVNQNLGMFMVHIS | MLNKAGEAVV |
| 28. XP_042380030_Seipin-2_Zingiber_officinale_Monocot | 298 | 6.18e-19 | PNKKLQLTIS | LKLPESDYNRKLGVFQVRVE | LLTSDGKVTS |
| 34. XP_024544198_Selaginella_moellendorffii_Lycophyte | 338 | 8.34e-19 | KTHSFHITAT | LQLPESDHNIRLGMFQVTAE | ILAVNGVVLS |
| 26. F4I340_SEI2_Arabidopsis_thaliana_Eudicot | 355 | 8.34e-19 | RDQKLDIILS | MTLPESAYNKNLGMFQVRVD | FLSVDGQTIA |
| 13. Q9V3X4_Drosophila_melanogaster_Flies | 132 | 1.72e-18 | VGQAYKVIVN | IDMPESPQNLELGMFMVCAE | MRDYDSMLRG |
| 6. XP_013976703_Canis_lupus_familiaris_Carnivors | 174 | 5.90e-18 | YGQPYRVTLE | LELPESPVNQDLGMFLVTIS | CYTRGGRIIS |
| 5. NP_001116427_Homo_sapiens_Primates | 112 | 5.90e-18 | YGQPYRVTLE | LELPESPVNQDLGMFLVTIS | CYTRGGRIIS |
| 8. A0A1L8GJQ5_Xenopus_laevis | 112 | 6.73e-18 | HGQPYRISLE | LQLPESIVNQDLGMFMVTMS | CYTRGGRQIS |
| **1. Phatr3_J47296_Phaeodactylum_tricornutum_Diatom** | **190** | **2.68e-17** | PKQDYYIEIV | LDLPESEHNQMLGMFGVVTE | LYSRNGTKLA |
| 7. A0A8C5QNX3_Leptobrachium_leishanense | 112 | 4.84e-17 | YGQPYRMSLE | LHVPESFVNQDLGMFMVSMS | CYTHGGKEIS |
| 47. EJK50087_Thalassiosira_oceanica_Diatom | 194 | 1.50e-16 | PDYKYFFELS | LTLPESTANRDIGVFMISVE | LQSKDRTVLA |
| 37. XP_002506104.1_Micromonas_commoda_Mamiellales | 136 | 1.68e-16 | PGQRFDVSVS | LTLPETRHNVDAGVFQVRAE | LLTARGEVIA |
| 9. A0A8B8E1X8_Crassostrea_virginica | 110 | 2.32e-16 | RGESYNIKMD | TELPESPINKELGMFMVKLQ | LYDKTGEVVS |
| 30. XP_020106670_Seipin-1_Ananas_comosus_Monocot | 179 | 2.87e-16 | TRHTTHVLLN | MLMPESNHNHKIGIFQVTAE | AIASNGDVIE |
| 29. Q9FFD9_SEI1_Arabidopsis_thaliana_Eudicot | 167 | 3.55e-16 | VGHSVHVSLV | LWMPESEINRRIGVFQLKVE | LLSLKGETIA |
| 44. A0A6P6UED4_COFAR | 190 | 4.85e-16 | VGHTFYVSLV | FLMPESDYNREIGLFQVTAE | VISRNGNIMA |
| 48. KAG7356349_Nitzschia_inconspicua_Diatom | 252 | 1.62e-15 | RNVAHYIEVV | LDLPESEHNRKVGIFGVLVE | LQSNNGTLLA |
| 22. KAF1316794_Globisporangium_splendens_Oomycete | 153 | 1.97e-15 | PGVKYDIYIE | LTVPESRANVDIGVFMVSTT | LKSTDGQYLA |
| 2. XP_002286702_Thalassiosira_pseudonana_Diatom | 192 | 3.49e-15 | QGQHYFLEVV | LILPESEINKQVGVFMLTVD | LLSDKKQLLA |
| 31. XP_042467429_Seipin-1_Zingiber_officinale_Monocot | 161 | 5.07e-15 | PGHSVTVSLL | ILLPESDYNLLIGVFQVQAE | VISSTGEIIA |
| 10. A0A6P7TH74_Octopus_vulgaris | 110 | 1.05e-14 | SGQAYKIMLV | LEIPESTVNQKTGMFLVKAH | FYNQSNHYIQ |
| 23. KAF4129844_Phytophthora_infestans_Oomycete | 150 | 1.26e-14 | PGVTYDVIVE | LTVPESRTNAEVGVFMVSTT | LYSNQERGLA |
| 36. XP_001416623.1_Ostreococcus_lucimarinus_Mamiellale | 114 | 1.50e-14 | AKQNFDVDIE | FVVPDSEYNVNVGMFQVNAK | LLTPSGKTLL |
| 21. TFJ84559_Microchloropsis_salina_Eustigmatophyte | 202 | 3.58e-14 | PERVYVIDVE | VTVPDSETNAMLGNFMLFVD | LFTKDDDLLA |
| 20. Naga_100503g2_Microchloropsis_gaditana_eustigmato | 156 | 3.58e-14 | PERVYVIDVE | VTVPDSETNAMLGNFMLFVD | LFTKDDDLLA |
| 17. A0A2I1GAN6_Rhizophagus_irregularis | 113 | 4.62e-14 | ADQAYDILID | LDVPSSDRNVALGNFMVGLE | LMAKNETVQY |
| 24. A0A2S4WCN4_Puccinia_striiformis | 138 | 5.94e-14 | VFLAYDVHLH | LTVPTNDRNMNLGNFMVHFA | LVTTPPHNTT |
| 18. KAH1270221_Aspergillus_fumigatus_Ascomycota | 111 | 1.47e-12 | SQQAYDVAVK | LEMPRTPSNLAAGNFMIDLS | LFSRPSTSAT |
| 19. XP_045271980_Blastomyces_dermatitidis_Ascomycota | 104 | 5.76e-12 | PAQQYDISVA | LYLPRTPSNLDAGNFMLDLA | LVSTVDTNIN |
| 39. A0A2A2K4L1_Diploscapter_pachys | 102 | 1.77e-10 | RSVSYSLSVR | LNFADLDQNRQLSMFQNQLS | VHDAEGRLLK |

#### Motif B

***E*-value:** 2.1e-217
**Site Count:** 31
**Width:** 20

**Log Likelihood Ratio:** 1087
**Information Content:** 50
**Relative Entropy:** 50.6
**Bayes Threshold:** 10.134

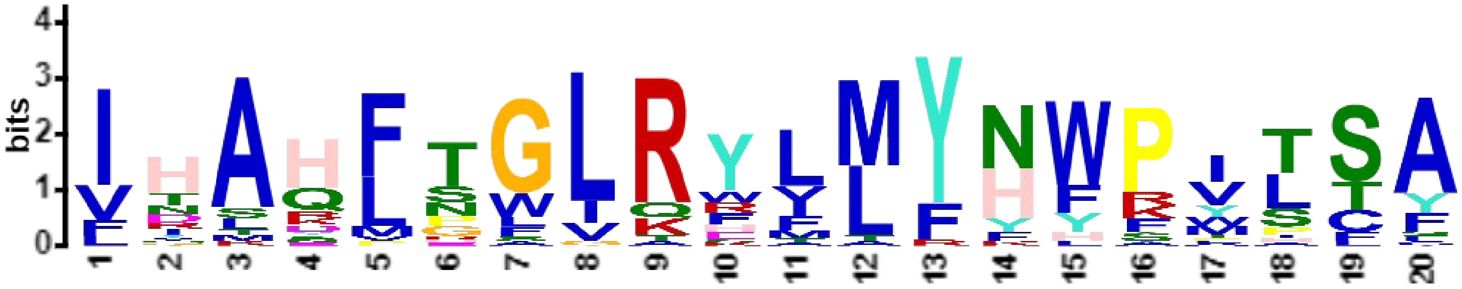


| | | | | | |
|---|---|---|---|---|---|
| 6. XP_013976703_Canis_lupus_familiaris_Carnivors | 278 | 3.26e-23 | RIQMYGAYLR | IHAHFTGLRYLLYNFPMTCA | FIGVASNFTF |
| 5. NP_001116427_Homo_sapiens_Primates | 216 | 3.26e-23 | RIQLYGAYLR | IHAHFTGLRYLLYNFPMTCA | FIGVASNFTF |
| 4. A0A8D2IST9_Varanus_komodoensis | 215 | 4.54e-23 | RIQIYGAYLR | IHAHFTGLRYLLYNFPVTSA | ILGVVSNFAF |
| 3. A0A2D4I9W1_Micrurus_lemniscatus_lemniscatus | 215 | 4.54e-23 | RIQIYGAHLR | IHAHFTGLRYLLYNFPMTSA | ILGVASNFMF |
| 12. XP_003702285_Megachile_rotundata | 216 | 1.06e-21 | HIEFYSATIM | INAHLSGLRYLMYHWPILSA | VVGIGTNLFF |
| 14. XP_026461669_Ctenocephalides_felis_Fleas | 240 | 1.41e-21 | KIEFYSASLH | ITAHFTGLRYIMFHWPILSA | AIGISTNLFF |
| 10. A0A6P7TH74_Octopus_vulgaris | 214 | 2.14e-21 | HVQLYSAVVK | IYAHFTGLRYLMFNWPVSCA | VAGTGIILTV |
| 16. A0A7R9A2R1_Darwinula_stevensoni | 216 | 3.71e-21 | HVQIYGAQLR | VHAHFTGLRYLMFHFPVLSA | VIGIATNLFF |
| 15. A0A7R9BHT2_Notodromas_monacha | 214 | 4.87e-21 | FIEIYSASLH | IHAHFHGLRFLMYHYPVLSA | VCGVSMNMAF |
| 9. A0A8B8E1X8_Crassostrea_virginica | 216 | 7.26e-21 | KIQIYSAVLK | IHAHFTGLRYMLFHWPLLSA | VIGTTLNMCL |
| 7. A0A8C5QNX3_Leptobrachium_leishanense | 216 | 8.29e-21 | RIQIYSAELR | VHAHFTGLRYLLYNYPISTA | IIGVSSNFFF |
| 11. A0A154P6K5_Dufourea_novaeangliae | 216 | 1.23e-20 | HIEFYSATVM | INAHLSGLRYLMFHWPILSA | IVGIGTNLFF |
| 8. A0A1L8GJQ5_Xenopus_laevis | 216 | 3.02e-20 | RIQIYSAELR | VHAYFTGLRYLLYNFPITSA | VIGISSNFIF |
| 13. Q9V3X4_Drosophila_melanogaster_Flies | 236 | 1.38e-18 | KIQFYTVTLH | IVADFTGLRYIMFNWPVLSA | IVAISTNLFF |
| 18. KAH1270221_Aspergillus_fumigatus_Ascomycota | 229 | 6.50e-18 | PLQVYSAQVE | FHARFTGLRWVMYNWRILSF | LVFSFGFWSV |
| 23. KAF4129844_Phytophthora_infestans_Oomycete | 254 | 3.52e-17 | KLQVYSAKLT | VIAQLTGLRYLMYHWAVPTA | ILFILNIVFL |
| 21. TFJ84559_Microchloropsis_salina_Eustigmatophyte | 306 | 5.86e-17 | RLHLYKASMS | ITAQLNWLQFVMYNWFYTTA | FLIVLLISSI |
| 20. Naga_100503g2_Microchloropsis_gaditana_eustigmato | 260 | 5.86e-17 | RLHLYKASMS | ITAQLNWLQFVMYNWFYTTA | FLIILLISSI |
| 22. KAF1316794_Globisporangium_splendens_Oomycete | 257 | 1.07e-16 | AIQIYSAKLT | IIAQLSGVRYLMYHWSVSTA | VLVILNIVFL |
| 19. XP_045271980_Blastomyces_dermatitidis_Ascomycota | 224 | 2.14e-16 | RMQVYNAVVR | FDAKFSGLRWIMYNWKILSF | LTFSSTFWIV |
| 17. A0A2I1GAN6_Rhizophagus_irregularis | 216 | 3.96e-15 | NLDVYNAQIR | LDAHFRGLRYFMYYHSIPTA | ITFMSMFLTW |
| 38. Q8MXG1_Caenorhabditis_elegans | 189 | 5.90e-14 | FANIEEAELI | VTARFGLIRHLLYYWPTTSY | ATIFVSTFVI |
| 39. A0A2A2K4L1_Diploscapter_pachys | 205 | 8.95e-14 | YAQVDSAYLV | IHANFGLIRHFTYFWPVSFY | LLIFIPTFSF |
| 24. A0A2S4WCN4_Puccinia_striiformis | 266 | 1.15e-13 | ELQIYKAVLV | FDAHLEGLRWALYYHPYISF | CFFSAFFFTA |
| 30. XP_020106670_Seipin-1_Ananas_comosus_Monocot | 286 | 1.35e-12 | LPQLYEADIV | IRTQLPWGKEFMYNWRWTFY | VWTSFYMYIV |
| 34. XP_024544198_Selaginella_moellendorffii_Lycophyte | 451 | 2.31e-12 | LPEIYNAEVL | VQSQLPWLKSVLYKWKWTCY | VWGAVLIFVL |
| 29. Q9FFD9_SEI1_Arabidopsis_thaliana_Eudicot | 274 | 3.65e-11 | LPQLYEAEIV | INSKPPWIKRMAYNWKWTLC | VWTSMYLYVA |
| **1. Phatr3_J47296_Phaeodactylum_tricornutum_Diatom** | **302** | **5.54e-11** | TVEITGGEIW | IGKELNSVQEIMRHWFYTCA | TLGTAVFAFF |
| 36. XP_001416623.1_Ostreococcus_lucimarinus_Mamiellale | 223 | 7.82e-10 | LPQIYEARAI | VHLSMNFVAKLLYFYPIASS | FVLVGLFWSG |
| 41. KAG7673487_Chlorella_desiccata_Trebouxiophyceae | 490 | 1.07e-9 | SPKVLSASLR | IHLQVGFIRRTLYHLRPHSL | IALAMGAGAI |
| 37. XP_002506104.1_Micromonas_commoda_Mamiellales | 247 | 1.14e-9 | VPQMWRAHAD | LRMDMSALTRFLYHYPASSF | AVMVLVTWGY |

### Motif C

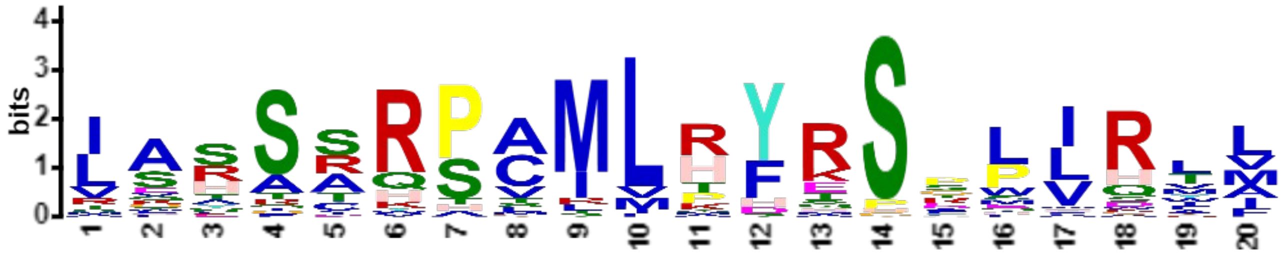


***E*-value:** 1.1e-205
**Site Count:** 43
**Width:** 20

**Log Likelihood Ratio:** 1183
**Information Content:** 43.5
**Relative Entropy:** 39.7
**Bayes Threshold:** 8.8254

| | | | | | |
|---|---|---|---|---|---|
| 29. Q9FFD9_SEI1_Arabidopsis_thaliana_Eudicot | 195 | 8.74e-19 | VELLSLKGET | IARSSQPCMLRFRSKPIRLA | RTFVMSVPLI |
| 44. A0A6P6UED4_COFAR | 218 | 3.25e-18 | AEVISRNGNI | MARSSHPCMLRFRSWPIRTM | QTFLMGLPLL |
| 31. XP_042467429_Seipin-1_Zingiber_officinale_Monocot | 189 | 3.25e-18 | AEVISSTGEI | IAASSRPCMLRFRSFPVRLM | RTLFMSIPLL |
| 26. F4I340_SEI2_Arabidopsis_thaliana_Eudicot | 383 | 6.79e-17 | VDFLSVDGQT | IASIRRPCMLRFRSEPIRLV | QTFFKVVPLV |
| 45. A0A6P6W971_COFAR | 395 | 2.02e-16 | VDFLDPDGKA | VASSRHPCMLPFKSRPIRLL | LTFLKVAPLL |
| 13. Q9V3X4_Drosophila_melanogaster_Flies | 160 | 2.02e-16 | AEMRDYDSML | RGHSCRSAMMRYRSPLIRMI | STWVLSPLYV |
| 8. A0A1L8GJQ5_Xenopus_laevis | 140 | 2.55e-16 | MSCYTRGGRQ | ISYTARSAMLHYKSPLLRTM | ETMASSPLLL |
| 4. A0A8D2IST9_Varanus_komodoensis | 139 | 2.87e-16 | ISCYTKGGRI | ISSSARSAMLHYRSGLLQIL | DTLAFASLFL |
| 30. XP_020106670_Seipin-1_Ananas_comosus_Monocot | 207 | 3.61e-16 | AEAIASNGDV | IEASSQPCMLRYRSLPVRLM | RTMLMGVPLL |
| 21. TFJ84559_Microchloropsis_salina_Eustigmatophyte | 230 | 5.71e-16 | VDLFTKDDDL | LAHSARPLILTYRSPWIRLF | RSLLLAPLLA |
| 3. A0A2D4I9W1_Micrurus_lemniscatus_lemniscatus | 139 | 7.15e-16 | ISCYTKGGRI | ISSSARAAMLHYRSGLLQML | DTLAFSGLFL |
| 7. A0A8C5QNX3_Leptobrachium_leishanense | 140 | 8.93e-16 | MSCYTHGGKE | ISHTARSAILHYKSPLLRTL | ETFAFLPLLL |
| 20. Naga_100503g2_Microchloropsis_gaditana_eustigmato | 184 | 9.98e-16 | VDLFTKDDDL | LAHSARPLILTYRSPWIRFF | RSLLLAPVLA |
| 6. XP_013976703_Canis_lupus_familiaris_Carnivors | 202 | 1.55e-15 | ISCYTRGGRI | ISTSSRSVMLHYRSDLLQML | DTLVFSSLLL |
| 5. NP_001116427_Homo_sapiens_Primates | 140 | 1.55e-15 | ISCYTRGGRI | ISTSSRSVMLHYRSDLLQML | DTLVFSSLLL |
| 34. XP_024544198_Selaginella_moellendorffii_Lycophyte | 366 | 2.94e-15 | AEILAVNGVV | LSRFTHPCMLRYRSRPIRYA | KTAMLAVPLI |
| 12. XP_003702285_Megachile_rotundata | 140 | 5.51e-15 | AQLHSRDGFL | VEHACRSAMLHYRSTLLHAL | TTLTFSPMMI |
| 14. XP_026461669_Ctenocephalides_felis_Fleas | 164 | 6.10e-15 | AEMRDTASKL | RGHSCRSAMLHYRSDLLHSL | NTLALSPLLI |
| 25. Q8L615_SEI3_Arabidopsis_thaliana_Eudicot | 354 | 7.49e-15 | VDFLSASGHV | LASSRRPCMVKFSSEPIRLV | QTLLKIAPLV |
| 33. KAH8935238_Sphagnum_fallax_Moss | 470 | 1.01e-14 | AELLSIRGQV | ITRATRPCMLKFHSSPIRYA | KNLLMGVPFL |
| 11. A0A154P6K5_Dufourea_novaeangliae | 140 | 2.47e-14 | AQLHSRDGFL | VEHACRSAMMHYRSTLLHAL | TTFTFSPMMI |
| 35. PTQ31108_Marchantia_polymorpha_Liverworts | 651 | 3.63e-14 | AEMLSARNQV | LMRQSKPCMLRFRSAPIRLF | KTVLYSIPLL |
| 27. XP_020079770_Seipin-2-like_Ananas | 352 | 4.00e-14 | VEFLSAEGKV | IFSSRQPCMLRFKSSHMHII | ETFLKSGSLL |
| 19. XP_045271980_Blastomyces_dermatitidis_Ascomycota | 145 | 1.34e-13 | NAEASAEENI | IARSRRPAILTYASPMVDTA | RRVSKMPLYV |
| 18. KAH1270221_Aspergillus_fumigatus_Ascomycota | 150 | 1.76e-13 | TTGQNTSSNR | IVHSRRPAILTYTSPMVDTA | SKISFMPLYV |
| 17. A0A2I1GAN6_Rhizophagus_irregularis | 140 | 3.28e-13 | GLELMAKNET | VQYSSRPCILTYQSGLFRVI | YTFWRLIPLV |
| 32. KAG0592613_Ceratodon_purpureus_Moss | 449 | 3.58e-13 | AELLSVRGQY | LKRASWPCMLRFQSSSIRYA | KQVMLGVPLL |
| 15. A0A7R9BHT2_Notodromas_monacha | 138 | 3.91e-13 | ISMLNKAGEA | VVKSDRSAMLRYRSGLLRAI | YTFCYAPMLV |
| 10. A0A6P7TH74_Octopus_vulgaris | 138 | 4.65e-13 | AHFYNQSNHY | IQSAAKTTMLRYRSSLLHTM | STLFYLPFLI |
| 28. XP_042380030_Seipin-2_Zingiber_officinale_Monocot | 326 | 9.23e-13 | VELLTSDGKV | TSSSRHPCLLRFKSSQIHFL | QTFLKSLFLL |
| 47. EJK50087_Thalassiosira_oceanica_Diatom | 222 | 1.95e-12 | VELQSKDRTV | LARSRQHSMLPYESPLVSTF | RKITLLAPLM |
| 46. A0A6P6S6A8_COFAR | 365 | 1.95e-12 | VECLSSEGKV | TASSSYPTMLQFKSQPIRFV | GTIFKSPLLL |
| 22. KAF1316794_Globisporangium_splendens_Oomycete | 181 | 6.52e-12 | TTLKSTDGQY | LASSARPAIVHDSHSLVRWI | RVGALAISHA |
| 9. A0A8B8E1X8_Crassostrea_virginica | 140 | 1.41e-11 | LYDKTGEVVS | SSSRSVSVMLHYRSELLRIM | DTFVFSPLLL |
| 16. A0A7R9A2R1_Darwinula_stevensoni | 140 | 2.06e-11 | IDLKDYDGNV | RKSGTRAAMLRYRSFLHQLV | YTLAFAPVLL |
| 37. XP_002506104.1_Micromonas_commoda_Mamiellales | 164 | 1.18e-10 | AELLTARGEV | IANATRPAMLAHTGTEVRLL | RLLVSWPLHA |
| 23. KAF4129844_Phytophthora_infestans_Oomycete | 178 | 1.27e-10 | TTLYSNQERG | LAASARPVTLHDMPASVRWM | RLAFWLMPYA |
| 2. XP_002286702_Thalassiosira_pseudonana_Diatom | 220 | 4.09e-10 | VDLLSDKKQL | LATSIQSSMFPYESRLVGTL | RKMTVLLPLV |
| 48. KAG7356349_Nitzschia_inconspicua_Diatom | 280 | 7.44e-10 | VELQSNNGTL | LASSLRTARMPHESKWIAVV | RKAVCIVPLL |
| **1. Phatr3_J47296_Phaeodactylum_tricornutum_Diatom** | **218** | **3.66e-9** | TELYSRNGTK | LALSRRSMRIPHESHWISVV | RKMALLAPLL |
| 24. A0A2S4WCN4_Puccinia_striiformis | 168 | 9.64e-9 | LVTTPPHNTT | LHQSSRHASLMYEPGFARVL | HAFKYLRTLF |
| 41. KAG7673487_Chlorella_desiccata_Trebouxiophyceae | 414 | 1.30e-8 | AELKSVDNRS | AARSTQPVLLRSHSAILWRV | LSAPLHWTGL |
| 36. XP_001416623.1_Ostreococcus_lucimarinus_Mamiellale | 142 | 1.30e-8 | AKLLTPSGKT | LLERSRPGIVKYTSKEVKWL | KTIVWWPFHA |

### Motif D

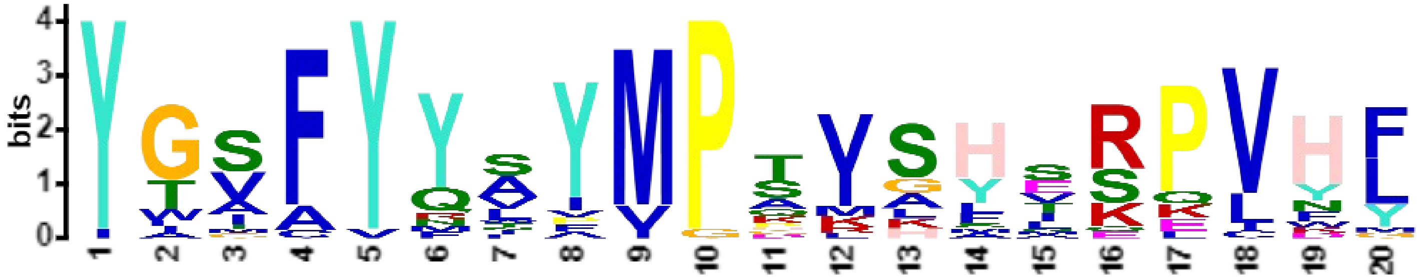

***E*-value:** 1.7e-164
**Site Count:** 23
**Width:** 20

**Log Likelihood Ratio:** 865
**Information Content:** 53.5
**Relative Entropy:** 54.2
**Bayes Threshold:** 10.261

| | | | | | |
|---|---|---|---|---|---|
| 6. XP_013976703_Canis_lupus_familiaris_Carnivors | 111 | 3.38e-23 | LLLLWVSVFL | YGSFYYSYMPTVSHLSPVHF | YYRTDCDSST |
| 5. NP_001116427_Homo_sapiens_Primates | 49 | 3.38e-23 | LLLLWVSVFL | YGSFYYSYMPTVSHLSPVHF | YYRTDCDSST |
| 16. A0A7R9A2R1_Darwinula_stevensoni | 51 | 1.54e-22 | TVLIWLSVFL | YGVFYYLYMPAVAHTRPVHF | IFEPCKEEAG |
| 4. A0A8D2IST9_Varanus_komodoensis | 48 | 2.83e-22 | LLLLWISIFL | YGSFYYSYMPTVSYVSPVHY | HFRTDCGLPG |
| 3. A0A2D4I9W1_Micrurus_lemniscatus_lemniscatus | 48 | 2.83e-22 | LLLLWISIFL | YGSFYYSYMPTVSYVSPVHY | QFRTDCGQPG |
| 8. A0A1L8GJQ5_Xenopus_laevis | 49 | 6.73e-22 | LLLLWVSVFL | YGSFYYSYMPTVKYSSPVHY | QYSSTCEPPP |
| 10. A0A6P7TH74_Octopus_vulgaris | 49 | 5.68e-21 | IALLWIALFV | YGSFYYVYMPSVSHERPCNF | VFDVCDNGVG |
| 14. XP_026461669_Ctenocephalides_felis_Fleas | 77 | 8.28e-21 | SIIVWLAIFM | YIVFYYTYMPSISHIRPVHL | QFKSCEEGQA |
| 12. XP_003702285_Megachile_rotundata | 53 | 8.28e-21 | VFIVWLSVFL | YTAFYYAYMPSMSYIRPVHL | QFKSCNEQRG |
| 11. A0A154P6K5_Dufourea_novaeangliae | 53 | 8.28e-21 | VLIVWLSVFL | YTAFYYAYMPSMSYIRPVHL | QFKSCNEQKG |
| 7. A0A8C5QNX3_Leptobrachium_leishanense | 49 | 2.48e-20 | IFLLWVSVFL | YGSFYYSYMPAVKFSSPVHY | QYSSFCDPPP |
| 15. A0A7R9BHT2_Notodromas_monacha | 49 | 2.79e-20 | TVMVWMSVFM | YGVFYYVYMPDVSHMRPVDF | KFKPCGEKPG |
| 13. Q9V3X4_Drosophila_melanogaster_Flies | 73 | 5.63e-20 | VLIIWLAVFM | YAAFYYVYMPAISHTRPVHM | QFKTCLETST |
| 9. A0A8B8E1X8_Crassostrea_virginica | 49 | 1.71e-17 | VILIWFSAFL | YGSFYFAYMPPVSLTRPVNL | QFKICEDGIG |
| 19. XP_045271980_Blastomyces_dermatitidis_Ascomycota | 56 | 6.27e-17 | LVLFCISVVA | YWIFYYNYVPQIGMERQVHL | QFGDGHPYGT |
| 18. KAH1270221_Aspergillus_fumigatus_Ascomycota | 63 | 6.74e-16 | VCMFFVSSFA | YGIFYYWFIPQIGLVRQVHL | QFGDDQPWGT |
| 17. A0A2I1GAN6_Rhizophagus_irregularis | 58 | 1.49e-14 | FILISVAFVS | YLGFYMIYVPKIAHAKPVYL | QYHKDDSPYA |
| 21. TFJ84559_Microchloropsis_salina_Eustigmatophyte | 63 | 1.79e-13 | LALITSATAL | YTVAYQLIMPTKLHEKELFF | DYFPPANPLG |
| 20. Naga_100503g2_Microchloropsis_gaditana_eustigmato | 63 | 1.79e-13 | LALITSATAL | YTVAYQLIMPTKLHEKELFF | DYFPPANPLD |
| 24. A0A2S4WCN4_Puccinia_striiformis | 96 | 2.07e-13 | VISLLLSIIA | YTTFYRAYVPHVGFSQPIWL | QYGYVHLPSS |
| 48. KAG7356349_Nitzschia_inconspicua_Diatom | 93 | 2.42e-12 | TILLCSSLCL | YGVFYNAVMPGLHASEKLYF | DYEGMAQRPR |
| **1. Phatr3_J47296_Phaeodactylum_tricornutum_Diatom** | **98** | **5.06e-12** | GLLLTVSLAS | YWMCYQAAMPSRHSSKLLYF | DYTGSALPRL |
| 47. EJK50087_Thalassiosira_oceanica_Diatom | 421 | 4.30e-10 | VTDEDSVTHG | IGSAVQFPMGKIAHNRKVKG | HGSPFAAKET |

### Motif E

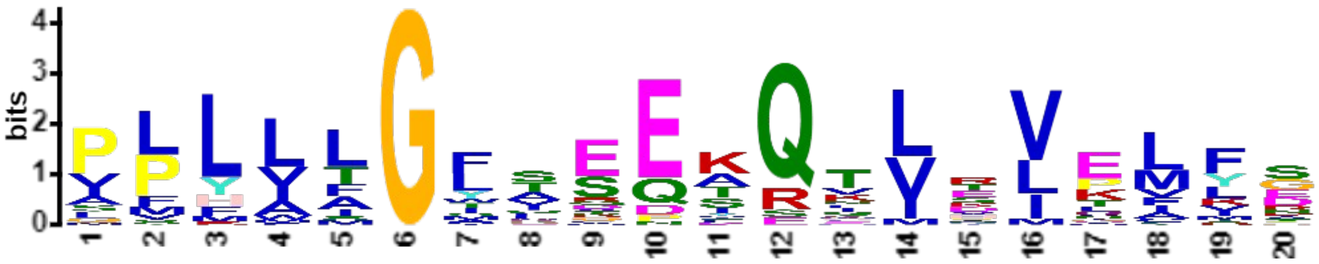

***E*-value:** 2.7e-149
**Site Count:** 44
**Width:** 20

**Log Likelihood Ratio:** 1062
**Information Content:** 40.4
**Relative Entropy:** 34.8
**Bayes Threshold:** 9.2765

| | | | | | |
|---|---|---|---|---|---|
| 10.A0A6P7TH74_Octopus_vulgaris | 164 | 6.10e-15 | LHTMSTLFYL | PFLITGANEEKQILHVELFS | KYTEDPYNRA |
| 7.A0A8C5QNX3_Leptobrachium_leishanense | 166 | 7.13e-15 | LRTLETFAFL | PLLLFGMSEQKQSLEVELYS | AYREDSYVPT |
| 8.A0A1L8GJQ5_Xenopus_laevis | 166 | 8.31e-15 | LRTMETMASS | PLLLLGLSDQKQILEVELYS | EYREDSYVPT |
| 3.A0A2D4I9W1_Micrurus_lemniscatus_lemniscatus | 165 | 8.31e-15 | LQMLDTLAFS | GLFLAGFTEQKQTVDVELYS | DYKEDSYIPT |
| 14.XP_026461669_Ctenocephalides_felis_Fleas | 190 | 2.35e-14 | LHSLNTLALS | PLLIIGTAEQKQIVPVELFS | NYEEDQNYPV |
| 11.A0A154P6K5_Dufourea_novaeangliae | 166 | 2.35e-14 | LHALTTFTFS | PMMIFGTTEEKQNVVLELFG | NFEEDQSHPV |
| 6.XP_013976703_Canis_lupus_familiaris_Carnivors | 228 | 3.13e-14 | LQMLDTLVFS | SLLLFGFAEQKQLLEVELYS | EYRENSYVPT |
| 21.TFJ84559_Microchloropsis_salina_Eustigmatophyte | 256 | 3.60e-14 | IRLFRSLLLA | PLLALGFAQEAQVLRVTFFD | RFRESLAHPL |
| 9.A0A8B8E1X8_Crassostrea_virginica | 166 | 5.48e-14 | LRIMDTFVFS | PLLLSGFVEQKQFLTVEFYN | NYIDDSYNPA |
| 12.XP_003702285_Megachile_rotundata | 166 | 1.08e-13 | LHALTTLTFS | PMMIFGSTEEKQNVVLELFG | NFEEDQSHPV |
| 4.A0A8D2IST9_Varanus_komodoensis | 165 | 1.40e-13 | LQILDTLAFA | SLFLTGFTEQKQMVEIELYS | DYKEDSYTPT |
| 5.NP_001116427_Homo_sapiens_Primates | 166 | 1.82e-13 | LQMLDTLVFS | SLLLFGFAEQKQLLEVELYA | DYRENSYVPT |
| 16.A0A7R9A2R1_Darwinula_stevensoni | 166 | 3.43e-13 | HQLVYTLAFA | PVLLTGTFEEKQLVTVELLS | LFEDDPNDPV |
| 20.Naga_100503g2_Microchloropsis_gaditana_eustigmato | 210 | 3.88e-13 | IRFFRSLLLA | PVLALGFAQEAQVLRVTFFD | RFRESLAHPL |
| 13.Q9V3X4_Drosophila_melanogaster_Flies | 186 | 3.88e-13 | IRMISTWVLS | PLYVLGWKEEFQQVPVEIFS | RYLEERQHPI |
| 15.A0A7R9BHT2_Notodromas_monacha | 164 | 4.39e-13 | LRAIYTFCYA | PMLVTGSWEEKQTVDIELVP | NFNENAHDPT |
| 26.F4I340_SEI2_Arabidopsis_thaliana_Eudicot | 409 | 5.60e-13 | IRLVQTFFKV | VPLVTGYVSEIQTLSLKLKG | FVEKDIPTAC |
| 17.A0A2I1GAN6_Rhizophagus_irregularis | 166 | 9.04e-13 | FRVIYTFWRL | IPLVLGFTKEDQQLKVVMFE | NMIESAEKPI |
| 25.Q8L615_SEI3_Arabidopsis_thaliana_Eudicot | 380 | 1.14e-12 | IRLVQTLLKI | APLVTGYVSEIQTLNLKLKG | LVEKDIIPTA |
| 18.KAH1270221_Aspergillus_fumigatus_Ascomycota | 176 | 2.53e-12 | VDTASKISFM | PLYVLGWQREAETLRVPMLE | KIEFSRGWRN |
| 37.XP_002506104.1_Micromonas_commoda_Mamiellales | 190 | 3.52e-12 | VRLLRLLVSW | PLHALGLVEEVRTVTLPMFE | GVGERLTRPF |
| 46.A0A6P6S6A8_COFAR | 391 | 3.93e-12 | IRFVGTIFKS | PLLLAGFQSEIQHLKIGVMD | FTEGYEPTAC |
| 23.KAF4129844_Phytophthora_infestans_Oomycete | 204 | 4.38e-12 | VRWMRLAFWL | MPYALGFTEPAQTLRVTAIN | GYQESAEYPL |
| 35.PTQ31108_Marchantia_polymorpha_Liverworts | 677 | 7.45e-12 | IRLFKTVLYS | IPLLMGFSTETQKIEVTVLH | AQEKLVPTAS |
| 19.XP_045271980_Blastomyces_dermatitidis_Ascomycota | 171 | 7.45e-12 | VDTARRVSKM | PLYVLGWQREAEKLKVNMMG | RVEFARKKGA |
| 45.A0A6P6W971_COFAR | 421 | 8.27e-12 | IRLLLTFLKV | APLLTGYTSESQDLIIKFKG | FTEGGRPTSC |
| 36.XP_001416623.1_Ostreococcus_lucimarinus_Mamiellale | 168 | 9.17e-12 | VKWLKTIVWW | PFHALGLIEEQQNVRVAMVQ | NYKEDSESPF |
| 44.A0A6P6UED4_COFAR | 244 | 1.25e-11 | IRTMQTFLMG | LPLLLGIRAETQKVTVPMLK | HKEDFPRTEA |
| 32.KAG0592613_Ceratodon_purpureus_Moss | 475 | 1.53e-11 | IRYAKQVMLG | VPLLMGLSGESQTISLRLFE | NEEETKIPTA |
| 28.XP_042380030_Seipin-2_Zingiber_officinale_Monocot | 352 | 1.53e-11 | IHFLQTFLKS | LFLLAGYSDESQVLKLPMKG | LTEGTKPTTC |
| 30.XP_020106670_Seipin-1_Ananas_comosus_Monocot | 233 | 1.86e-11 | VRLMRTMLMG | VPLLAGVSSETQKITMEILR | YKERQPKTEL |
| 27.XP_020079770_Seipin-2-like_Ananas | 378 | 2.76e-11 | MHIIETFLKS | GSLLAGYSSESQTLRLKMRG | FTEKPEPTAC |
| **1.Phatr3_J47296_Phaeodactylum_tricornutum_Diatom** | **244** | **4.46e-11** | ISVVRKMALL | APLLIGAIEETRTVTVPSFR | HIVESLDLPL |
| 31.XP_042467429_Seipin-1_Zingiber_officinale_Monocot | 215 | 6.48e-11 | VRLMRTLFMS | IPLLLGISSETQEMSMEVLR | YKESRAKRSG |
| 34.XP_024544198_Selaginella_moellendorffii_Lycophyte | 392 | 8.54e-11 | IRYAKTAMLA | VPLILGLEVESQTLVLKLVE | AYEERSSSSY |
| 29.Q9FFD9_SEI1_Arabidopsis_thaliana_Eudicot | 221 | 1.12e-10 | IRLARTFVMS | VPLIAGIANEAQTMRIDALK | HQEKMPRTKA |
| 41.KAG7673487_Chlorella_desiccata_Trebouxiophyceae | 437 | 1.34e-10 | SAILWRVLSA | PLHWTGLLSDARVVSVKVFS | NYIERRDIPF |
| 33.KAH8935238_Sphagnum_fallax_Moss | 496 | 1.34e-10 | IRYAKNLLMG | VPFLIGLSSESQKLAIRIIE | NHEDTSVPTV |
| 47.EJK50087_Thalassiosira_oceanica_Diatom | 248 | 5.72e-10 | VSTFRKITLL | APLMSGLLSETRTVHLICFD | SYVEIDTQRP |
| 2.XP_002286702_Thalassiosira_pseudonana_Diatom | 246 | 7.92e-10 | VGTLRKMTVL | LPLVFGMISETRAISLLCFD | NYVDANIERS |
| 39.A0A2A2K4L1_Diploscapter_pachys | 155 | 1.38e-9 | VTRLTCLAFL | PLYLLGFICNYSTIDILLTD | SHIESLERAS |
| 22.KAF1316794_Globisporangium_splendens_Oomycete | 207 | 1.61e-9 | VRWIRVGALA | ISHALGLTEPAQVIDVLAIN | GITEGKTHPL |
| 40.A0A2P6VDK0_Micractinium_conductrix | 206 | 2.03e-9 | WWSLSSLLRA | PLRWLGLAEDLRRLHVPLFT | GYREKWEVPF |
| 38.Q8MXG1_Caenorhabditis_elegans | 139 | 2.32e-8 | IQKAMHLFLF | PFYFLGFFHDYSTLAIPMSA | DYLEGIDSPS |

### Motif F

***E*-value:** 1.7e-102
**Site Count:** 18
**Width:** 20

**Log Likelihood Ratio:** 668
**Information Content:** 55.7
**Relative Entropy:** 53.6
**Bayes Threshold:** 10.3464

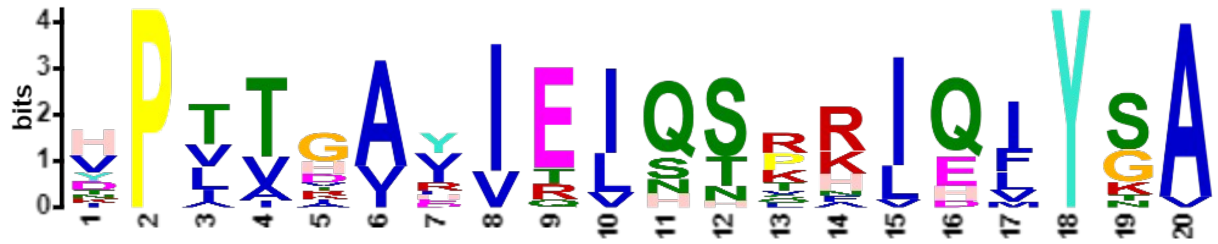


| | | | | | |
|---|---|---|---|---|---|
| 7. A0A8C5QNX3_Leptobrachium_leishanense | 193 | 5.57e-21 | LYSAYREDSY | VPTTGAVIEIQSLRIQIYSA | ELRVHAHFTG |
| 6. XP_013976703_Canis_lupus_familiaris_Carnivors | 255 | 1.84e-20 | LYSEYRENSY | VPTTGAIIEIHSKRIQMYGA | YLRIHAHFTG |
| 5. NP_001116427_Homo_sapiens_Primates | 193 | 3.27e-20 | LYADYRENSY | VPTTGAIIEIHSKRIQLYGA | YLRIHAHFTG |
| 4. A0A8D2IST9_Varanus_komodoensis | 192 | 9.89e-20 | LYSDYKEDSY | TPTVGAVIEIQTTRIQIYGA | YLRIHAHFTG |
| 11. A0A154P6K5_Dufourea_novaeangliae | 193 | 1.13e-19 | LFGNFEEDQS | HPVTVIYIEIQSRHIEFYSA | TVMINAHLSG |
| 3. A0A2D4I9W1_Micrurus_lemniscatus_lemniscatus | 192 | 2.83e-19 | LYSDYKEDSY | IPTIGAVIEIQTTRIQIYGA | HLRIHAHFTG |
| 8. A0A1L8GJQ5_Xenopus_laevis | 193 | 3.22e-19 | LYSEYREDSY | VPTVGAVIQIQSVRIQIYSA | ELRVHAYFTG |
| 12. XP_003702285_Megachile_rotundata | 193 | 4.15e-19 | LFGNFEEDQS | HPVTIIHIEIQSRHIEFYSA | TIMINAHLSG |
| 16. A0A7R9A2R1_Darwinula_stevensoni | 193 | 6.05e-19 | LLSLFEDDPN | DPVTHAIIEVQSRHVQIYGA | QLRVHAHFTG |
| 9. A0A8B8E1X8_Crassostrea_virginica | 193 | 1.60e-18 | FYNNYIDDSY | NPAVGAYIEVQNKKIQIYSA | VLKIHAHFTG |
| 14. XP_026461669_Ctenocephalides_felis_Fleas | 217 | 2.89e-18 | LFSNYEEDQN | YPVTDVYVEIQSRKIEFYSA | SLHITAHFTG |
| 13. Q9V3X4_Drosophila_melanogaster_Flies | 213 | 2.39e-17 | IFSRYLEERQ | HPITDVYVEIQSQKIQFYTV | TLHIVADFTG |
| 15. A0A7R9BHT2_Notodromas_monacha | 191 | 6.14e-17 | LVPNFNENAH | DPTVAAYIELQTRFIEIYSA | SLHIHAHFHG |
| 22. KAF1316794_Globisporangium_splendens_Oomycete | 234 | 1.96e-15 | AINGITEGKT | HPLTTAEITLNHPAIQIYSA | KLTIIAQLSG |
| 23. KAF4129844_Phytophthora_infestans_Oomycete | 231 | 1.04e-14 | AINGYQESAE | YPLTRVDIELNTPKLQVYSA | KLTVIAQLTG |
| 21. TFJ84559_Microchloropsis_salina_Eustigmatophyte | 283 | 1.12e-13 | FFDRFRESLA | HPLAHARVRLSSPRLHLYKA | SMSITAQLNW |
| 20. Naga_100503g2_Microchloropsis_gaditana_eustigmato | 237 | 1.12e-13 | FFDRFRESLA | HPLAHARVRLSSPRLHLYKA | SMSITAQLNW |
| 17. A0A2I1GAN6_Rhizophagus_irregularis | 193 | 1.94e-12 | MFENMIESAE | KPITKALITISNSNLDVYNA | QIRLDAHFRG |

### Motif G

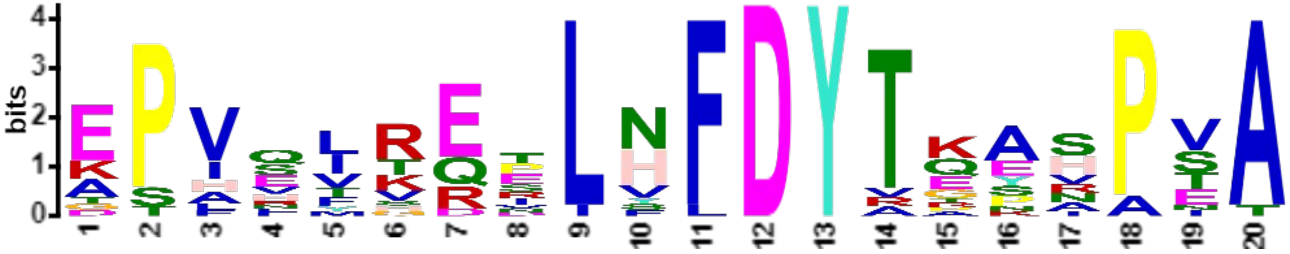

***E*-value:** 3.3e-092
**Site Count:** 18
**Width:** 20

**Log Likelihood Ratio:** 639
**Information Content:** 54.2
**Relative Entropy:** 51.2
**Bayes Threshold:** 10.3464

| | | | | | |
|---|---|---|---|---|---|
| 44. A0A6P6UED4_COFAR | 134 | 2.98e-20 | GVGLVRFWAE | EPVFIGERLHFDYTKAHPVA | AFSFHCDSGC |
| 35. PTQ31108_Marchantia_polymorpha_Liverworts | 565 | 8.37e-20 | DLILVSRIVE | EPVQLREVLNFDYTQARPSA | TFPVLSGQDL |
| 27. XP_020079770_Seipin-2-like_Ananas | 265 | 1.16e-19 | GSWLAGKVLE | KPIQLTEELNFDYTKASPVA | LVPVISCSNV |
| 45. A0A6P6W971_COFAR | 305 | 6.26e-19 | GGILMNAAVE | EPVRIKEILNFDYTQKSPIA | YVPIIGCPGP |
| 32. KAG0592613_Ceratodon_purpureus_Moss | 362 | 1.68e-18 | NIWFARGFIE | EPVEFREILHFDYRQEHPSA | TVSLLPPTIL |
| 34. XP_024544198_Selaginella_moellendorffii_Lycophyte | 285 | 4.25e-18 | DMVLMSRLVE | EPVQLRESINFDYTEARPSA | RVPLFPQVVT |
| 33. KAH8935238_Sphagnum_fallax_Moss | 381 | 8.03e-18 | DVFLVRSFIE | DPIEIHQTLHFDYTREHPTA | VVPLLPPKVL |
| 26. F4I340_SEI2_Arabidopsis_thaliana_Eudicot | 293 | 9.10e-18 | GGYVINRIAD | KPFEVKETLNFDYTKNSPEA | YVPISSCAGV |
| 30. XP_020106670_Seipin-1_Ananas_comosus_Monocot | 136 | 1.03e-17 | SLSLVKLLID | EPVHVRQTLYFDYTQPHPNT | VIALGGPKTR |
| 25. Q8L615_SEI3_Arabidopsis_thaliana_Eudicot | 269 | 5.41e-17 | SGFVITYLAH | EPLVIKESLNFDYTKSSPEA | YVPISSCAGV |
| 29. Q9FFD9_SEI1_Arabidopsis_thaliana_Eudicot | 123 | 3.47e-16 | GVGIVSLYVE | KPVVVRDRLFFDYTEENPSA | VFSFDKKKRS |
| 46. A0A6P6S6A8_COFAR | 273 | 4.28e-16 | GGITMRHLVV | ESIQTTENLNFDYTTSSPVA | FVPVTSSPVT |
| 31. XP_042467429_Seipin-1_Zingiber_officinale_Monocot | 115 | 4.28e-16 | GVALVRMWAE | EPVLLRRPLHLDYTEANPTA | SVALAGGTWH |
| 28. XP_042380030_Seipin-2_Zingiber_officinale_Monocot | 239 | 2.67e-14 | TFLAMRRVVE | APVSMTEELSFDYAKPIPEA | LVPVTTSDGC |
| 42. XP_042914764_Chlamydomonas_reinhardtii_Chlorophyta | 45 | 9.07e-14 | FLFLTRYYAP | ATHHFVRPLHFDYTGYVAVA | VAPLSPNLHE |
| 36. XP_001416623.1_Ostreococcus_lucimarinus_Mamiellale | 60 | 9.07e-14 | FALRYIVVPR | GPASLSQDLVFDYTAAAPVA | RVSFIPQKFA |
| 37. XP_002506104.1_Micromonas_commoda_Mamiellales | 72 | 1.78e-13 | VVFRILVVPT | TPASTRQELVFDYVPAAPTA | VSSFLGAKET |
| 43. GIL71655_Volvox_reticuliferus_Chlorophyceae | 69 | 2.11e-13 | FLYLTRRYAP | ASHNYVRPLHFDYTGYVAVA | LAHLSPNLRE |

### Motif H

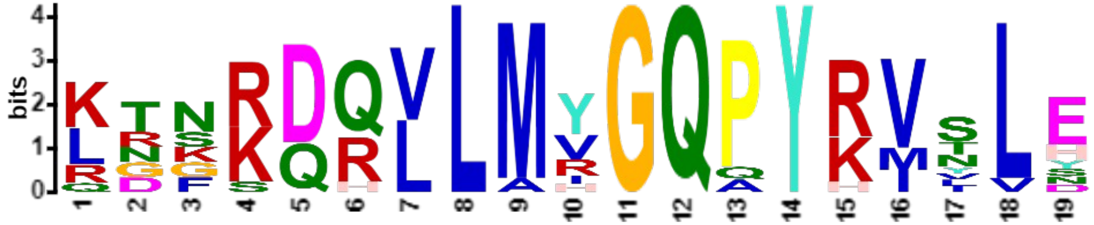

***E*-value:** 3.4e-073
**Site Count:** 12
**Width:** 19

**Log Likelihood Ratio:** 493
**Information Content:** 60.4
**Relative Entropy:** 59.3
**Bayes Threshold:** 11.2483

| | | | | | |
|---|---|---|---|---|---|
| 7. A0A8C5QNX3_Leptobrachium_leishanense | 93 | 4.45e-22 | SFPTANVSLL | KNNRDRVLMYGQPYRMSLE | LHVPESFVNQ |
| 3. A0A2D4I9W1_Micrurus_lemniscatus_lemniscatus | 92 | 4.50e-21 | SFPVANISFI | KDSRDQVLMYGQPYRMSLE | LELPESPVNQ |
| 12. XP_003702285_Megachile_rotundata | 93 | 8.01e-21 | ICSFPSAHVQ | LTNKQQLLMVGQPYKVNLH | LEMPESPANR |
| 11. A0A154P6K5_Dufourea_novaeangliae | 93 | 8.01e-21 | ICSFPSAYVQ | LTNKQQLLMVGQPYKVNLH | LEMPESPANK |
| 4. A0A8D2IST9_Varanus_komodoensis | 92 | 9.26e-21 | SFPTANISFV | KDSRDQVLMYGQPYRISLE | LELPESPVNQ |
| 6. XP_013976703_Canis_lupus_familiaris_Carnivors | 155 | 2.09e-20 | SFPVANVSLA | KGGRDRVLMYGQPYRVTLE | LELPESPVNQ |
| 5. NP_001116427_Homo_sapiens_Primates | 93 | 2.09e-20 | SFPVANVSLT | KGGRDRVLMYGQPYRVTLE | LELPESPVNQ |
| 8. A0A1L8GJQ5_Xenopus_laevis | 93 | 8.21e-20 | SFPMANVSLL | RNNRDRVLMHGQPYRISLE | LQLPESIVNQ |
| 14. XP_026461669_Ctenocephalides_felis_Fleas | 117 | 9.13e-19 | VCSFPSAHVS | LTKKQQLLMIGQPYKVYLY | IEMPESDANK |
| 13. Q9V3X4_Drosophila_melanogaster_Flies | 113 | 9.70e-17 | PCTFPHAHVS | LTKKQQLLMVGQAYKVIVN | IDMPESPQNL |
| 16. A0A7R9A2R1_Darwinula_stevensoni | 93 | 8.32e-16 | SFPSANVTLA | QRFKDHLLMRGQQYHVLLD | LDMPESPMNQ |
| 15. A0A7R9BHT2_Notodromas_monacha | 91 | 1.31e-14 | TYPSAYVELT | RRFSDQLLARGQPYRVVLS | LDVPESHVNQ |

#### Motif I

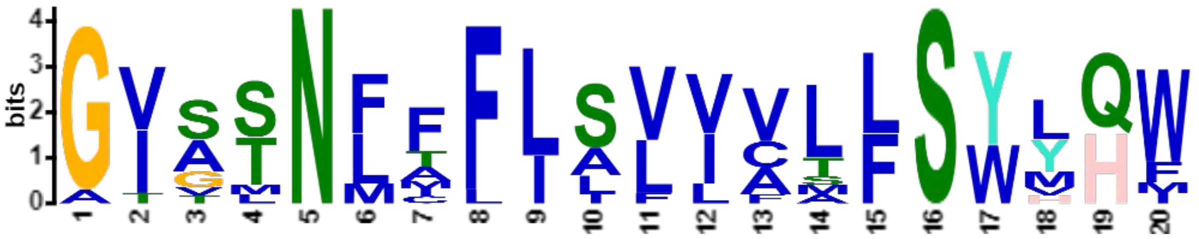

***E*-value:** 7.0e-078
**Site Count:** 13
**Width:** 20

**Log Likelihood Ratio:** 536
**Information Content:** 61.9
**Relative Entropy:** 59.5
**Bayes Threshold:** 10.4373

| | | | | | |
|---|---|---|---|---|---|
| 6. XP_013976703_Canis_lupus_familiaris_Carnivors | 300 | 7.70e-22 | YNFPMTCAFI | GVASNFTFLSVIVLFSYMQW | VWGGIWPRHR |
| 5. NP_001116427_Homo_sapiens_Primates | 238 | 7.70e-22 | YNFPMTCAFI | GVASNFTFLSVIVLFSYMQW | VWGGIWPRHR |
| 3. A0A2D4I9W1_Micrurus_lemniscatus_lemniscatus | 237 | 1.30e-21 | YNFPMTSAIL | GVASNFMFLSVIVLFSYLQW | IWGSLWPKET |
| 4. A0A8D2IST9_Varanus_komodoensis | 237 | 2.63e-20 | YNFPVTSAIL | GVVSNFAFLSVIVLFSYLQW | MWGSMWPREP |
| 7. A0A8C5QNX3_Leptobrachium_leishanense | 238 | 3.36e-20 | YNYPISTAII | GVSSNFFFLSVVVMLSYVQW | GLGRPRGQAD |
| 8. A0A1L8GJQ5_Xenopus_laevis | 238 | 4.80e-20 | YNFPITSAVI | GISSNFIFLSVLVLLSYLQW | GFGRTRLQDV |
| 16. A0A7R9A2R1_Darwinula_stevensoni | 238 | 2.09e-19 | FHFPVLSAVI | GIATNLFFLILVASFSWYHW | VFLPYHENVV |
| 14. XP_026461669_Ctenocephalides_felis_Fleas | 262 | 3.58e-19 | FHWPILSAAI | GISTNLFFIALVCLLSWYHM | SDTTWLEEVG |
| 13. Q9V3X4_Drosophila_melanogaster_Flies | 258 | 1.53e-18 | FNWPVLSAIV | AISTNLFFILVVFLLSWYHW | SDAKWLHSVQ |
| 12. XP_003702285_Megachile_rotundata | 238 | 6.69e-18 | YHWPILSAVV | GIGTNLFFIALVCTLSYLHF | TTYEDTDDDD |
| 11. A0A154P6K5_Dufourea_novaeangliae | 238 | 6.69e-18 | FHWPILSAIV | GIGTNLFFIALVCTLSYLHF | TTYEDTGDDS |
| 15. A0A7R9BHT2_Notodromas_monacha | 236 | 6.11e-17 | YHYPVLSAVC | GVSMNMAFLLLLAAFSWHHW | FLTPFFHQQS |
| 9. A0A8B8E1X8_Crassostrea_virginica | 238 | 3.17e-14 | FHWPLLSAVI | GTTLNMCLLSFIALLSWYQI | YYSQSNGNSN |

### Motif J

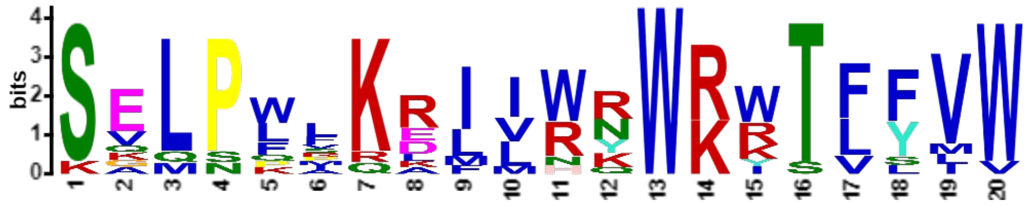

***E*-value:** 5.0e-074
**Site Count:** 12
**Width:** 20

**Log Likelihood Ratio:** 505
**Information Content:** 58.4
**Relative Entropy:** 60.7
**Bayes Threshold:** 11.8974

| | | | | | |
|---|---|---|---|---|---|
| 25. Q8L615_SEI3_Arabidopsis_thaliana_Eudicot | 439 | 2.30e-22 | EIYDASLFLE | SKLPFLKRIIWNWRKTLFVW | ISMSLFIMEL |
| 32. KAG0592613_Ceratodon_purpureus_Moss | 534 | 8.68e-22 | QIYSAEAQVL | SVLPWRKDILRRWKWTFYVW | SFLSVFMFEV |
| 44. A0A6P6UED4_COFAR | 299 | 1.44e-21 | QLYEAEILLK | SELPWAKELVHRWKWTFYVW | TSMHIYVFLL |
| 35. PTQ31108_Marchantia_polymorpha_Liverworts | 735 | 4.41e-21 | ELYEAELHLE | SALPWPKAIMRRWKWTFYVW | NGVCLFLFEV |
| 28. XP_042380030_Seipin-2_Zingiber_officinale_Monocot | 410 | 2.27e-20 | EIYSASMKLE | SELPLVKRILWNWKITVFVW | TTMVVFILLL |
| 27. XP_020079770_Seipin-2-like_Ananas | 436 | 4.47e-20 | EIYTASLKLE | SELPLFKRIIWNWRRTLLMW | LSMGIFIFEL |
| 46. A0A6P6S6A8_COFAR | 449 | 1.33e-19 | EIYAASLHIE | SELPQIKRMIWYWRRSVFVW | ISIITFLTEL |
| 26. F4I340_SEI2_Arabidopsis_thaliana_Eudicot | 467 | 1.48e-19 | ELYDASLSVE | SGLPFFRKIIWKWRKTLFVW | ISMSLFITEL |
| 45. A0A6P6W971_COFAR | 479 | 3.09e-19 | EIYAASLTLE | SEQPLLKRIVWYWRRTLFIW | VSMTVFTVEL |
| 33. KAH8935238_Sphagnum_fallax_Moss | 555 | 7.79e-19 | ELYSAELQVI | SVLSPMKDFIRKWRWTFYVW | SLLGLFMVEV |
| 31. XP_042467429_Seipin-1_Zingiber_officinale_Monocot | 271 | 1.18e-17 | QLYESELLMR | SQLPWSKELVNRWRWTFSLW | TSLNVFFALL |
| 47. EJK50087_Thalassiosira_oceanica_Diatom | 311 | 3.40e-12 | QIHSAQVKFG | KEMNKLQLLLRQWKYTFFVV | GTITIFVCYI |

### Motif K

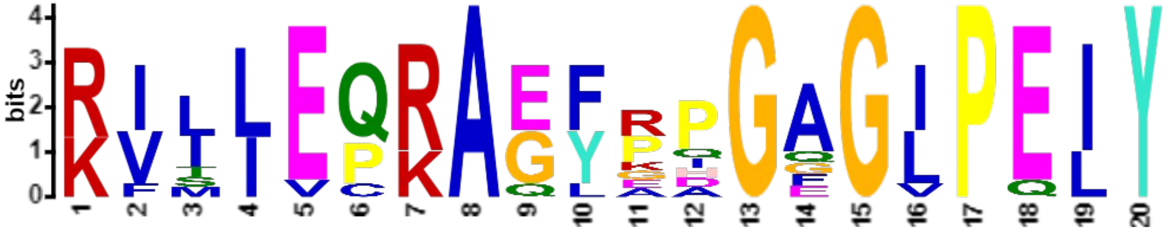


***E*-value:** 1.7e-056
**Site Count:** 10
**Width:** 20

**Log Likelihood Ratio:** 435
**Information Content:** 66.7
**Relative Entropy:** 62.8
**Bayes Threshold:** 11.3122

| | | | | | |
|---|---|---|---|---|---|
| 25. Q8L615_SEI3_Arabidopsis_thaliana_Eudicot | 412 | 5.13e-24 | EKDIIPTACL | KIMIEQRAEFRPGAGIPEIY | DASLFLESKL |
| 27. XP_020079770_Seipin-2-like_Ananas | 409 | 5.50e-23 | TEKPEPTACI | RVILEQRAEFKPGAGIPEIY | TASLKLESEL |
| 26. F4I340_SEI2_Arabidopsis_thaliana_Eudicot | 440 | 7.39e-23 | VEKDIPTACL | KIIIEQRAEFRPGAGIPELY | DASLSVESGL |
| 34. XP_024544198_Selaginella_moellendorffii_Lycophyte | 426 | 7.14e-21 | RSSSSYPGSA | KILLEPRAGFGPGAGLPEIY | NAEVLVQSQL |
| 28. XP_042380030_Seipin-2_Zingiber_officinale_Monocot | 383 | 1.47e-20 | TEGTKPTTCI | RVSLEQRAEYRDGAGIPEIY | SASMKLESEL |
| 46. A0A6P6S6A8_COFAR | 422 | 6.99e-20 | TEGYEPTACF | KVIIEQRAEYEAGFGIPEIY | AASLHIESEL |
| 35. PTQ31108_Marchantia_polymorpha_Liverworts | 708 | 3.31e-18 | QEKLVPTASV | RILLEPKAGYPQGGGLPELY | EAELHLESAL |
| 33. KAH8935238_Sphagnum_fallax_Moss | 528 | 6.34e-18 | EDTSVPTVFV | RILLECKAGLPPGEGLPELY | SAELQVISVL |
| 45. A0A6P6W971_COFAR | 452 | 3.83e-17 | TEGGRPTSCL | RVTIVQRAQFAHGAGVPEIY | AASLTLESEQ |
| 32. KAG0592613_Ceratodon_purpureus_Moss | 507 | 2.11e-16 | EETKIPTAMV | RFLLEPKAGYPIGQGLPQIY | SAEAQVLSVL |

### Motif L

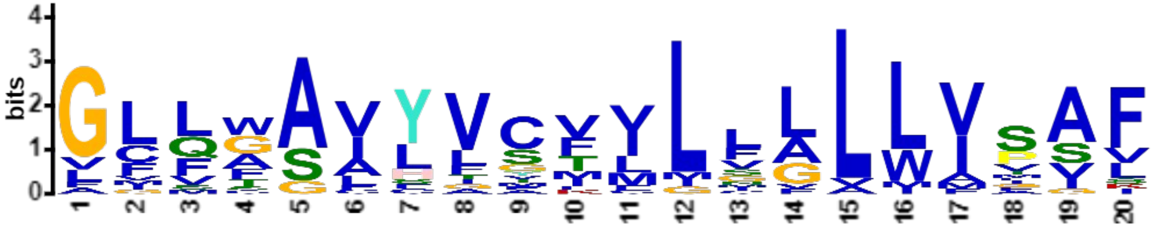


***E*-value:** 2.8e-049
**Site Count:** 21
**Width:** 20

**Log Likelihood Ratio:** 604
**Information Content:** 47.5
**Relative Entropy:** 41.5
**Bayes Threshold:** 9.48391

| | | | | | |
|---|---|---|---|---|---|
| 45. A0A6P6W971_COFAR | 273 | 1.04e-18 | LLELGLKFVW | GLLWSVYVCVVLVVLLVSAF | VVGGILMNAA |
| 25. Q8L615_SEI3_Arabidopsis_thaliana_Eudicot | 237 | 1.58e-17 | VLELVRRVTW | GLFCAVYVGIMLFALLVSAF | MISGFVITYL |
| 33. KAH8935238_Sphagnum_fallax_Moss | 349 | 3.39e-17 | IKHITKRCGC | GCLAAIYVSFILGFLLIPAF | FLDVFLVRSF |
| 35. PTQ31108_Marchantia_polymorpha_Liverworts | 533 | 4.23e-16 | VTKLSKKTMM | GCLGSAYVVFVLSLLLIPAF | FVDLILVSRI |
| 32. KAG0592613_Ceratodon_purpureus_Moss | 330 | 8.36e-16 | LRQTAKRCGF | GCVAAVYVMFMLGALLVPAL | FLNIWFARGF |
| 44. A0A6P6UED4_COFAR | 102 | 1.05e-15 | CGVLLNKVFL | GFLGAVHVCKILMLLLVVAV | ILGVGLVRFW |
| 26. F4I340_SEI2_Arabidopsis_thaliana_Eudicot | 261 | 5.31e-15 | MLSIVCKFGW | GMFWAVYVGIVLFGLLVSSL | MIGGYVINRI |
| 27. XP_020079770_Seipin-2-like_Ananas | 233 | 7.27e-15 | LWGLAVRLII | GGVWAFYICFVLCGLLATAF | LGGSWLAGKV |
| 28. XP_042380030_Seipin-2_Zingiber_officinale_Monocot | 207 | 2.03e-14 | AVKLLVRLAW | GCFWSLYVCFILFALLAASS | LATFLAMRRV |
| 8. A0A1L8GJQ5_Xenopus_laevis | 28 | 6.01e-14 | LLMFLRARRI | FLQAAILLCVLLLLLWVSVF | LYGSFYYSYM |
| 4. A0A8D2IST9_Varanus_komodoensis | 27 | 7.29e-14 | ALLMLRVRRT | VLQTAILLCVLLLLLWISIF | LYGSFYYSYM |
| 3. A0A2D4I9W1_Micrurus_lemniscatus_lemniscatus | 27 | 7.29e-14 | SLLMLRVRRT | VLQTAILLCVLLLLLWISIF | LYGSFYYSYM |
| 6. XP_013976703_Canis_lupus_familiaris_Carnivors | 90 | 9.73e-14 | HILAGRARKL | LLQFGVLFCTILLLLWVSVF | LYGSFYYSYM |
| 5. NP_001116427_Homo_sapiens_Primates | 28 | 9.73e-14 | QVLAGRARRL | LLQFGVLFCTILLLLWVSVF | LYGSFYYSYM |
| 46. A0A6P6S6A8_COFAR | 241 | 7.41e-13 | MLKVAVRFCW | AFFWSVYVYLILVGLLIMGF | VIGGITMRHL |
| 34. XP_024544198_Selaginella_moellendorffii_Lycophyte | 253 | 7.41e-13 | VSRMVKRMGY | GVLSAAYVFMLLNLLLVPAI | LMDMVLMSRL |
| 29. Q9FFD9_SEI1_Arabidopsis_thaliana_Eudicot | 91 | 1.56e-11 | AGRVVRRTWF | GILGACHVSMVMVLALILAV | VIGVGIVSLY |
| 30. XP_020106670_Seipin-1_Ananas_comosus_Monocot | 104 | 2.95e-11 | SAATLHQISF | GLLGAACTSVMLIGVMVVAV | LLSLSLVKLL |
| 43. GIL71655_Volvox_reticuliferus_Chlorophyceae | 271 | 9.36e-11 | LVASVALLLI | GLMASISGTVCGFALLIFAF | FLRRWIAKSV |
| 31. XP_042467429_Seipin-1_Zingiber_officinale_Monocot | 83 | 1.09e-10 | AAAIARRFAL | GLLGALYAATFLLALLFISL | LLGVALVRMW |
| 36. XP_001416623.1_Ostreococcus_lucimarinus_Mamiellale | 252 | 3.02e-9 | SFVLVGLFWS | GFSFAAFVSTVISLLIMGSK | IDEKDAESPM |

### Motif M

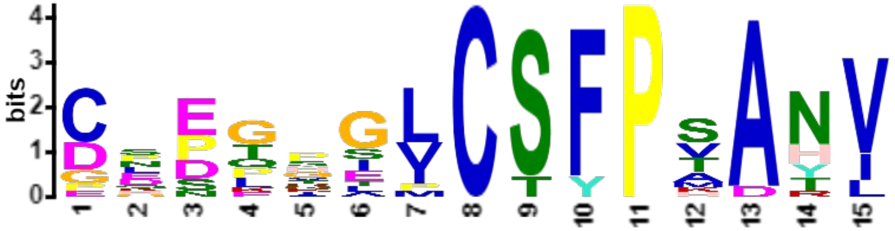

***E*-value:** 3.2e-052
**Site Count:** 16
**Width:** 15

**Log Likelihood Ratio:** 445
**Information Content:** 40.3
**Relative Entropy:** 40.2
**Bayes Threshold:** 9.54936

| | | | | | |
|---|---|---|---|---|---|
| 12. XP_003702285_Megachile_rotundata | 77 | 4.84e-16 | IRPVHLQFKS | CNEQRGICSFPSAHV | QLTNKQQLLM |
| 11. A0A154P6K5_Dufourea_novaeangliae | 77 | 5.76e-16 | IRPVHLQFKS | CNEQKGICSFPSAYV | QLTNKQQLLM |
| 16. A0A7R9A2R1_Darwinula_stevensoni | 75 | 6.83e-16 | TRPVHFIFEP | CKEEAGLCSFPSANV | TLAQRFKDHL |
| 14. XP_026461669_Ctenocephalides_felis_Fleas | 101 | 1.04e-14 | IRPVHLQFKS | CEEGQAVCSFPSAHV | SLTKKQQLLM |
| 7. A0A8C5QNX3_Leptobrachium_leishanense | 75 | 4.32e-14 | PVHYQYSSFC | DPPPGILCSFPTANV | SLLKNNRDRV |
| 10. A0A6P7TH74_Octopus_vulgaris | 73 | 1.77e-13 | ERPCNFVFDV | CDNGVGVCSFPKARV | DLTDESNRYV |
| 3. A0A2D4I9W1_Micrurus_lemniscatus_lemniscatus | 74 | 1.98e-13 | PVHYQFRTDC | GQPGPELCSFPVANI | SFIKDSRDQV |
| 15. A0A7R9BHT2_Notodromas_monacha | 73 | 3.43e-13 | MRPVDFKFKP | CGEKPGLCTYPSAYV | ELTRRFSDQL |
| 4. A0A8D2IST9_Varanus_komodoensis | 74 | 3.43e-13 | PVHYHFRTDC | GLPGPELCSFPTANI | SFVKDSRDQV |
| 8. A0A1L8GJQ5_Xenopus_laevis | 75 | 6.47e-13 | PVHYQYSSTC | EPPPGILCSFPMANV | SLLRNNRDRV |
| 5. NP_001116427_Homo_sapiens_Primates | 75 | 7.17e-13 | PVHFYYRTDC | DSSTTSLCSFPVANV | SLTKGGRDRV |
| 13. Q9V3X4_Drosophila_melanogaster_Flies | 97 | 1.46e-12 | TRPVHMQFKT | CLETSTPCTFPHAHV | SLTKKQQLLM |
| 6. XP_013976703_Canis_lupus_familiaris_Carnivors | 137 | 1.46e-12 | PVHFYYRTDC | DSSTSLLCSFPVANV | SLAKGGRDRV |
| 9. A0A8B8E1X8_Crassostrea_virginica | 73 | 2.61e-12 | TRPVNLQFKI | CEDGIGMCSYPADNI | TLVQEGQTEV |
| 39. A0A2A2K4L1_Diploscapter_pachys | 68 | 1.24e-11 | IPLNLVFSTC | PDDLHGVCSFPTATL | EYDQNSLFTR |
| 38. Q8MXG1_Caenorhabditis_elegans | 52 | 2.70e-11 | YQLNIVFQTC | DHDLHSVCSFPAATL | EYEKNSLFSP |

### Motif N

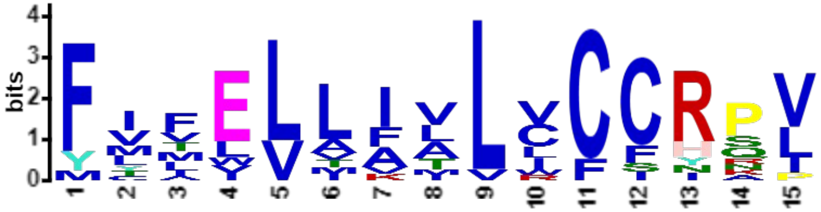


***E*-value:** 6.3e-030
**Site Count:** 14
**Width:** 15

**Log Likelihood Ratio:** 379
**Information Content:** 39.4
**Relative Entropy:** 39
**Bayes Threshold:** 11.197

| | | | | | |
|---|---|---|---|---|---|
| 27. XP_020079770_Seipin-2-like_Ananas | 461 | 1.06e-17 | TLLMWLSMGI | FIFELLILLVCCRPV | IIPRTRSPIA |
| 33. KAH8935238_Sphagnum_fallax_Moss | 580 | 6.67e-16 | TFYVWSLLGL | FMVEVLIVLCCCHQV | LLPRRLLQGL |
| 26. F4I340_SEI2_Arabidopsis_thaliana_Eudicot | 492 | 8.89e-16 | TLFVWISMSL | FITELLFTLVCCRPL | IIPRTQPRDR |
| 35. PTQ31108_Marchantia_polymorpha_Liverworts | 760 | 2.34e-15 | TFYVWNGVCL | FLFEVAIILCCCRQV | LLPGLGGRGQ |
| 32. KAG0592613_Ceratodon_purpureus_Moss | 559 | 4.26e-14 | TFYVWSFLSV | FMFEVMVVLCCCHRV | LLPSSLLQGI |
| 28. XP_042380030_Seipin-2_Zingiber_officinale_Monocot | 435 | 1.12e-13 | TVFVWTTMVV | FILLLLAVLVCCRSI | LTPGARRGGG |
| 34. XP_024544198_Selaginella_moellendorffii_Lycophyte | 478 | 4.20e-13 | TCYVWGAVLI | FVLEVAAMVCCCRNV | LLVSPSGASP |
| 46. A0A6P6S6A8_COFAR | 474 | 1.74e-12 | SVFVWISIIT | FLTELIIALIFCRPV | IFPGRSTIKV |
| 45. A0A6P6W971_COFAR | 504 | 4.69e-12 | TLFIWVSMTV | FTVELLFTLLCCNSI | IIPRVNLGRT |
| 25. Q8L615_SEI3_Arabidopsis_thaliana_Eudicot | 464 | 1.32e-11 | TLFVWISMSL | FIMELLFALVFFRPL | IIPRTGQRTQ |
| 30. XP_020106670_Seipin-1_Ananas_comosus_Monocot | 312 | 1.02e-10 | WTFYVWTSFY | MYIVLLIVLICCYKP | SIFPTVRSTN |
| 29. Q9FFD9_SEI1_Arabidopsis_thaliana_Eudicot | 301 | 2.22e-10 | TLCVWTSMYL | YVAILTALLWCFRPV | LFPYTSSRTI |
| 2. XP_002286702_Thalassiosira_pseudonana_Diatom | 51 | 5.48e-10 | HDIFFQGILP | FCMWLLKALLCIIAL | LILSMGTYWL |
| 44. A0A6P6UED4_COFAR | 324 | 5.91e-10 | TFYVWTSMHI | YVFLLVILLRCSRPL | ILPVMRKSPS |

### Motif O

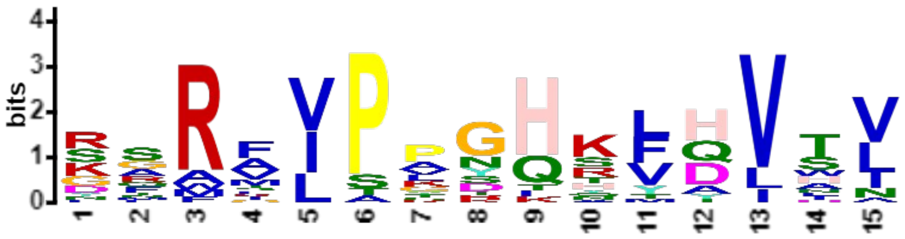


***E*-value:** 2.4e-018
**Site Count:** 21
**Width:** 15

**Log Likelihood Ratio:** 442
**Information Content:** 32
**Relative Entropy:** 30.4
**Bayes Threshold:** 10.3107

| | | | | | |
|---|---|---|---|---|---|
| 33. KAH8935238_Sphagnum_fallax_Moss | 426 | 4.79e-14 | LAVKKPEKAY | SFRAIPKSHKFHVTV | FLTLPESDHN |
| 32. KAG0592613_Ceratodon_purpureus_Moss | 405 | 5.35e-13 | KGGIYPNKVL | YARAIPPSHKLQVTV | FLTLPESHYN |
| 45. A0A6P6W971_COFAR | 351 | 1.38e-12 | SENFDVLKFD | GMRVIPLDHKLQVTV | SLTLPESDYN |
| 35. PTQ31108_Marchantia_polymorpha_Liverworts | 607 | 2.61e-12 | AKNLPLEKLV | SFRKIPAGHYFHVNV | ILTMPESEYN |
| 27. XP_020079770_Seipin-2-like_Ananas | 308 | 4.28e-12 | EGKGRAGAFL | DRRMVPANHKLQLTI | SLTLPESDYN |
| 46. A0A6P6S6A8_COFAR | 321 | 9.81e-12 | EDRMLFSKSA | GGRTIPYNHKLQLTV | LLTMPESEYN |
| 41. KAG7673487_Chlorella_desiccata_Trebouxiophyceae | 368 | 3.00e-11 | AIAADIKIDP | NSRFLPPGQRMDVWL | DFTVPALGDG |
| 31. XP_042467429_Seipin-1_Zingiber_officinale_Monocot | 145 | 6.97e-11 | SVALAGGTWH | SGRAVPPGHSVTVSL | LILLPESDYN |
| 21. TFJ84559_Microchloropsis_salina_Eustigmatophyte | 389 | 1.28e-10 | RDGLGGREGG | RQAFVPGYHHYHVAN | DEAEVLLGAG |
| 20. Naga_100503g2_Microchloropsis_gaditana_eustigmato | 342 | 1.28e-10 | RDGLGGREGA | RQAFVPGYHHYHVAN | DEAEVLLGAG |
| 40. A0A2P6VDK0_Micractinium_conductrix | 142 | 1.03e-9 | DGLPPAPLPS | DARFLAPGQAVDVWV | ELVVPRSTQQ |
| 34. XP_024544198_Selaginella_moellendorffii_Lycophyte | 322 | 1.03e-9 | VVTDKSVSSL | KAWAIPKTHSFHITA | TLQLPESDHN |
| 30. XP_020106670_Seipin-1_Ananas_comosus_Monocot | 163 | 1.12e-9 | PNTVIALGGP | KTRMVPTRHTTHVLL | NMLMPESNHN |
| 37. XP_002506104.1_Micromonas_commoda_Mamiellales | 120 | 1.34e-9 | PKLITDAALT | SSRVLTPGQRFDVSV | SLTLPETRHN |
| 42. XP_042914764_Chlamydomonas_reinhardtii_Chlorophyta | 96 | 1.47e-9 | PGAAAAAVLP | KSRFLPLGTQVAVHV | KLVIPWDHTD |
| 29. Q9FFD9_SEI1_Arabidopsis_thaliana_Eudicot | 151 | 2.70e-9 | SAVFSFDKKK | RSFSVPVGHSVHVSL | VLWMPESEIN |
| 43. GIL71655_Volvox_reticuliferus_Chlorophyceae | 118 | 4.14e-9 | WPVSAANLLP | RSRFLPSGLRVAVHV | VLTIPADHTD |
| 44. A0A6P6UED4_COFAR | 174 | 6.82e-9 | QGYTHQINYK | RNMGVPVGHTFYVSL | VFLMPESDYN |
| 36. XP_001416623.1_Ostreococcus_lucimarinus_Mamiellale | 98 | 1.02e-8 | FANTGESKPR | KGRIVSAKQNFDVDI | EFVVPDSEYN |
| 26. F4I340_SEI2_Arabidopsis_thaliana_Eudicot | 339 | 1.41e-8 | KESNEMSKIR | GLRVIPRDQKLDIIL | SMTLPESAYN |
| 28. XP_042380030_Seipin-2_Zingiber_officinale_Monocot | 282 | 3.59e-8 | GETIDVVGHE | FRRLVSPNKKLQLTI | SLKLPESDYN |

### Motif P

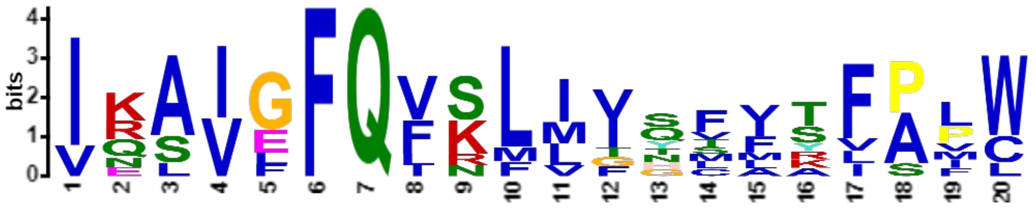

***E*-value:** 5.4e-016
**Site Count:** 9
**Width:** 20

**Log Likelihood Ratio:** 330
**Information Content:** 54.4
**Relative Entropy:** 52.9
**Bayes Threshold:** 11.347

| | | | | | |
|---|---|---|---|---|---|
| 45. A0A6P6W971_COFAR | 192 | 2.77e-19 | SFLMVLVRLV | IKAIGFQFSLLFSFFTFPVW | AIYTSYMFVM |
| 33. KAH8935238_Sphagnum_fallax_Moss | 268 | 9.87e-19 | QLLVWPADFI | VQAIGFQIKLIVQSVYFALW | LCSLTYAVLV |
| 32. KAG0592613_Ceratodon_purpureus_Moss | 249 | 3.66e-18 | QILASPVGIA | IELVGFQVKLIVQMFSFALW | FMSFGVSMMS |
| 25. Q8L615_SEI3_Arabidopsis_thaliana_Eudicot | 167 | 1.03e-17 | STDWSLTSLV | IRSIEFQVSLMITFIRFPPW | LISKCLSFVF |
| 26. F4I340_SEI2_Arabidopsis_thaliana_Eudicot | 183 | 1.55e-16 | SLLGFLVGLV | IKAIEFQVSFMTSLLTFPPW | LLRNCFLFFF |
| 35. PTQ31108_Marchantia_polymorpha_Liverworts | 452 | 1.87e-16 | HFLSWPAELL | VQAVGFQVRLIIHVVAVALW | MFNFTCSLIS |
| 27. XP_020079770_Seipin-2-like_Ananas | 152 | 1.01e-15 | TLLECVAGFV | IRAVLFQLNLVVNCITFPIC | LLHCSFLLVI |
| 34. XP_024544198_Selaginella_moellendorffii_Lycophyte | 172 | 1.45e-14 | HLLLWPVGFL | INAIGFQFKMIGYTMSIAML | LFSICFSMVT |
| 28. XP_042380030_Seipin-2_Zingiber_officinale_Monocot | 126 | 9.79e-13 | GFLESSVDVV | IKSVFFQFSLLIGFAKLSFC | VLFCPFYFFL |

### Motif Q

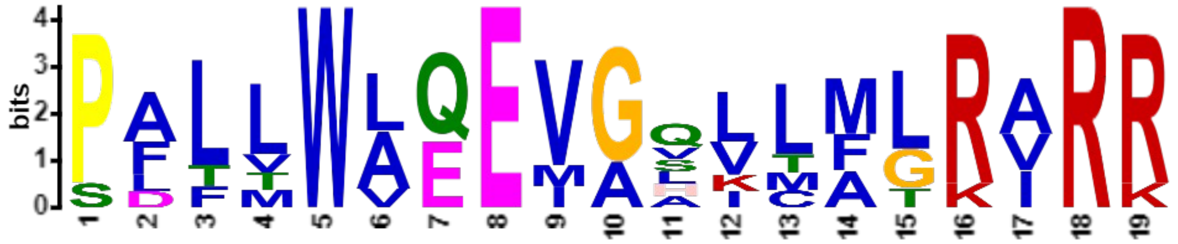


***E*-value:** 2.6e-014
**Site Count:** 7
**Width:** 19

**Log Likelihood Ratio:** 274
**Information Content:** 60.7
**Relative Entropy:** 56.4
**Bayes Threshold:** 11.4224

| | | | | | |
|---|---|---|---|---|---|
| 5. NP_001116427_Homo_sapiens_Primates | 8 | 9.44e-21 | MVNDPPV | PALLWAQEVGQVLAGRARR | LLLQFGVLFC |
| 6. XP_013976703_Canis_lupus_familiaris_Carnivors | 70 | 1.25e-18 | LPAMVNDPPV | PALLWAQEVGHILAGRARK | LLLQFGVLFC |
| 4. A0A8D2IST9_Varanus_komodoensis | 7 | 2.59e-18 | MSGNAG | PFLVWAQEVAALLMLRVRR | TVLQTAILLC |
| 3. A0A2D4I9W1_Micrurus_lemniscatus_lemniscatus | 7 | 2.91e-18 | MSASPG | PFLMWVQEVASLLMLRVRR | TVLQTAILLC |
| 7. A0A8C5QNX3_Leptobrachium_leishanense | 8 | 1.61e-17 | MSHNVSP | PALLWLEEIGVVTFLRIRR | LILQAGLLVC |
| 8. A0A1L8GJQ5_Xenopus_laevis | 8 | 3.52e-16 | MSPTVSP | PLFLWLEEMGLLMFLRARR | IFLQAAILLC |
| 14. XP_026461669_Ctenocephalides_felis_Fleas | 282 | 1.24e-14 | LVCLLSWYHM | SDTTWLEEVGQKCMTKIRR | SKSKTVETED |

### Motif R

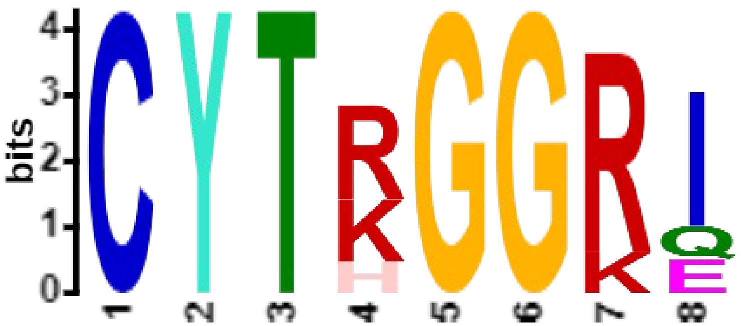


***E*-value:** 1.7e-011
**Site Count:** 6
**Width:** 8

**Log Likelihood Ratio:** 138
**Information Content:** 31.2
**Relative Entropy:** 33.2
**Bayes Threshold:** 12.2837

| | | | | | |
|---|---|---|---|---|---|
| 6. XP_013976703_Canis_lupus_familiaris_Carnivors | 194 | 1.45e-11 | DLGMFLVTIS | CYTRGGRI | ISTSSRSVML |
| 5. NP_001116427_Homo_sapiens_Primates | 132 | 1.45e-11 | DLGMFLVTIS | CYTRGGRI | ISTSSRSVML |
| 4. A0A8D2IST9_Varanus_komodoensis | 131 | 2.35e-11 | NLGMFMVVIS | CYTKGGRI | ISSSARSAML |
| 3. A0A2D4I9W1_Micrurus_lemniscatus_lemniscatus | 131 | 2.35e-11 | NLGMFMVVIS | CYTKGGRI | ISSSARAAML |
| 8. A0A1L8GJQ5_Xenopus_laevis | 132 | 3.89e-11 | DLGMFMVTMS | CYTRGGRQ | ISYTARSAML |
| 7. A0A8C5QNX3_Leptobrachium_leishanense | 132 | 1.68e-9 | DLGMFMVSMS | CYTHGGKE | ISHTARSAIL |

#### Motif S

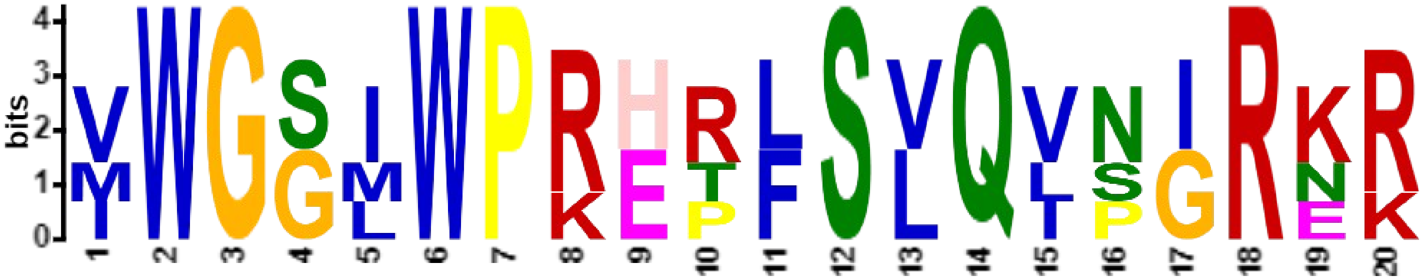

***E*-value:** 2.3e-009
**Site Count:** 4
**Width:** 20

**Log Likelihood Ratio:** 197
**Information Content:** 70.8
**Relative Entropy:** 71.1
**Bayes Threshold:** 12.3829

| | | | | | | |
|---|---|---|---|---|---|---|
| 6. | XP_013976703_Canis_lupus_familiaris_Carnivors | 320 | 3.40e-26 | VIVLFSYMQW | VWGGIWPRHRFSLQVNIRKR | DSSQKPVQRR |
| 5. | NP_001116427_Homo_sapiens_Primates | 258 | 3.40e-26 | VIVLFSYMQW | VWGGIWPRHRFSLQVNIRKR | DNSRKEVQRR |
| 4. | A0A8D2IST9_Varanus_komodoensis | 257 | 4.14e-19 | VIVLFSYLQW | MWGSMWPREPLSVQLSGRER | LGAAPQAEDG |
| 3. | A0A2D4I9W1_Micrurus_lemniscatus_lemniscatus | 257 | 1.83e-18 | VIVLFSYLQW | IWGSLWPKETLSVQIPGRNK | SGPVQTLDDS |

#### Motif T

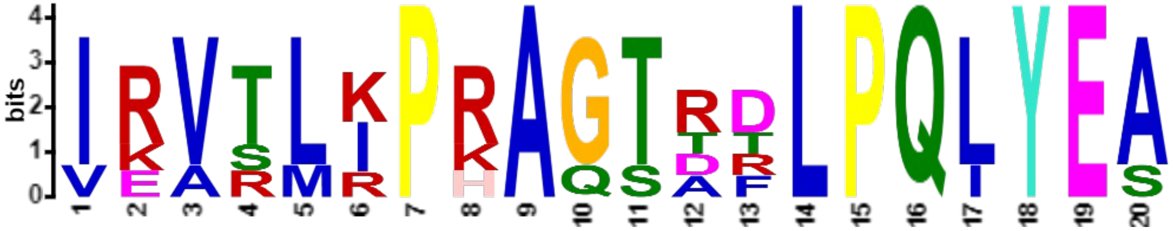

***E*-value:** 6.1e-012
**Site Count:** 5
**Width:** 20

**Log Likelihood Ratio:** 232
**Information Content:** 71.9
**Relative Entropy:** 66.8
**Bayes Threshold:** 12.3125

| | | | | | |
|---|---|---|---|---|---|
| 4. A0A6P6UED4_COFAR | 274 | 2.57e-22 | HKEDFPRTEA | IKVTLIPRAGTDFLPQLYEA | EILLKSELPW |
| 30. XP_020106670_Seipin-1_Ananas_comosus_Monocot | 263 | 3.28e-22 | YKERQPKTEL | IRVSLRPRAGTTDLPQLYEA | DIVIRTQLPW |
| 31. XP_042467429_Seipin-1_Zingiber_officinale_Monocot | 246 | 8.73e-21 | KESRAKRSGS | IRVRLKPKAGTRDLPQLYES | ELLMRSQLPW |
| 29. Q9FFD9_SEI1_Arabidopsis_thaliana_Eudicot | 251 | 2.18e-19 | HQEKMPRTKA | VRATLIPRAQTRTLPQLYEA | EIVINSKPPW |
| 36. XP_001416623.1_Ostreococcus_lucimarinus_Mamiellale | 200 | 9.28e-19 | KEDSESPFTD | IEVTMKPHAGSARLPQIYEA | RAIVHLSMNF |

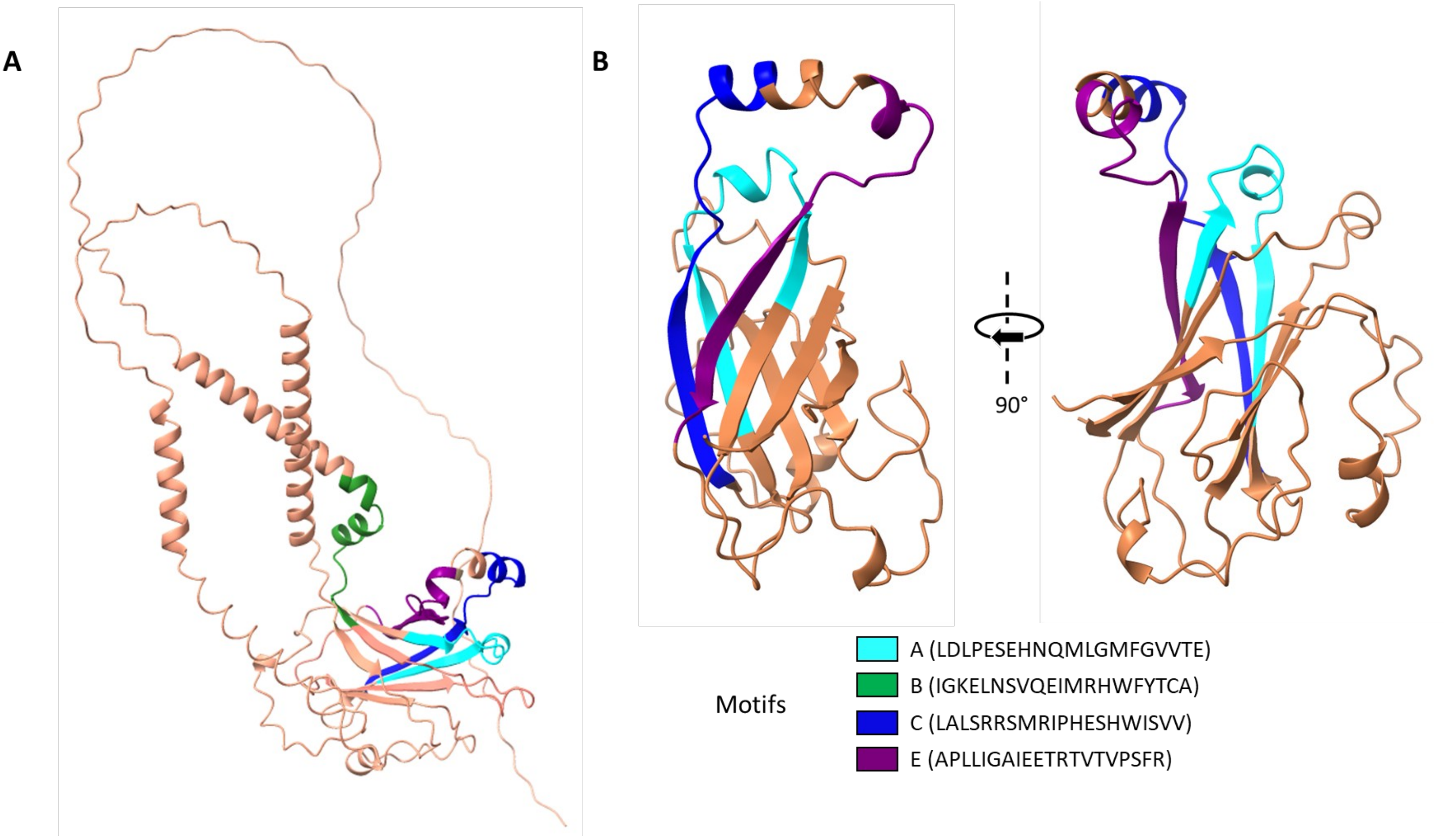


**Supplementary Figure S4: Positioning of the A, B, C and E motifs on the PtSeipin luminal domain**

Structure prediction was retrieved from the AlphaFold Database and visualization was performed using ChimeraX. Motif A is shown in light blue, motif B in green, motif C in dark blue and motif E in purple.

A. Complete structure overview
B. Zoom on the luminal domain

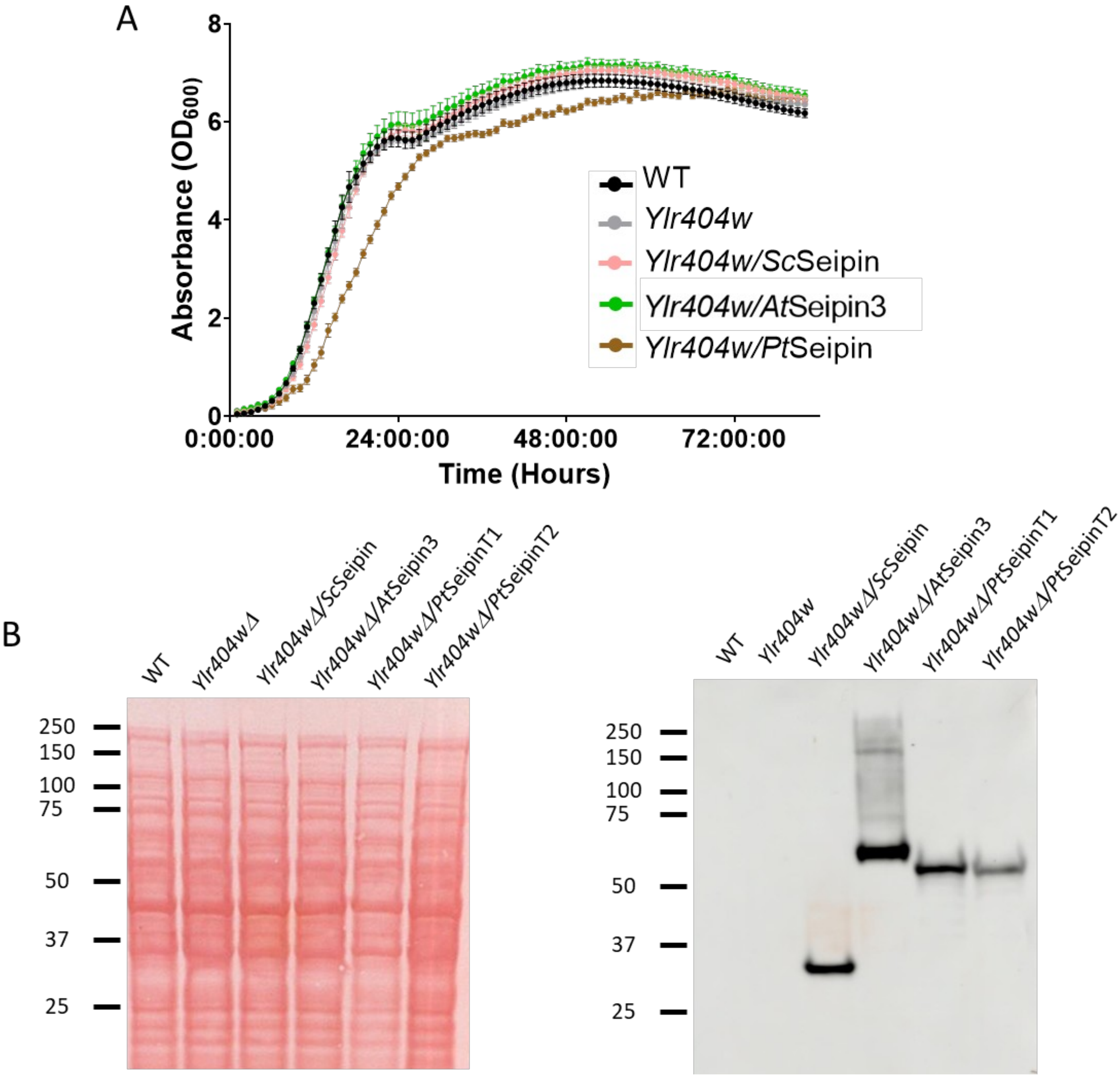


**Supplementary Figure S5: Heterologous expression in yeast**

A. Growth curves of yeast strains with or without Seipin complementation. Absorbance at 600 nm ($OD_{600}$) was measured every hour for 80 hours. Each point corresponds to the mean of 12 replicates grown in parallel ± standard deviation (SD).
B. Western Blot analysis of yeast strains with or without Seipin complementation. Ponceau staining of the membrane is shown on the left and immunoblotting with anti-HA antibody is shown on the right.
   Wild-type (WT) and Seipin deficient (*ylr404wΔ*) strains were transformed with the empty plasmid YCplac33. *ylr404wΔ* was complemented by yeast Seipin (*Ylr404wΔ/Sc*Seipin), *Arabidopsis* Seipin 3 (*Ylr404wΔ/At*Seipin*3*) or by *Phaeodactylum tricornutum* Seipin (*Ylr404wΔ/Pt*Seipin). As the latter led to growth delay, two different time points are shown. *Ylr404wΔ/Pt*SeipinT1 cells were harvested at the same time but at a lower $OD_{600}$ while *Ylr404wΔ/Pt*SeipinT2 were harvested at a later time but at similar $OD_{600}$.

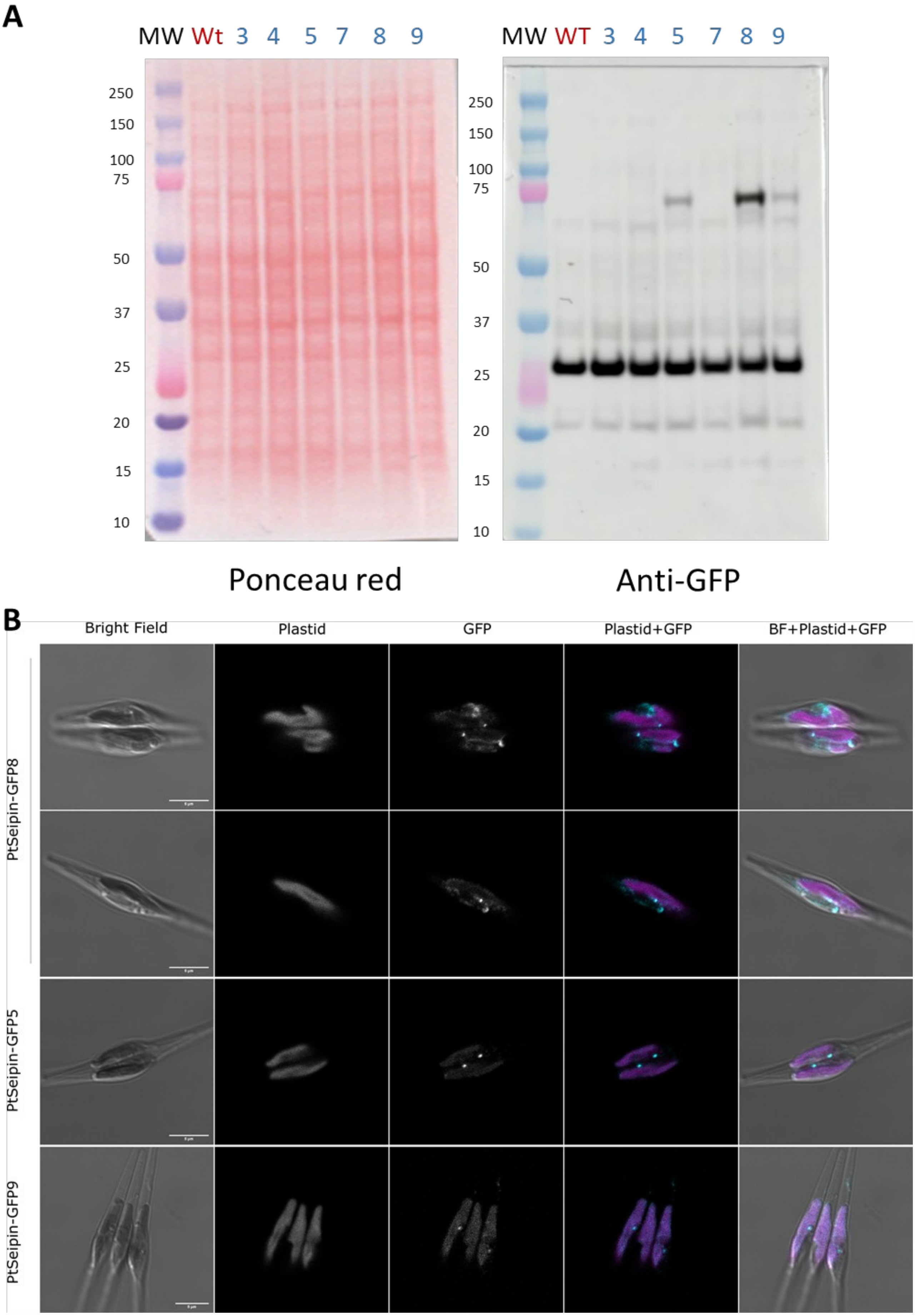
A
MW Wt 3 4 5 7 8 9
MW WT 3 4 5 7 8 9
250
150
100
75
50
37
25
20
15
10
Ponceau red
Anti-GFP
B
Bright Field
Plastid
GFP
Plastid+GFP
BF+Plastid+GFP
PtSeipin-GFP8
PtSeipin-GFP5
PtSeipin-GFP9

**Supplementary Figure S6: PtSeipin-eGFP overexpressing lines**

A. Western Blot analysis of PtSeipin-eGFP overexpressing lines. Ponceau staining of the membrane is shown on the left and immunoblotting with anti-GFP antibody is shown on the right. PtSeipin-eGFP expression is expected at about 75 kDa. PtSeipin-eGFP expression is only detected in line 5, line 8 and line 9.
B. Representative confocal images of PtSeipin-eGFP overexpressing lines 5, 8 and 9. From left to right: pseudo-brightfield image (grayscale); Plastid autofluorescence (grayscale), GFP (grayscale), overlay of plastid autofluorescence (magenta) and GFP (cyan), overlay of pseudo-brightfield (grayscale), plastid autofluorescence (magenta) and GFP (cyan). Scale bar: 5 µm. In the highest expressing line (PtSeipin-GFP8), signal is observed around the plastid, in the ER and at foci in the vicinity of LD. In the lowest expressing lines (PtSeipin-GFP5 and 9), the bright foci are still detected but little to no signal is observed around the plastid or in the ER. The increased plastid fluorescence detected in the GFP channel is due to changes in imaging parameters (laser power, photomultiplier gain) necessary to detect the signal.

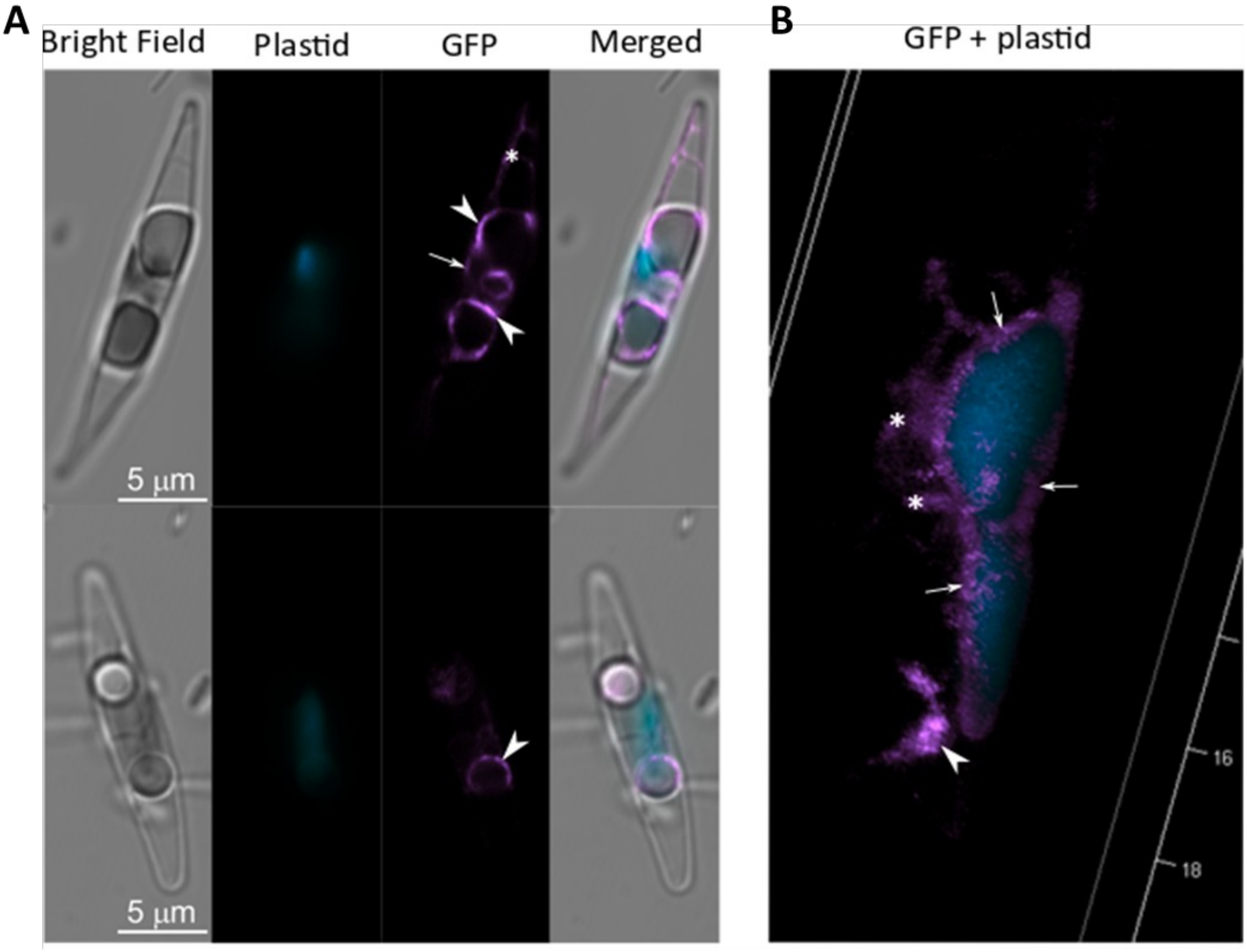


**Supplementary Figure S7: Laser Scanning Confocal Microscopy (LSCM) observation of the *in vivo* localization of PtSeipin *in Phaeodactylum tricornutum* in control culture conditions in stationary phase.**

A. Representative images of PtSeipin-eGFP overexpressing cells showing big lipid droplets. From left to right: BrightField signal (grey), Chlorophyll autofluorescence (cyan), GFP signal (magenta) and overlay of bright field with GFP and chlorophyll signals (respectively grey, magenta and cyan). Scale bar: 5 µm.
B. 3D reconstruction showing GFP signal (magenta) around the plastid (cyan).

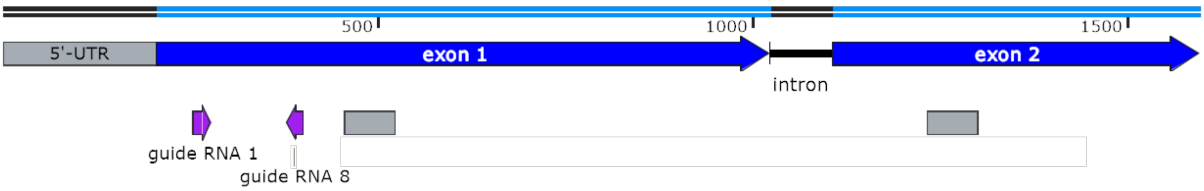


| Name | Mutation | Translated sequence |
|---|---|---|
| SeipinKO 1.1 | 884bp deletion including the first 78 bp of *PtSeipin* and the first 160 bp of *Phatr3_J7649* gene | N/A |
| SeipinKO 1.2 | 224 bp deletion including the region coding the first transmembrane domain of PtSeipin | MDRRVSRRRSQTSLVVAEEAHAVLDVLSSRHAVSALQ* |
| SeipinKO 1.3 | 2 bp insertion | MDRRVSRRRSQTSLVVAEEAHAVPRTHPRCNASLNTQRCVC* |
| SeipinKO 8.1 | 1 bp insertion | MDRRVSRRRSQTSLVVAEEAHAAEDPSALQRIVEYAAVRVLARQFPSR NAHGRSSTEIPFKSIPRAYS* |
| SeipinKO 8.2 | 5 bp deletion | MDRRVSRRRSQTSLVVAEEAHAAEDPSALQRIVEYAAVRVLARQFPSR NAHGRSSTEIPFIPRAYS* |
| SeipinKO 8.3 | Mix of 14 bp deletion<br><br>and 1 bp insertion | MDRRVSRRRSQTSLVVAEEAHAAEDPSALQRIVEYAAVRVLARQFPSR NAHGRSSTEIPFAYS*<br>MDRRVSRRRSQTSLVVAEEAHAAEDPSALQRIVEYAAVRVLARQFPSR NAHGRSSTEIPFKSIPRAYS* |

**Supplementary Figure S8: Generation of PtSeipin KO mutants**

Upper: Position of RNA guides 1 and 8 (purple arrows) are indicated on the *PtSeipin* gene layout. Positions of regions encoding the transmembrane domains are indicated with grey boxes.
Lower: description of obtained mutant lines

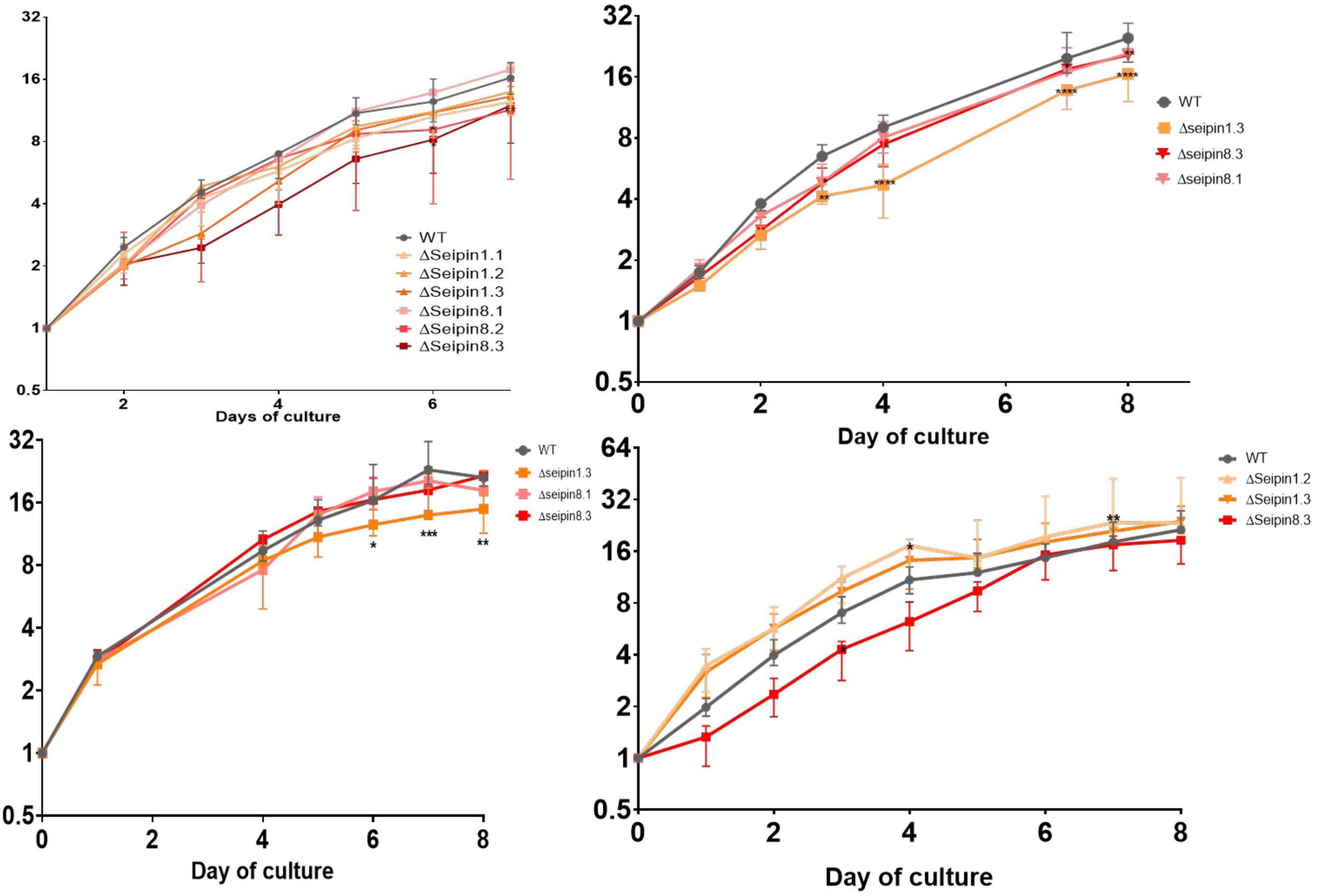


**Supplementary Figure S9: Growth curves of PtSeipin KO mutants**

Growth curves of WT (grey) and PtSeipin KO (orange and red shades) obtained from 4 independent experiments are shown. Cell concentrations (in million cells.mL$^{-1}$) were calculated based on absorbance at 730 nm, which was measured during 8 days. Median, min and max of triplicates are shown. Statistically significant differences between KO and WT were evaluated by 2-way ANOVA with Dunett's multiple comparison test and are indicated as follows: *: p-value<0.05; **: p-value<0.01; *** p-value <0.001.

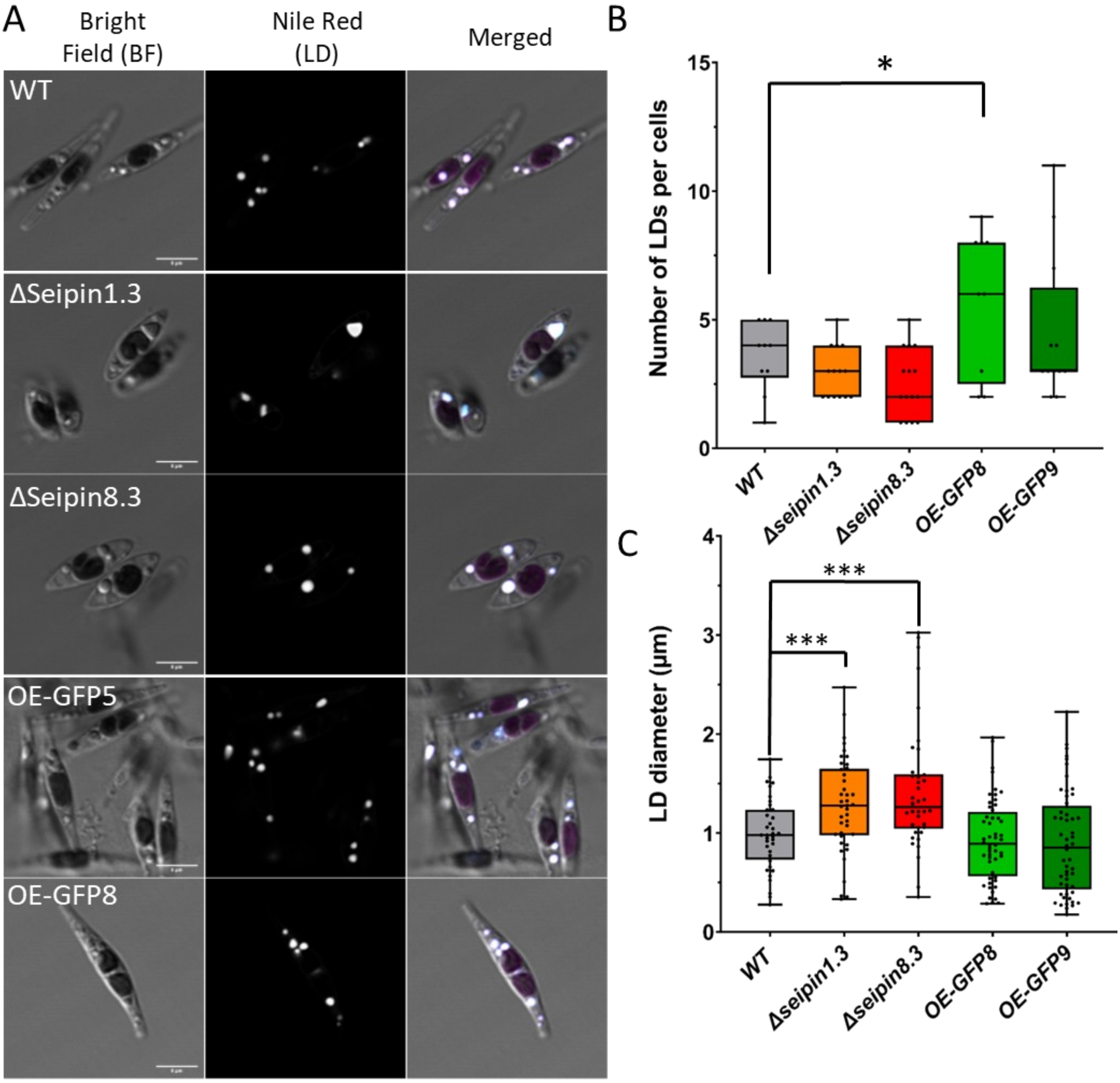


**Supplementary Figure S10: Observation and quantification of LD in PtSeipin mutants after 8 days of culture in control (CT) culture conditions.**

- **A)** Laser Scanning Confocal Microscopy (LSCM) images of WT, PtSeipin KO (ΔSeipin1.3 and ΔSeipin8.3) and PtSeipin-GFP overexpressing lines (OE-GFP5 and OE-GFP8) after staining with Nile Red. From left to right: pseudo-bright field (grey), Plastid and nile red fluorescence (greyscale), nile red fluorescence (greyscale) and overlay of pseudo-bright field with Nile redand chlorophyll signals (respectively grey, cyan and magenta). Scale bar: 5 µm.
- **B)** Quantification of the number of LD per cell in all the cell lines: WT (grey), PtSeipin KO (orange and red) and PtSeipin-GFP overexpressing lines (light and dark green). Results are presented as boxplots and each dot corresponds to an individual cell. *WT*: n=10, *Δseipin1.3*: n=14, *Δseipin8.3*: n=15, *OE-GFP8*: n=9, *OE-GFP9*: n=12. Statistically significant differences between mutants and WT were evaluated using multiple t-tests and are indicated as follows: *: p-value<0.05; **: p-value<0.01; *** p-value<0.001.

**C)** Quantification of the size of LD in all the cell lines: WT (grey), PtSeipin KO (orange and red) and PtSeipin-GFP overexpressing lines (light and dark green). Results are presented as boxplots and each dot corresponds to an individual LD. *WT*: n=36, *Δseipin1.3*: n=41, *Δseipin8.3*: n=38, *OE-GFP8*: n=52, *OE-GFP9*: n=54. Statistically significant differences between mutants and WT were evaluated using multiple t-tests and are indicated as follows: *: p-value<0.05; **: p-value<0.01; *** p-value<0.001.

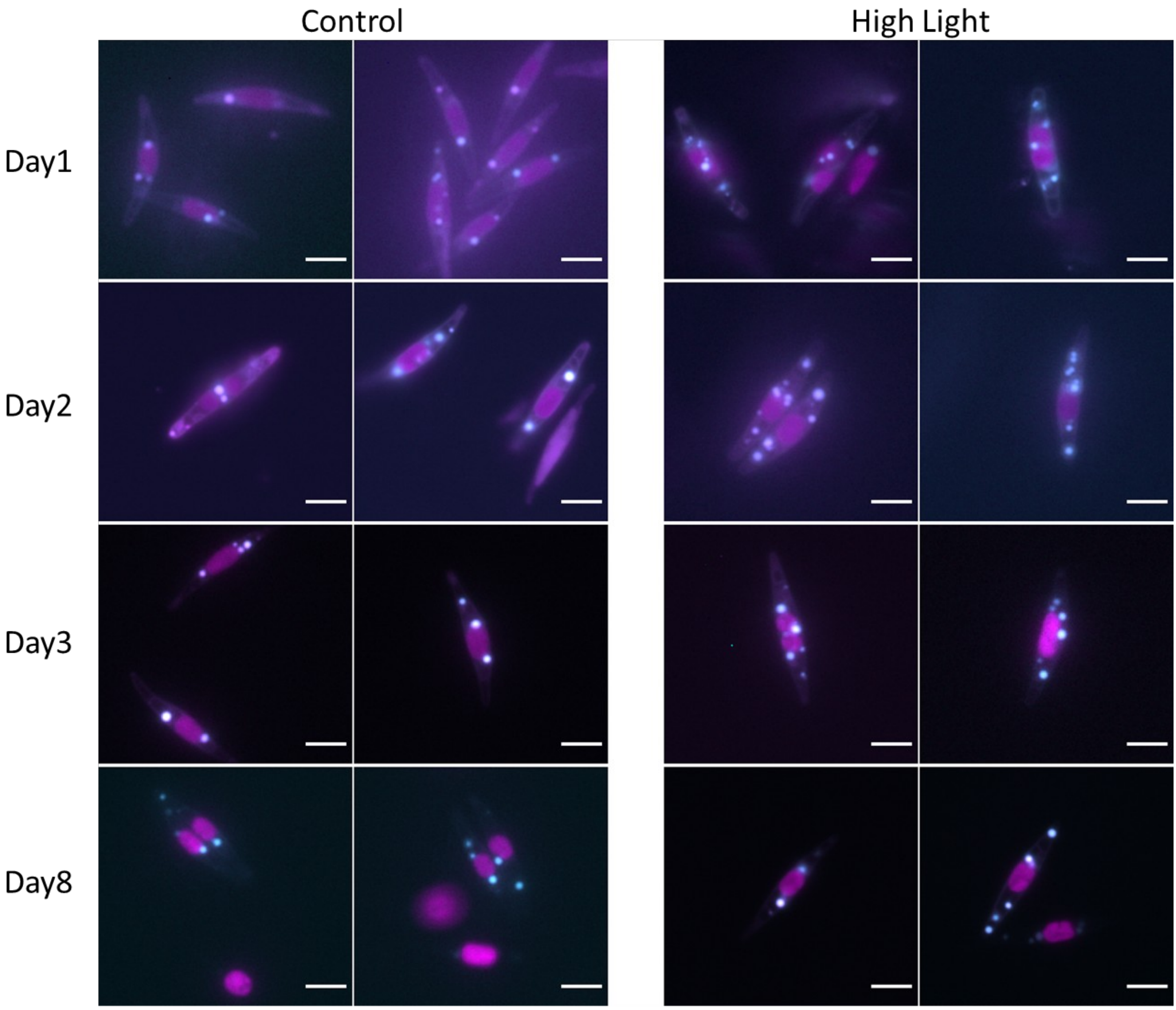


**Supplementary Figure S11: Observation of *Phaeodactylum tricornutum* WT after exposure to h light intensities**

Representative epifluorescence pictures of WT algae after 1, 2, 3 and 8 days of culture in control condition (light intensity 75 µmol.photons.m$^{-2}$.s$^{-1}$, left panel) or in high light (light intensity 200 µmol.photons.m$^{-2}$.s$^{-1}$, right panel). Magenta: chlorophyll autofluorescence; cyan: nile red staining. Scale bar: 5µm.

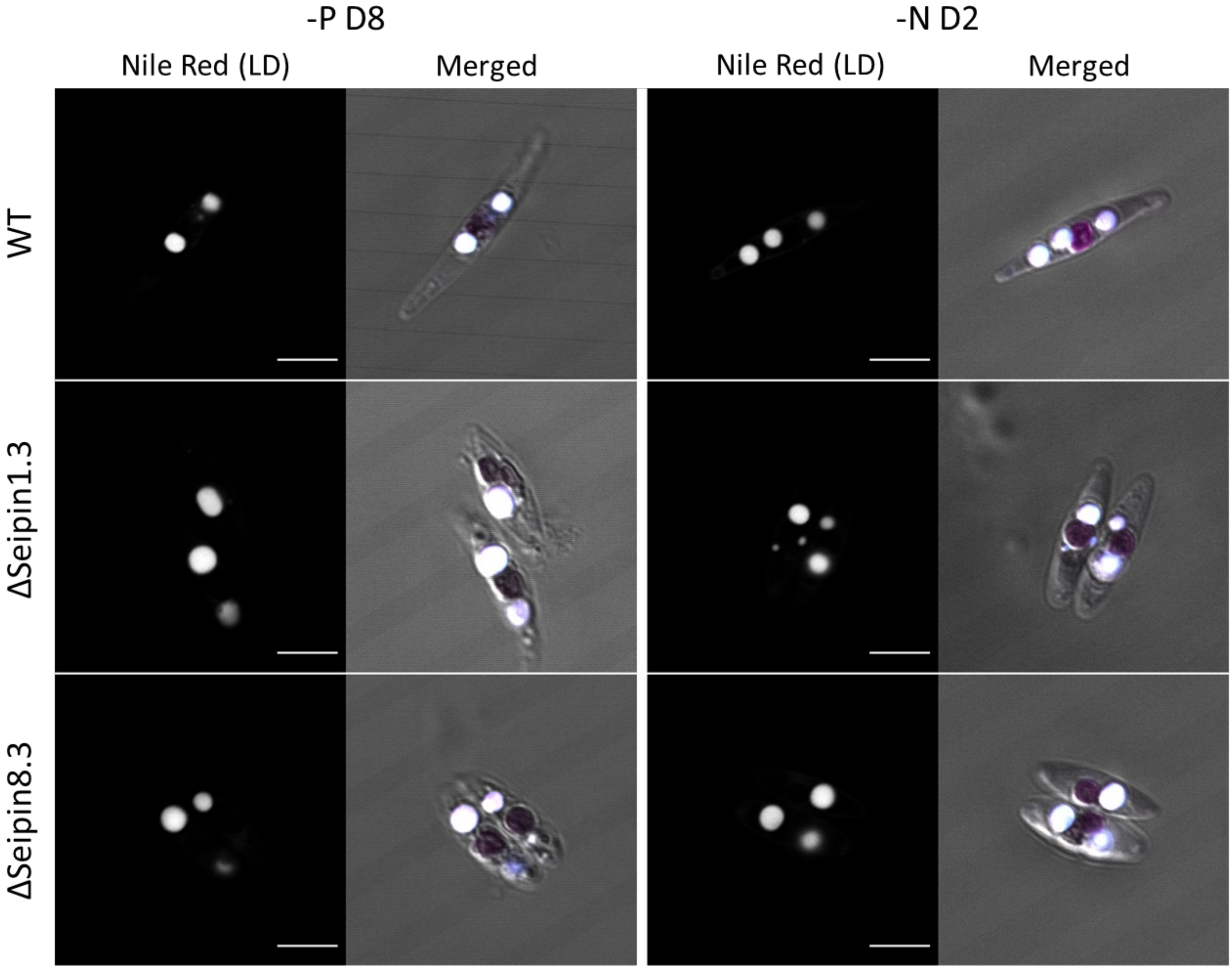


**Supplementary Figure S12**: **Observation of *Phaeodactylum tricornutum* WT and PtSeipin KO following phosphorus and nitrogen starvation**

Representative images of *Phaeodactylum tricornutum* WT and KO for PtSeipin (ΔSeipin1.3 and ΔSeipin8.3) following 8 days of phosphorus deprivation (-P D8) or 2 days of nitrogen deprivation (-N D2). Algae were stained with Nile Red and images were obtained by Laser Scanning Confocal Microscopy (LSCM). Nile Red images are presented in greyscale, merged shows the overlay of pseudo-brightfield (grey), Nile Red (cyan) and chlorophyll (magenta). Scale bar: 5 µm.

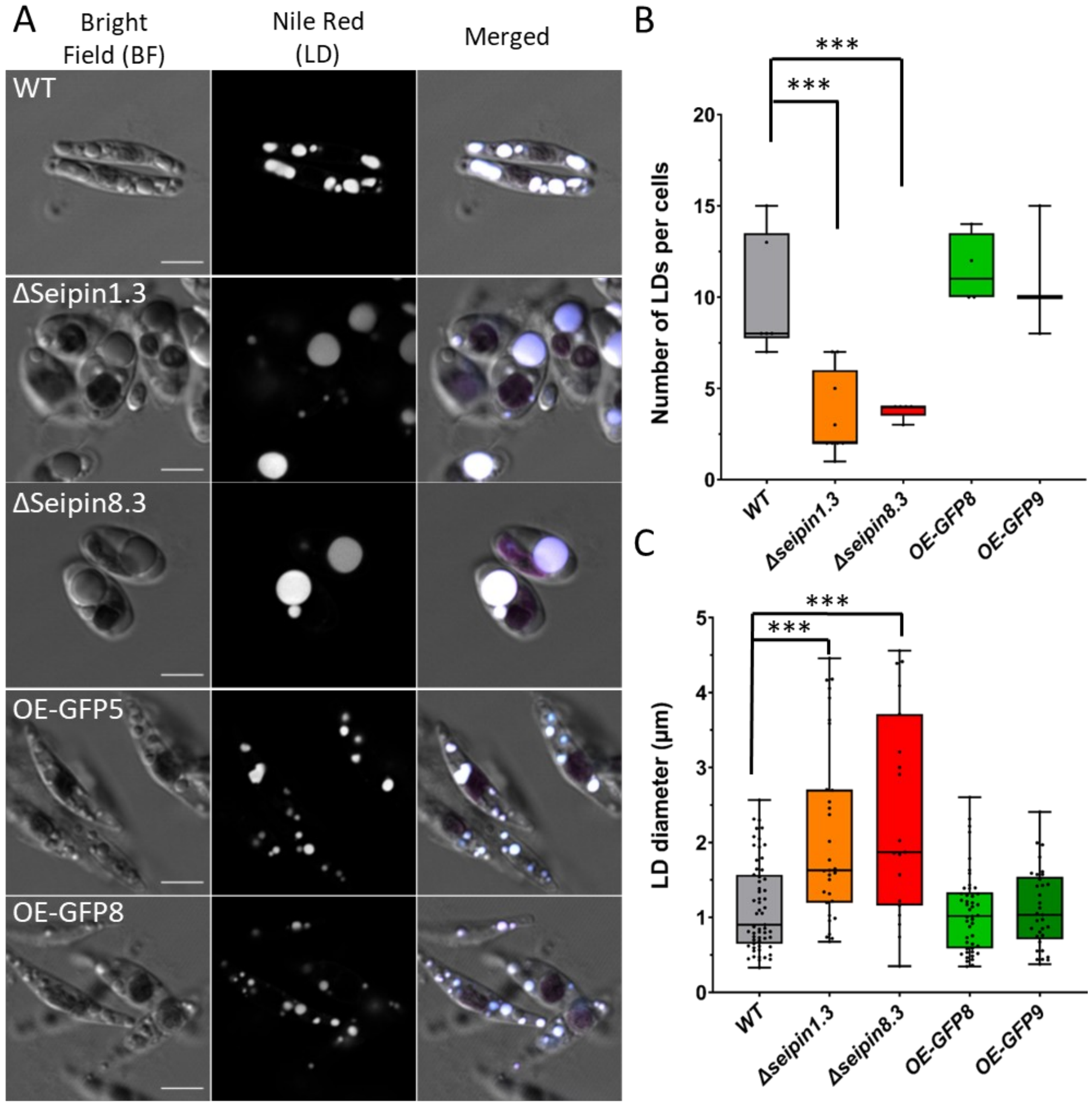


**Supplementary Figure S13: Observation and quantification of LD in PtSeipin mutants after 8 days of culture in high light (HL) culture conditions.**

- **A)** Laser Scanning Confocal Microscopy (LSCM) images of WT, PtSeipin KO (ΔSeipin1.3 and ΔSeipin8.3) and PtSeipin-GFP overexpressing lines (OE-GFP5 and OE-GFP8) after staining with Nile Red. From left to right: pseudo-bright field (grey), Nile Red fluorescence (greyscale) and overlay of pseudo-bright field with Nile Red and chlorophyll signals (respectively grey, cyan and magenta). Scale bar: 5 µm.
- **B)** Quantification of the number of LD per cell in all the cell lines: WT (grey), PtSeipin KO (orange and red) and PtSeipin-GFP overexpressing lines (light and dark green). Results are presented as boxplots and each dot corresponds to an individual cell. *WT*: n=6, *Δseipin1.3*: n=9, *Δseipin8.3*: n=5, *OE-GFP8*: n=10, *OE-GFP9*: n=3. Statistically significant differences between mutants and WT were evaluated using multiple t-tests and are indicated as follows: *: p-value<0.05; **: p-value<0.01; *** p-value<0.001.
- **C)** Quantification of the size of LD in all the cell lines: WT (grey), PtSeipin KO (orange and red) and PtSeipin-GFP overexpressing lines (light and dark green). Results are presented as boxplots and each dot corresponds to an individual LD. *WT*: n=59, *Δseipin1.3*: n=31, *Δseipin8.3*: n=19, *OE-*

*GFP8*: n=81, *OE-GFP9*: n=33. Statistically significant differences between mutants and WT were evaluated using multiple t-tests and are indicated as follows: *: p-value<0.05; **: p-value<0.01; *** p-value<0.001.

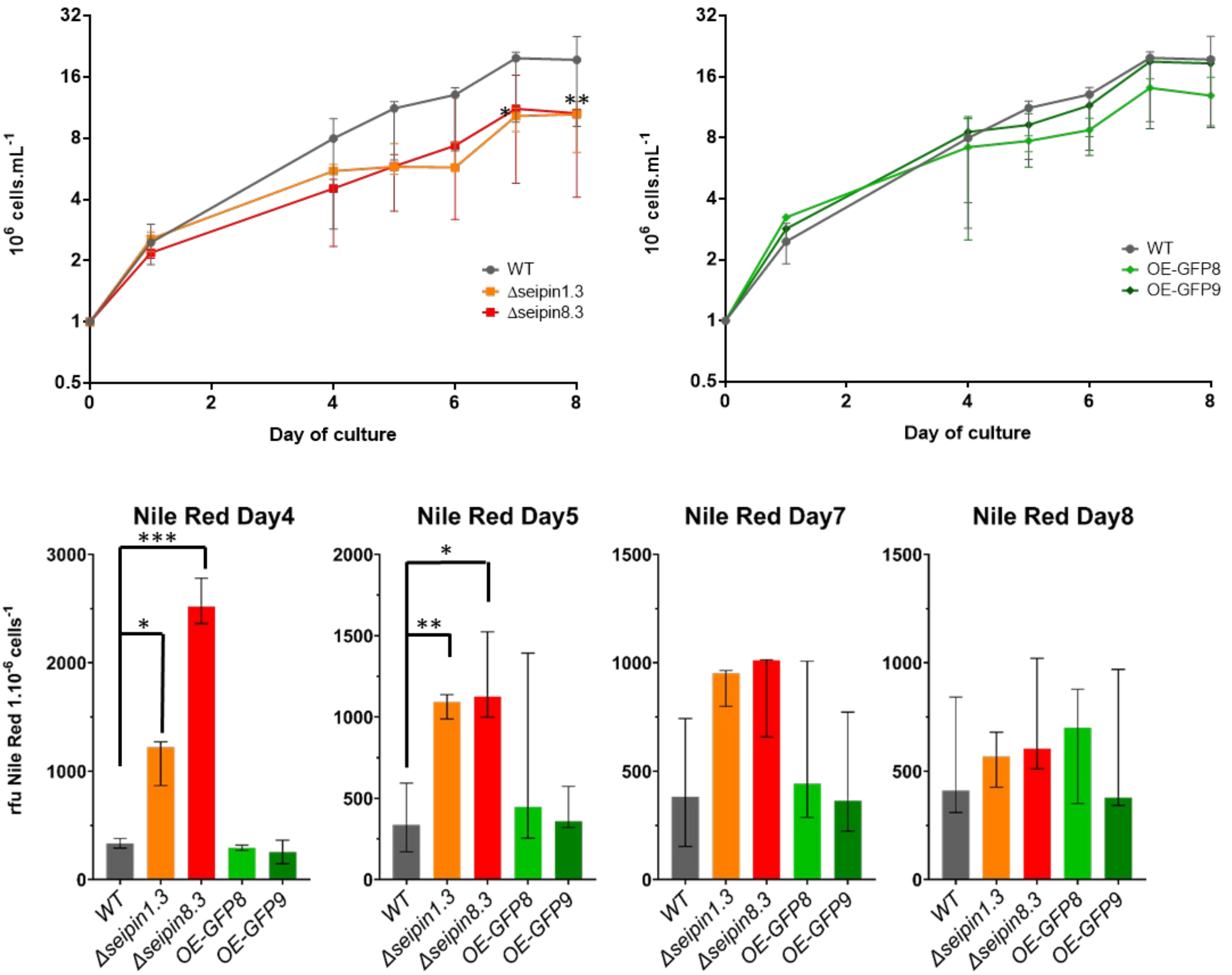


**Supplementary Figure S14: Cell physiology parameters in PtSeipin mutants under control culture conditions**

**A)** Growth curves of WT (grey), PtSeipin KO (orange and red, left panel) and PtSeipin-GFP overexpressing lines (light and dark green, right panel). Cell concentrations (expressed as log2(million cells.mL$^{-1}$)) were calculated based on absorbance at 730 nm, which was measured during 8 days. Median, min and max of triplicates are shown. Statistically significant differences between mutants and WT were evaluated by 2-way ANOVA with Dunett's multiple comparison test and are indicated as follows: *: p-value<0.05; **: p-value<0.01; *** p-value <0.001.

**B)** Neutral lipids accumulation during the stationary phase was evaluated by fluorescence intensity measures following Nile Red staining. WT is in grey, PtSeipin KO lines in orange and red and PtSeipin-GFP overexpressing lines in light and dark green. Statistically significant differences between mutants and WT were evaluated using multiple t-tests and are indicated as follows: *: p-value<0.05; **: p-value<0.01; *** p-value<0.001.

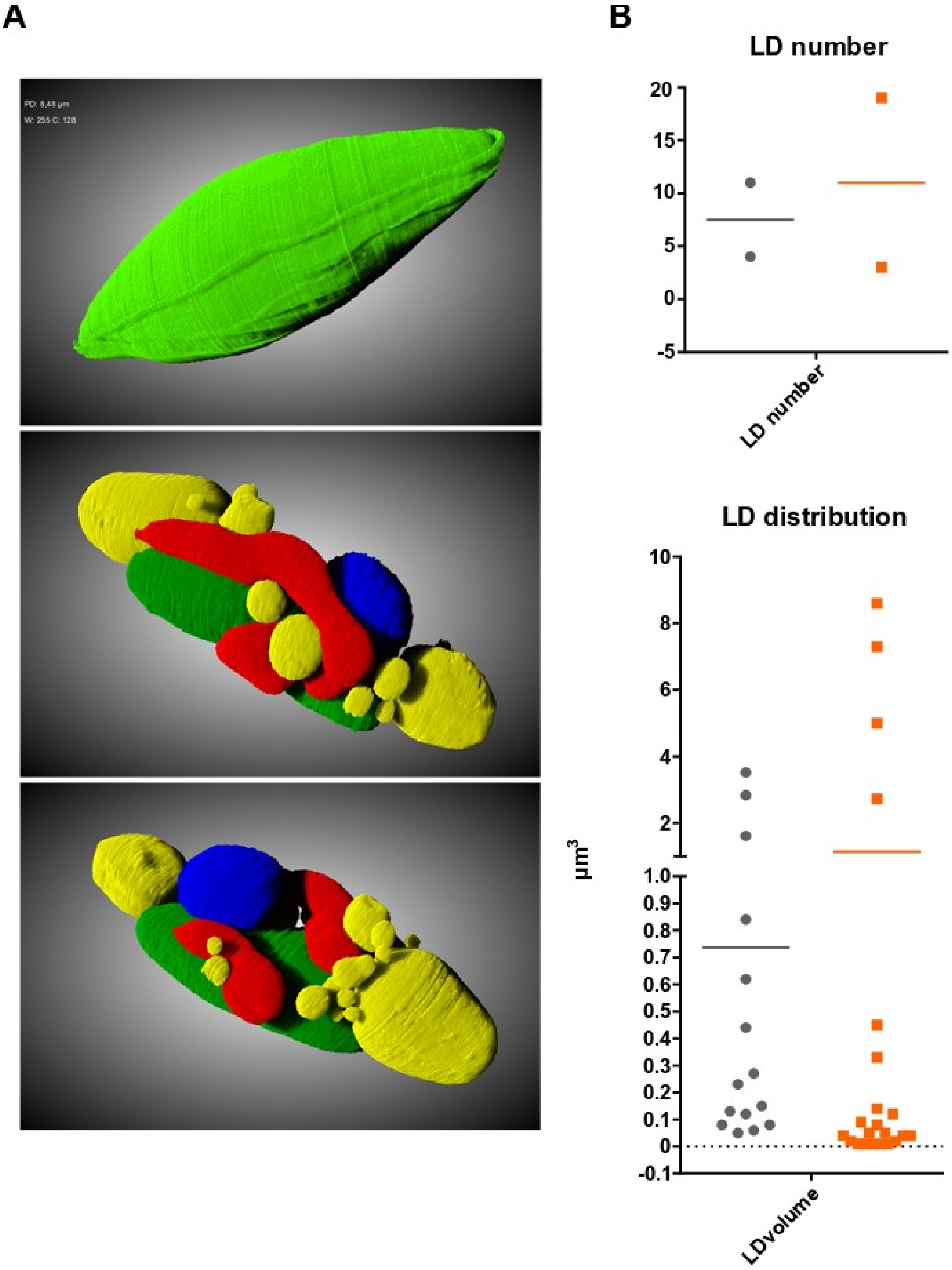


**Supplementary Figure S15: LD numbers and distributions in cells segmented after focused ion beam scanning electron microsopy (FIB-SEM) imaging**

A. Segmentation of PtSeipin KO cell2. Upper panel shows the segmentation of the total volume while the two lower panels show two 3D views of the segmentation of the different organelles. Yellow: LD; red: mitochondria; blue: nucleus; green: plastid
B. Measures of the LD numbers and distribution of LD volumes (in $\mu m^3$) in WT and PtSeipin KO cells. Upper graph: each dot represents the number of LD within one cell; lower graph: each dot represents a LD.

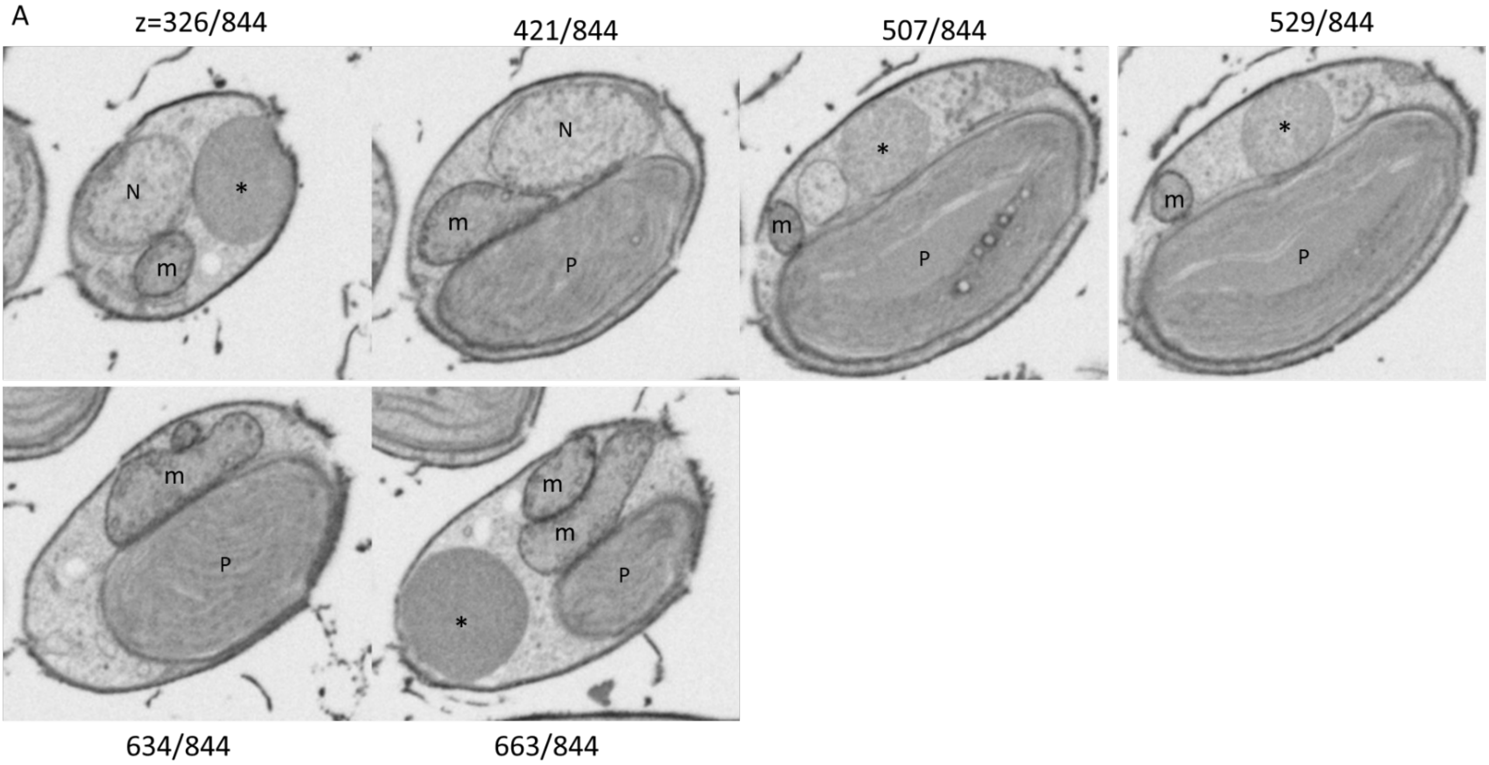
A
z=326/844
421/844
507/844
529/844
N
*
m
N
m
P
m
*
P
m
*
P
m
P
m
m
P
*
634/844
663/844

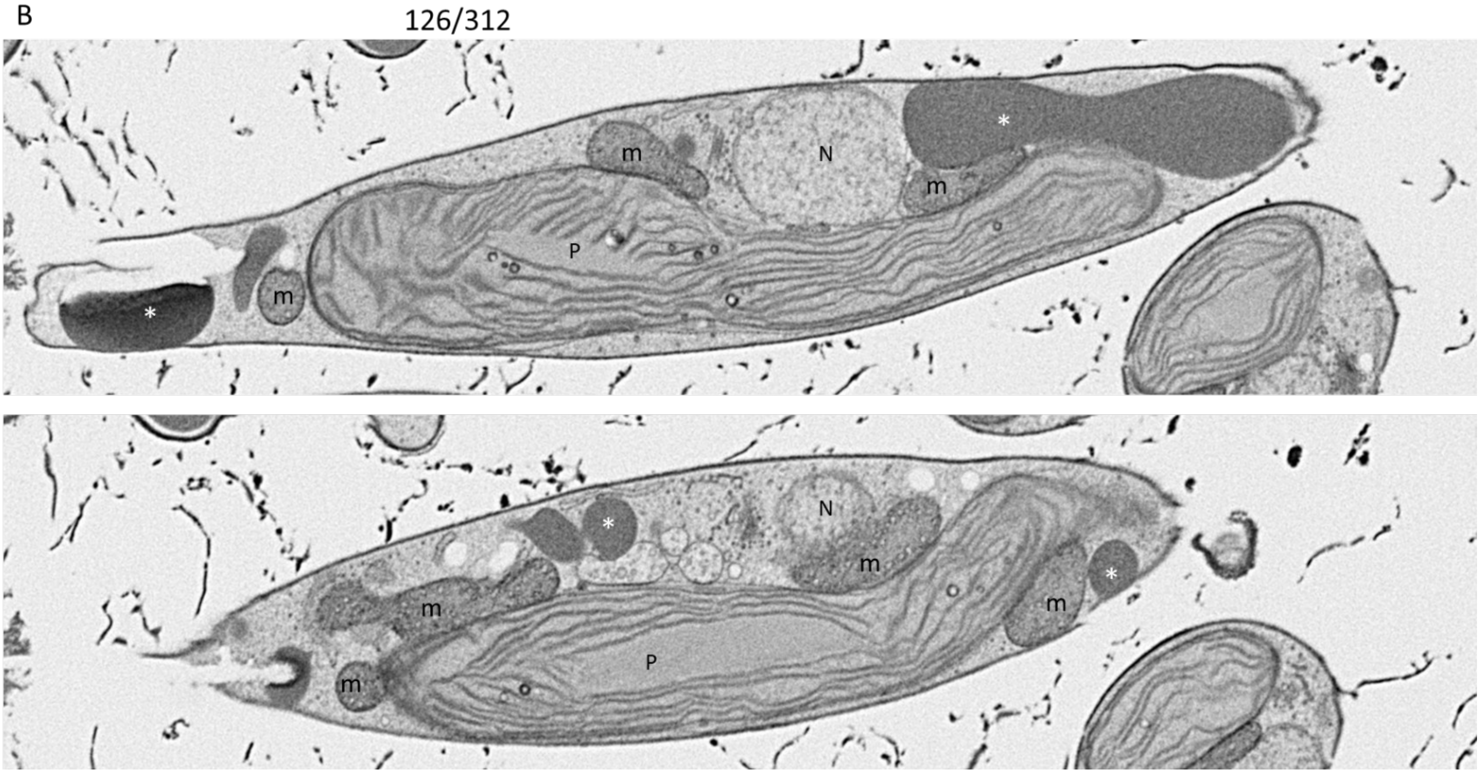
B
126/312
*
m
N
m
P
*
m
*
N
m
m
m
*
P
m

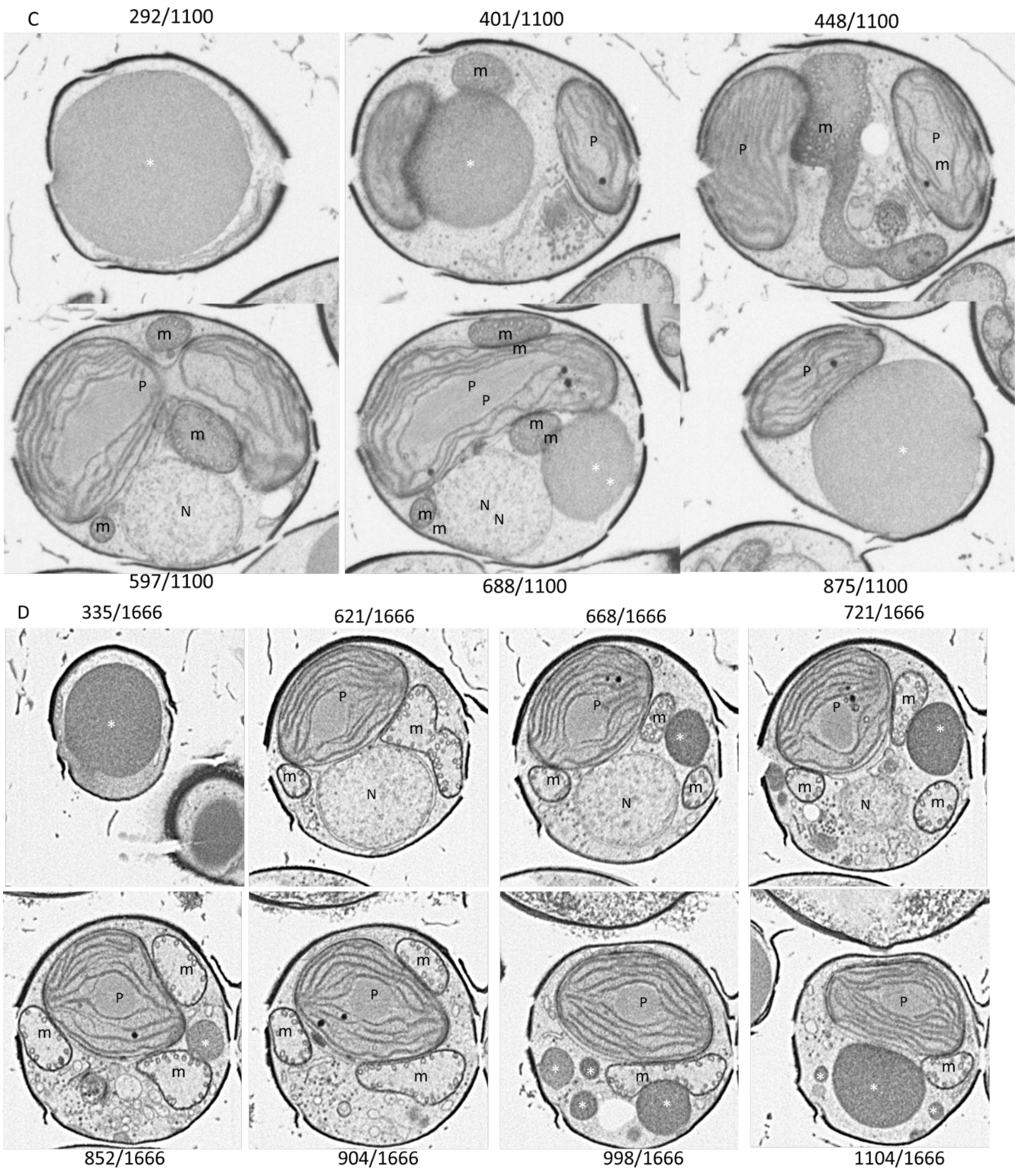


**Supplementary Figure S16**: Images extracted from the FIB-SEM stacks

A. WT cell 1 (*cf.* supplementary movies 1 and 5)
B. WT cell 2 (*cf.* supplementary movies 2 and 6)
C. PtSeipin KO cell 1 (*cf.* supplementary movies 3 and 7)
D. PtSeipin KO cell 2 ( (*cf.* supplementary movies 4 and 8)

For each image, its position in the corresponding stack is indicated. *: LD; N: nucleus: m: mitochondrion; P: plastid.

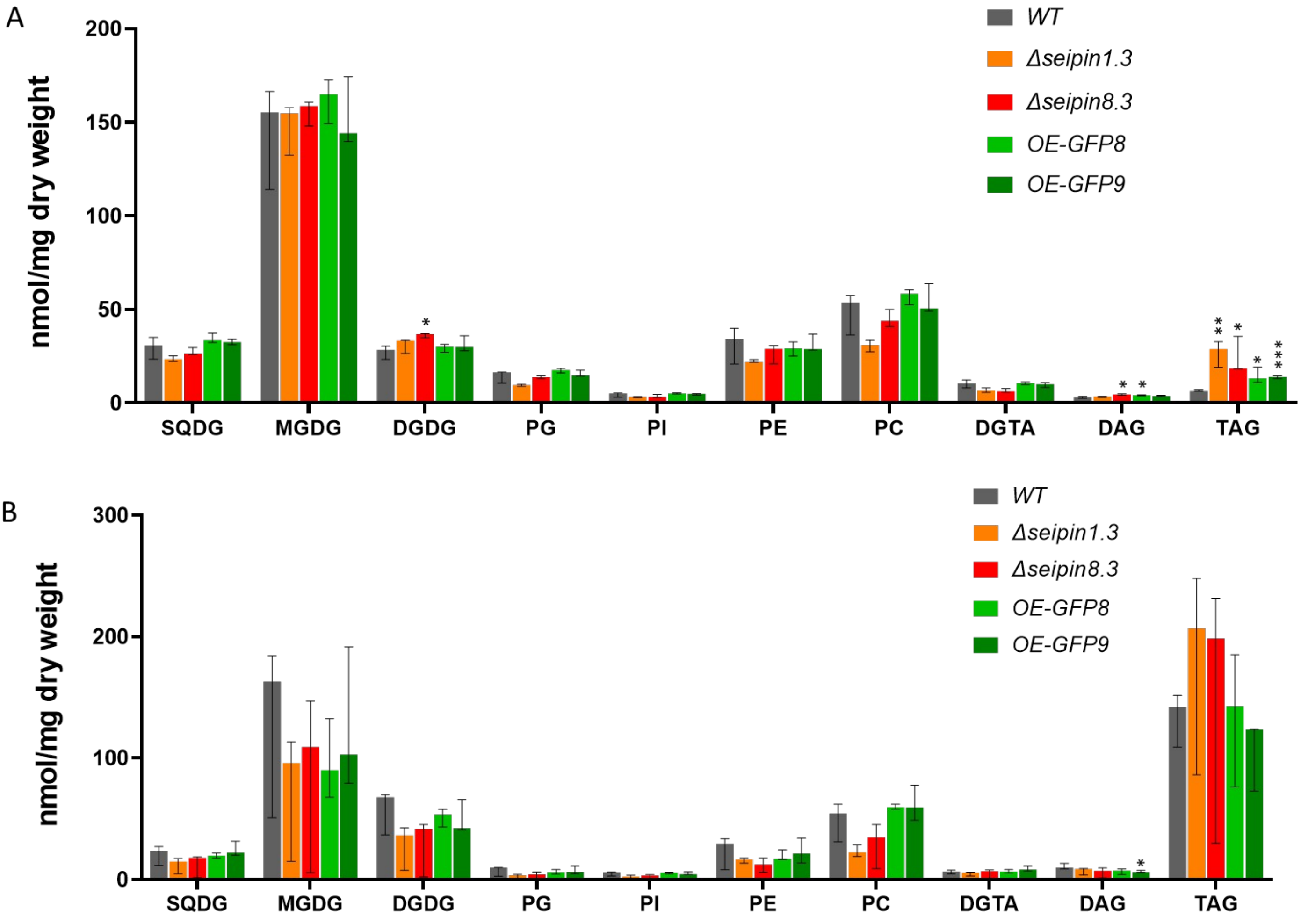


**Supplementary Figure S17: Glycerolipid profiles of WT and PtSeipin mutants after 8 days of culture in control condition (A) and high light condition (B).**

Glycerolipid classes were quantified following liquid chromatography and tandem mass spectrometry (LC-MS/MS) as described in the material and methods, and are presented in nmol/mg of dry weight. Median, min and max values of biological triplicates are shown. SQDG: sulfoquinovosyl-diacylglycerol; MGDG: monogalactosyl-diacylglycerol; DGDG: digalactosyl-diacylglycerol; PG: phosphatidylglycerol; PI: phosphatidylinositol, PE: phosphatidylethanolamine, PC: phosphatidylcholine, DGTA: 1(3),2-Diacylglyceryl-3(1)-O-2′-(hydroxymethyl)(N,N,N,-trimethyl)-β-alanine; DAG: diacylglycerol; TAG: triacylglycerol. Statistically significant differences between mutants and WT were evaluated using multiple t-tests and are indicated as follows: *: p-value<0.05; **: p-value<0.01; *** p-value<0.001.

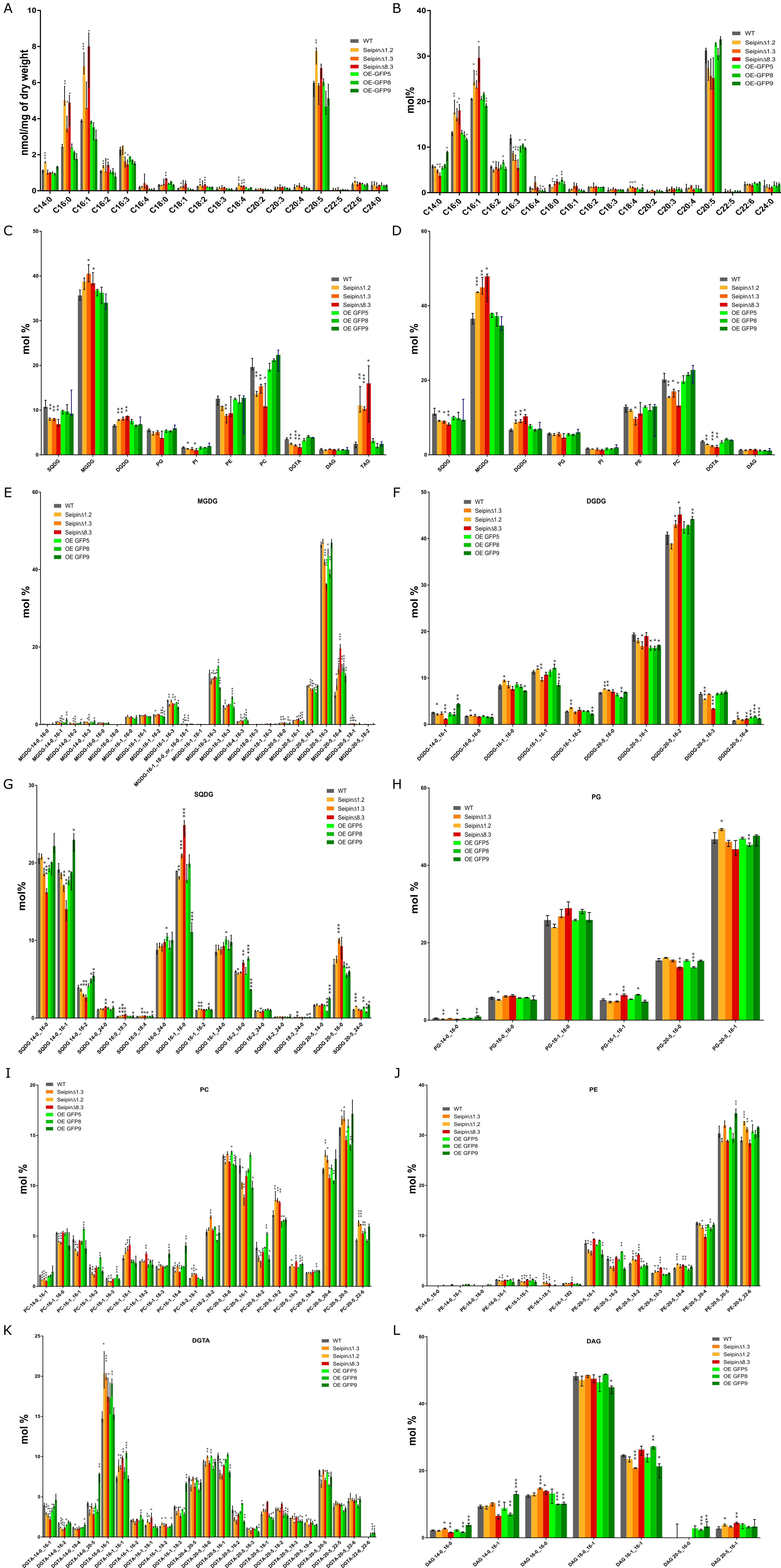

A
nmol/mg of dry weight
WT
SeipinΔ1.2
SeipinΔ1.3
SeipinΔ8.3
OE-GFP5
OE-GFP8
OE-GFP9
C14:0 C16:0 C16:1 C16:2 C16:3 C16:4 C18:0 C18:1 C18:2 C18:3 C18:4 C20:2 C20:3 C20:4 C20:5 C22:5 C22:6 C24:0
B
mol%
C
mol %
SQDG MGDG DGDG PG PI PE PC DGTA DAG TAG
OE GFP5
OE GFP8
OE GFP9
D
E
MGDG
F
DGDG
G
SQDG
H
PG
I
PC
J
PE
K
DGTA
L
DAG

**Supplementary Figure S18: Lipids profiles of WT and PtSeipin mutants after 4 days of culture in control condition.**

- **A.** Fatty acids profiles obtained by GC-FID in nmol/mg of dry weight.
- **B.** Fatty acids profiles obtained by GC-FID expressed in mol% of all fatty acids.
- **C.** Glycerolipids profile obtained by LC-MS/MS expressed in mol % of all glycerolipids
- **D.** Glycerolipids profile obtained by LC-MS/MS expressed in mol % of all glycerolipids after exclusion of TAG
- **E.** MGDG profiles obtained by LC-MS/MS expressed in mol %
- **F.** DGDG profiles obtained by LC-MS/MS expressed in mol %
- **G.** SQDG profiles obtained by LC-MS/MS expressed in mol %
- **H.** PG profiles obtained by LC-MS/MS expressed in mol %
- **I.** PC profiles obtained by LC-MS/MS expressed in mol %
- **J.** PE profiles obtained by LC-MS/MS expressed in mol %
- **K.** DGTA profiles obtained by LC-MS/MS expressed in mol %
- **L.** DAG profiles obtained by LC-MS/MS expressed in mol %

For each bar, median, min and max values of biological triplicates are shown. Statistically significant differences between mutants and WT were evaluated using multiple t-tests and are indicated as follows: *: p-value<0.05; **: p-value<0.01; *** p-value<0.001. SQDG: sulfoquinovosyl-diacylglycerol; MGDG: monogalactosyl-diacylglycerol; DGDG: digalactosyl-diacylglycerol; PG: phosphatidylglycerol; PI: phosphatidylinositol, PE: phosphatidylethanolamine, PC: phosphatidylcholine, DGTA: 1(3),2-Diacylglyceryl-3(1)-O-2′-(hydroxymethyl)(N,N,N,-trimethyl)-β-alanine; DAG: diacylglycerol; TAG: triacylglycerol.

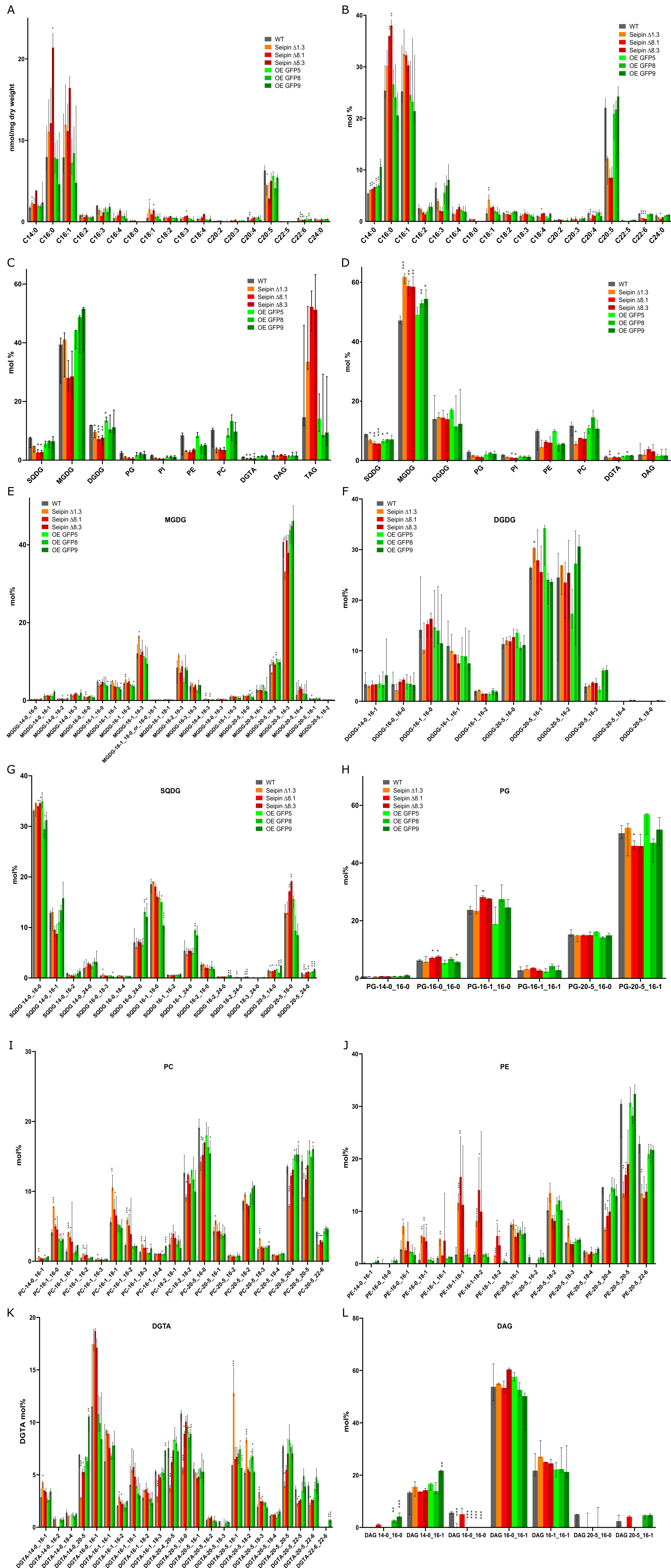
A
nmol/mg dry weight
WT
Seipin Δ1.3
Seipin Δ8.1
Seipin Δ8.3
OE GFP5
OE GFP8
OE GFP9
C14:0 C16:0 C16:1 C16:2 C16:3 C16:4 C18:0 C18:1 C18:2 C18:3 C18:4 C20:2 C20:3 C20:4 C20:5 C22:5 C22:6 C24:0
B
mol %
C
mol %
SQDG MGDG DGDG PG PI PE PC DGTA DAG TAG
D
mol %
SQDG MGDG DGDG PG PI PE PC DGTA DAG
E
MGDG
mol%
F
DGDG
mol%
DGDG-14-0_16-1 DGDG-16-0_16-0 DGDG-16-1_16-0 DGDG-16-1_16-1 DGDG-16-1_16-2 DGDG-20-5_16-0 DGDG-20-5_16-1 DGDG-20-5_16-2 DGDG-20-5_16-3 DGDG-20-5_16-4 DGDG-20-5_18-0
G
SQDG
mol%
H
PG
mol%
PG-14-0_16-0 PG-16-0_16-0 PG-16-1_16-0 PG-16-1_16-1 PG-20-5_16-0 PG-20-5_16-1
I
PC
mol%
J
PE
mol%
K
DGTA
DGTA mol%
L
DAG
mol%
DAG 14-0_16-0 DAG 14-0_16-1 DAG 16-0_16-0 DAG 16-0_16-1 DAG 16-1_16-1 DAG 20-5_16-0 DAG 20-5_16-1

**Supplementary Figure S19: Lipids profiles of WT and PtSeipin mutants after 4 days of culture in high light condition.**

- **A.** Fatty acids profiles obtained by GC-FID in nmol/mg of dry weight.
- **B.** Fatty acids profiles obtained by GC-FID expressed in mol% of all fatty acids.
- **C.** Glycerolipids profile obtained by LC-MS/MS expressed in mol % of all glycerolipids
- **D.** Glycerolipids profile obtained by LC-MS/MS expressed in mol % of all glycerolipids after exclusion of TAG
- **E.** MGDG profiles obtained by LC-MS/MS expressed in mol %
- **F.** DGDG profiles obtained by LC-MS/MS expressed in mol %
- **G.** SQDG profiles obtained by LC-MS/MS expressed in mol %
- **H.** PG profiles obtained by LC-MS/MS expressed in mol %
- **I.** PC profiles obtained by LC-MS/MS expressed in mol %
- **J.** PE profiles obtained by LC-MS/MS expressed in mol %
- **K.** DGTA profiles obtained by LC-MS/MS expressed in mol %
- **L.** DAG profiles obtained by LC-MS/MS expressed in mol %

For each bar, median, min and max values of biological triplicates are shown. Statistically significant differences between mutants and WT were evaluated using multiple t-tests and are indicated as follows: *: p-value<0.05; **: p-value<0.01; *** p-value<0.001. SQDG: sulfoquinovosyl-diacylglycerol; MGDG: monogalactosyl-diacylglycerol; DGDG: digalactosyl-diacylglycerol; PG: phosphatidylglycerol; PI: phosphatidylinositol, PE: phosphatidylethanolamine, PC: phosphatidylcholine, DGTA: 1(3),2-Diacylglyceryl-3(1)-O-2′-(hydroxymethyl)(N,N,N,-trimethyl)-β-alanine; DAG: diacylglycerol; TAG: triacylglycerol.

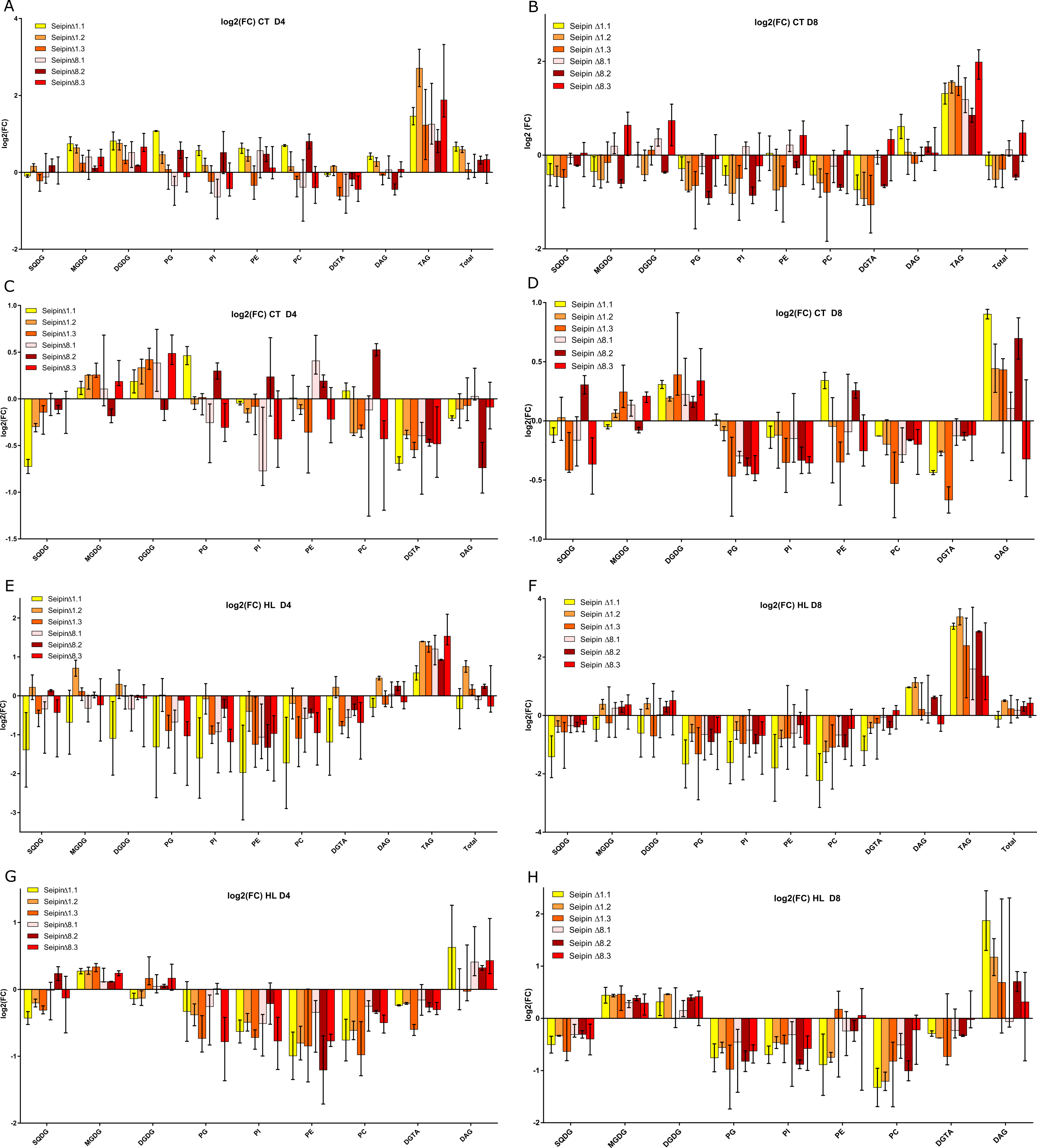
A
log2(FC) CT D4
Seipin∆1.1
Seipin∆1.2
Seipin∆1.3
Seipin∆8.1
Seipin∆8.2
Seipin∆8.3
log2(FC)
4
2
0
-2
SQDG
MGDG
DGDG
PG
PI
PE
PC
DGTA
DAG
TAG
Total
B
log2(FC) CT D8
Seipin ∆1.1
Seipin ∆1.2
Seipin ∆1.3
Seipin ∆8.1
Seipin ∆8.2
Seipin ∆8.3
log2 (FC)
2
0
-2
SQDG
MGDG
DGDG
PG
PI
PE
PC
DGTA
DAG
TAG
Total
C
log2(FC) CT D4
Seipin∆1.1
Seipin∆1.2
Seipin∆1.3
Seipin∆8.1
Seipin∆8.2
Seipin∆8.3
log2(FC)
1.0
0.5
0.0
-0.5
-1.0
-1.5
SQDG
MGDG
DGDG
PG
PI
PE
PC
DGTA
DAG
D
log2(FC) CT D8
Seipin ∆1.1
Seipin ∆1.2
Seipin ∆1.3
Seipin ∆8.1
Seipin ∆8.2
Seipin ∆8.3
log2(FC)
1.0
0.5
0.0
-0.5
-1.0
SQDG
MGDG
DGDG
PG
PI
PE
PC
DGTA
DAG
E
log2(FC) HL D4
Seipin∆1.1
Seipin∆1.2
Seipin∆1.3
Seipin∆8.1
Seipin∆8.2
Seipin∆8.3
log2(FC)
2
1
0
-1
-2
-3
SQDG
MGDG
DGDG
PG
PI
PE
PC
DGTA
DAG
TAG
Total
F
log2(FC) HL D8
Seipin ∆1.1
Seipin ∆1.2
Seipin ∆1.3
Seipin ∆8.1
Seipin ∆8.2
Seipin ∆8.3
log2(FC)
4
2
0
-2
-4
SQDG
MGDG
DGDG
PG
PI
PE
PC
DGTA
DAG
TAG
Total
G
log2(FC) HL D4
Seipin∆1.1
Seipin∆1.2
Seipin∆1.3
Seipin∆8.1
Seipin∆8.2
Seipin∆8.3
log2(FC)
1
0
-1
-2
SQDG
MGDG
DGDG
PG
PI
PE
PC
DGTA
DAG
H
log2(FC) HL D8
Seipin ∆1.1
Seipin ∆1.2
Seipin ∆1.3
Seipin ∆8.1
Seipin ∆8.2
Seipin ∆8.3
log2(FC)
2
1
0
-1
-2
SQDG
MGDG
DGDG
PG
PI
PE
PC
DGTA
DAG

**Supplementary Figure S20: Changes in glycerolipids distribution across all KO lines and all experiments.**

Glycerolipids were quantified by LC-MS/MS as described in the material and methods and expressed in nmol/mg of dry weight. For each experiment, the log2 of the fold change of each lipid in each KO sample was calculated relatively to the median of the same lipid in the WT in the same experiment (A, B, E and F). Alternatively, glycerolipids quantities were calculated as % of all glycerolipids excluding TAG. Log2 of fold changes of those % were calculated as described above (C, D, G and H).

**A and C.** Glycerolipids at day 4 (D4) in control (CT) condition
**B and D.** Glycerolipids at day 4 (D4) in high light (HL) condition
**E and G.** Glycerolipids at day 8 (D8) in control (CT) condition
**F and H.** Glycerolipids at day 8 (D8) in high light (HL) condition

Results are shown as median +-interquartile range. SQDG: sulfoquinovosyl-diacylglycerol; MGDG: monogalactosyl-diacylglycerol; DGDG: digalactosyl-diacylglycerol; PG: phosphatidylglycerol; PI: phosphatidylinositol, PE: phosphatidylethanolamine, PC: phosphatidylcholine, DGTA: 1(3),2-Diacylglyceryl-3(1)-O-2′-(hydroxymethyl)(N,N,N,-trimethyl)-β-alanine; DAG: diacylglycerol; TAG: triacylglycerol.

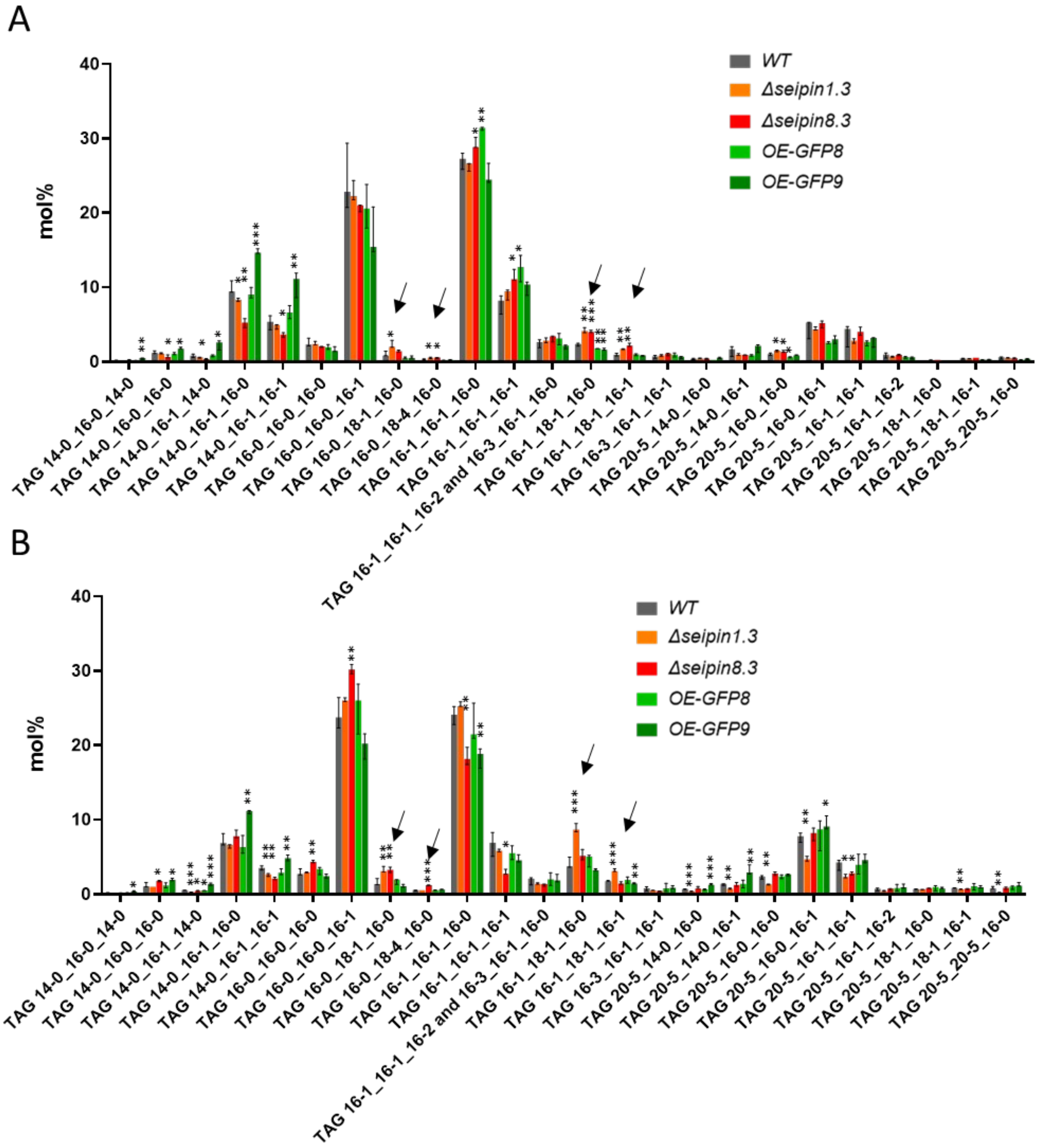


**Supplementary Figure S21: Triacylglycerol (TAG) profiles of WT and PtSeipin mutants after 8 days of culture in control condition (A) and high light condition (B).**

TAG species were quantified following liquid chromatography and tandem mass spectrometry (LC-MS/MS) as described in the material and methods, and are presented in mol%. Median, min and max values of biological triplicates are shown. Statistically significant differences between mutants and WT were evaluated using multiple t-tests and are indicated as follows: *: p-value<0.05; **: p-value<0.01; *** p-value<0.001.

**Supplementary Material**

The full-length coding sequence of *P. tricornutum Seipin* (*PtSeipin*) was amplified by PCR using *PtSeipin-F* sense (CCGGAATTCATGGACCGACGCGTTTCCCG) and *PtSeipin_R* antisense (CGCGGATCCAATAGCTTCGGCTGGAGTAACCG) primers with *P. tricornutum* cDNA as template. The PCR products were cloned into pClone007 Vector for DNA sequencing. Sequence-confirmed *PtSeipin* gene was excised from the vector with EcoRI and BamHI and inserted into the pPhaT1-eGFP vector between the EcoRI and KpnI sites (Zhang and Hu, 2014). The resulting pPha-*PtSeipin*-eGFP vector was linearized and introduced in *P. tricornutum* cells by electroporation as previously described (Zhang and Hu 2014). Fluorescence of eGFP and chlorophyll autofluorescence were excited at 488 nm, and were detected with two photomultiplier tubes at 500–520 nm and 625–720 nm respectively using a Leica TCS SP8 laser scanning confocal microscope.
